# Biophysical Mechanism of Allosteric Regulation of Actin Capping Protein

**DOI:** 10.1101/2023.08.16.553570

**Authors:** Olivia L. Mooren, Melissa D. Stuchell-Brereton, Patrick McConnell, Chenbo Yan, Emily M. Wilkerson, Dennis Goldfarb, John A. Cooper, David Sept, Andrea Soranno

**Author notes:** Equal contributions (Olivia L. Mooren, Melissa D. Stuchell-Brereton). Equal contributions (John A. Cooper, David Sept, Andrea Soranno). Correspondence after publication: Andrea Soranno; David Sept; John A. Cooper. Contact information before publication: John A Cooper, CB 8231, 660 S. Euclid Ave. St Louis, MO 63110.

## Abstract

Actin capping protein (CP) can be regulated by steric and allosteric mechanisms. The molecular mechanism of the allosteric regulation at a biophysical level includes linkage between the binding sites for three ligands: F-actin, Capping-Protein-Interacting (CPI) motifs, and V-1/myotrophin, based on biochemical functional studies and solvent accessibility experiments. Here, we investigated the mechanism of allosteric regulation at the atomic level using single-molecule Förster resonance energy transfer (FRET) and molecular dynamics (MD) to assess the conformational and structural dynamics of CP in response to linked-binding site ligands. In the absence of ligand, both single-molecule FRET and MD revealed two distinct conformations of CP in solution; previous crystallographic studies revealed only one. CPI-motif peptide association induced conformational changes within CP that propagate in one direction, while V-1 association induced conformational changes in the opposite direction. Comparing CPI-motif peptides from different proteins, we identified variations in CP conformations and dynamics that are specific to each CPI motif. MD simulations for CP alone and in complex with a CPI motif and V-1 reveal atomistic details of the conformational changes. Analysis of the interaction of CP with wildtype (wt) and chimeric CPI-motif peptides using single-molecule FRET, isothermal calorimetry (ITC) and MD simulation indicated that conformational and affinity differences are intrinsic to the C-terminal portion of the CPI-motif. We conclude that allosteric regulation of CP involves changes in conformation that disseminate across the protein to link distinct binding-site functions. Our results provide novel insights into the biophysical mechanism of the allosteric regulation of CP.

## Introduction

Actin assembly and disassembly are important for directing the shape and movement of cells and tissues during normal development and physiological function as well as for aiding in the motility of some pathogens upon infection ^1^. Actin filaments grow and shrink by gaining and losing monomeric subunits from their two ends – barbed and pointed. Regulation of filament polymerization and depolymerization via actin capping protein (CP) and its binding partners is essential to provide the force and energy for movements of many cellular membranes, including vesicles and the plasma membrane.

CP is an α / β heterodimer that adopts a mushroom-like shape. The mushroom cap is comprised of interlaced β sheets and α helices. The mushroom stalk protrudes perpendicularly from the bottom of the cap surface and is composed of α helices (Figure 1). The top surface of the mushroom cap binds to filament barbed ends, preventing association and dissociation of actin subunits ^2^. A number of biomolecules bind directly to CP and regulate the interaction of CP with barbed ends. Those molecules comprise polyphosphoinositides, the protein V-1 / myotrophin, and a diverse set of proteins with CP-interacting (CPI) motifs, including the CARMIL, CKIP and WASHCAP (FAM21) protein families ^2–4^.

**Figure 1.**
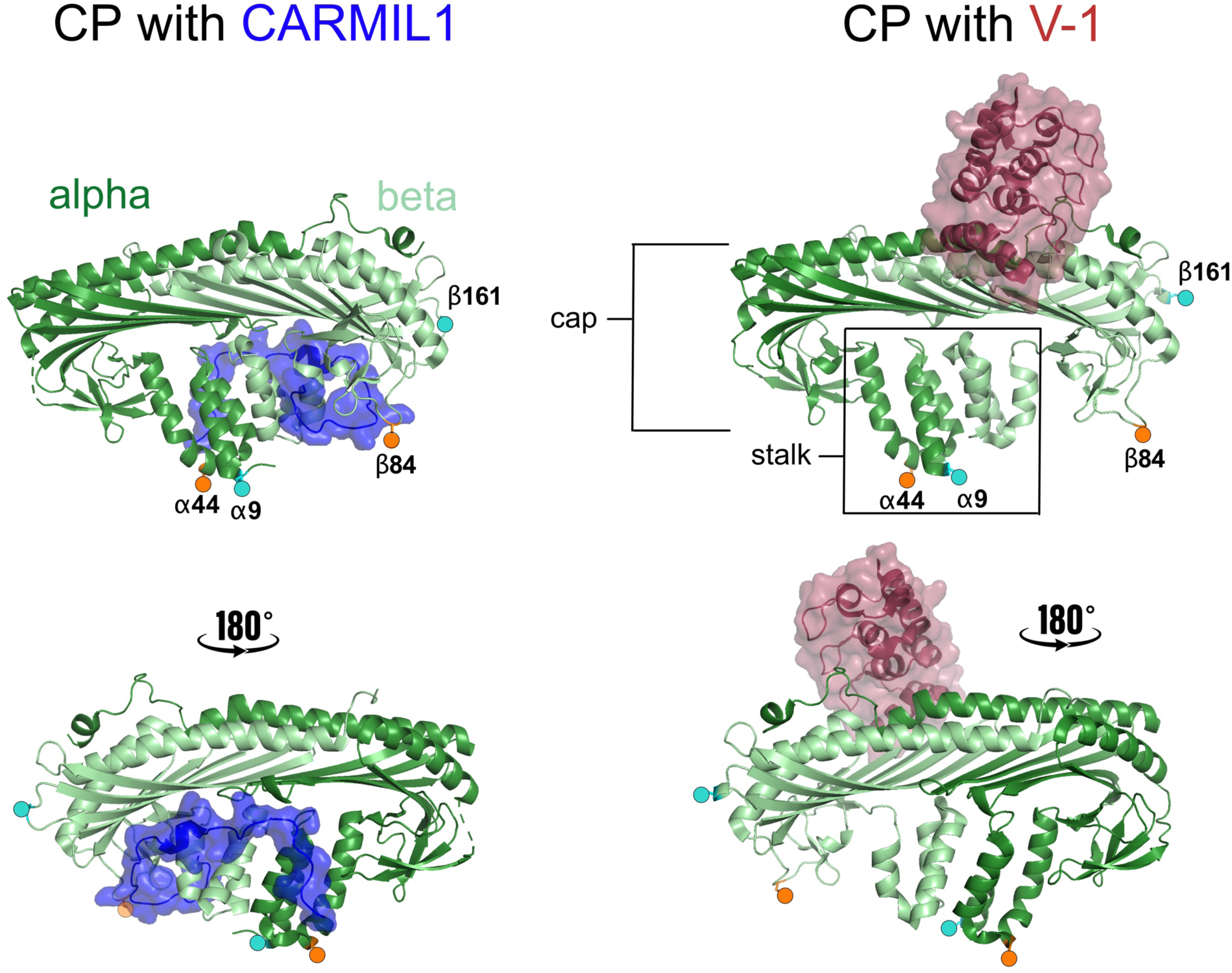
Illustration of dye positions, using previously published structures of CP-ligand complexes (PDB 3LK3 and 3AAA). FRET construct positions are show as teal circles (α9β161) and orange circles (α44β84). CP α subunit shown as dark green ribbon, and CP β subunit as light green ribbon. CARMIL1 CPI motif peptide shown as blue space-filling (modified from PDB 3LK3), and V-1 shown as red space-filling (modified from PDB 3AAA).

In cytoplasm, CP and V-1 are both present at high concentration (micromolar), they each diffuse freely, and they bind tightly to each other (nanomolar affinity) ^5^. In contrast, CPI-motif proteins are present in far smaller amounts, and they are generally targeted to specific membrane locations ^3, 4^. The effect of CPI-motif binding includes weakening the binding affinity of CP for V-1 ^6, 7^. These observations raised the possibility, proposed by Hammer and colleagues ^6^, that CPI-motif proteins activate CP locally at a membrane by promoting dissociation of the V-1 inhibitor.

The binding of CP to V-1, F-actin, and CPI motifs, are linked molecular processes ^7^. V-1 binds directly to the cap region of CP and sterically blocks the site required for binding to actin-filament barbed ends ^5, 6, 8–10^. Structures of co-complexes show that the binding sites for F-actin and V-1 overlap extensively, but not completely ^7^. In contrast, the binding sites for CPI-motif proteins are instead located in the stalk region of CP and are spatially different from those for F-actin and V-1; they bind to the stalk region of CP (Figure 1) ^3, 4, 9, 10^. Therefore, CPI-motif proteins inhibit the binding of CP to filament barbed ends and to V-1 by an allosteric mechanism ^3, 4, 9, 10^.

The physical basis of the allosteric regulation and linkage between these two distinct sites was investigated with hydrogen-deuterium exchange mass spectrometry (HDX-MS) ^11^. HDX-MS revealed that the solvent accessibility of CP was altered at both binding sites when either a CPI-motif peptide or V-1 was added to CP in solution. The findings indicate that interaction with either ligand can induce conformational changes to sites within CP that are distant from the ligand binding surface. Available X-ray crystal structures of co-complexes show only slight differences in the conformations of CP when complexed with a CPI-motif peptide or V-1 ^9, 12, 13^; however, rather large biochemical effects are observed when CP is bound to either ligand ^6, 7, 9^. Together, these findings are consistent with linked changes in the conformation and/or the structural dynamics of the two sites. The atomistic details of the linkage are not clear. In particular, the spatial resolution of HDX-MS is limited to proteolytic peptides, and crystal structures provide atomic resolution, but they do not capture potential dynamics within the different states.

Here, to advance our understanding of the allosteric linkage mechanism that regulates CP function, we sought evidence at the atomic level for changes in the conformation or dynamics of CP in solution, by comparing free CP (Apo-CP) with CP complexed with either a CPI-motif peptide or V-1. We used confocal single-molecule Förster resonance energy transfer (FRET) of molecules in solution, which allows one to quantify conformational changes in the protein alone and in complex with regulators. We complemented the experimental results with molecular dynamics (MD) analyses of the conformation of CP in those settings, to provide an atomistic description of the structural changes identified by single-molecule FRET.

## Materials and Methods

### Protein Purification, Fluorophore Labeling, and Analysis

To enable the introduction of specific labeling sites in CP, all nine intrinsic Cys residues were changed to Ser. Then, for each single-molecule FRET experimental construct, two Cys residues were introduced on the surface to permit labeling with donor and acceptor fluorophores. Residues were chosen for labeling based on considerations described in the Results section. Dual-labeled variants were maximized using a sequential labeling strategy, and labeling efficiency was confirmed by mass spectrometry. Individual steps are described in detail below.

*Cys-null CP.* Cys residues were changed to Ser using the mutagenesis service of GeneWiz (South Plainfield, NJ). Starting with a bacterial expression plasmid for mouse CPα1β2 (pBJ 2041) ^14^, we changed all nine Cys residues to Ser, producing a “Cys-null” CP expression plasmid (pBJ 2454). This plasmid simultaneously expressed two CP subunits: His-tagged mouse CP α1 (Q5RKN9, *Mus musculus*, C124S, C141S, C157S) and non-tagged mouse CP β2 (Q923G3, *Mus musculus*, C8S, C36S, C62S, C147S, C206S, C272S). The CP heterodimer, assembled from the two co-expressed subunits, was purified as described ^11^. The heterodimer was stable during purification.

We assayed Cys-null CP for two biochemical activities: capping F-actin barbed ends with pyrene-actin polymerization assays and binding to a CARMIL1 CPI-motif peptide with isothermal calorimetry (ITC), both performed as previously described ^7^. The assays for capping F-actin barbed ends showed similar activities for wt and Cys-null CP (Supplementary Figure 1). Cys-null CP and wt CP showed equivalent binding to a CARMIL1 CPI-motif peptide (Supplementary Table 1).

### CP FRET constructs

For each of two FRET constructs, Cys residues corresponding to the chosen labeling positions were introduced in the Cys-null CP expression plasmid (pBJ2454) by GeneWiz. The first construct used α1 S9C and β2 S161C mutations (pBJ 2478), and the second construct used α1 N44C and β2 E84C mutations (pBJ 2488).

### Expression and labeling of CP FRET constructs

CP subunits were co-expressed from one plasmid in *E. coli* NiCo21(DE3) (New England BioLabs, Ipswich, MA) as described ^7^. The fusion protein was isolated on Ni Sepharose^®^ 6 Fast Flow (Cytiva, Marlborough, MA). PreScission protease (GenScript, Piscataway, NJ) was added to cleave the His-tags. Eluted CPs were bound to ceramic hydroxyapatite type I, 40 mm (Bio-Rad, Hercules, CA) in 20 mM Tris-HCl, 20 mM NaCl, 2 mM NaH_2_PO_4_, 125 mM CaCl_2_, 1 mM NaN_3_, 5 mM DTT (pH 7.5) and eluted with a linear gradient to 200 mM NaH_2_PO_4_, 20 mM NaCl, 125 mM CaCl_2_, 1 mM NaN_3_, 5 mM DTT (pH 7.6). Fractions containing purified CPs were dialyzed into 20 mM Bis-Tris-propane, 1 mM Tris(2-carboxyethyl)phosphine (TCEP), 0.1 M NaCl, 1 mM NaN_3_ (pH 7.0).

CPs were diafiltered into 20 mM Bis-Tris-propane, 0.1 M NaCl, 1 mM NaN_3_ (pH 7.0), and immediately labeled with 0.7 mol Alexa Fluor 488 C5 maleimide (Invitrogen, Carlsbad, CA) per mol CP. Alexa 488-labeled CPs were isolated on Mono-Q HR 5/5 (Cytiva) in 20 mM Bis-Tris-propane, 1 mM TCEP, 1 mM NaN_3_ (pH 7.0), and eluted with a NaCl gradient. Alexa 488-labeled CPs were diafiltered into 20 mM Bis-Tris-propane, 0.1 M NaCl, 1 mM NaN_3_ (pH 7.0), and immediately labeled with 5 mol Alexa Fluor 594 C5 maleimide (Invitrogen) per mol CP. Alexa 488, Alexa 594-labeled CPs were isolated on Mono-Q HR 5/5 (Cytiva) in 20 mM Bis-Tris-propane, 1 mM TCEP, 1 mM NaN_3_ (pH 7.0), and eluted with a NaCl gradient. Column fractions were frozen individually at -70°C.

### Mass spectrometry analysis of labeled CP

Labeled CP samples were prepared by exchanging into volatile salt solutions, with a final exchange into 2.5 mM ammonium acetate. The solution was diluted into 50 / 50 acetonitrile / water with 0.1% formic acid for infusion into the mass spectrometer. Intact protein samples were analyzed by direct infusion on a Thermo Scientific Orbitrap Eclipse using Protein Mode. Spectra were collected at 240K with 5 microscans and averaged over 100 scans prior to deconvolution using Thermo Scientific Xtract.

### Labeling of CP Constructs for Isothermal Calorimetry (ITC)

We used ITC to test whether addition of the dyes altered binding of CPI-motif peptides to the CP FRET constructs. The sequential labeling strategy did not produce quantities of dual-labeled donor-acceptor material sufficient for multiple ITC experiments. As an alternative, we labeled the CP FRET constructs with acceptor dye in quantities sufficient for coupling at both Cys residues, and we tested those double-labeled CP preparations by ITC.

Cys-null CP with α1 N44C β2 E84C (pBJ 2488) and Cys-null CP with α1 S9C β2 S161C (pBJ 2478) were expressed and purified using the same methods described above. The CPs were diafiltered into 20 mM Bis-Tris-propane, 0.1 M NaCl, 1 mM NaN3 (pH 7.0), and immediately labeled with 6 mol Alexa Fluor 594 C_5_ maleimide (Invitrogen) per mol CP. Alexa 594-labeled CPs were isolated on Mono-Q HR 5/5 (Cytiva) in 20 mM Bis-Tris-propane, 1 mM TCEP, 1 mM NaN3 (pH 7.0), and eluted with a NaCl gradient. Fractions containing Alexa 594-labeled CPs were dialyzed into 20 mM 3-(N-morpholino)propanesulfonic acid (MOPS), 100 mM KCl, 1 mM TCEP, 1 mM NaN3 (pH 7.2) and stored at -70 °C.

### ITC experiments

ITC experiments were performed as described ^7^. The concentrations of Alexa 594 labeled CPs were determined using UV-visible absorbance with Alexa 594 extinction coefficient ε_594_ = 96,000 M^-1^ cm^-1^. Concentrations of unlabeled CP and CPI peptides were determined as described ^7^. ITC data were fit to a single-site binding model, constraining the stoichiometry to 1.0.

### Single-molecule FRET experiments

Single-molecule FRET measurements were performed on a modified Picoquant MT200 instrument (Picoquant, Berlin, Germany) as described ^15^. Experiments were conducted at a protein concentration of 100-250 pM (estimated from dilutions of samples with known concentration based on absorbance measurements), 20 mM 3-(N-morpholino)propanesulfonic acid (MOPS), 100 mM KCl, 1 mM TCEP, 1 mM NaN3 (pH 7.2), 200 mM β-mercaptoethanol (for photoprotection), 0.001% Tween 20 (for surface passivation), at a room temperature of 295 ± 0.5 K. Pulsed interleaved excitation (PIE) enabled selection of bursts with 1:1 donor:acceptor stoichiometry.

Single-molecule FRET data were analyzed using the “Fretica” package developed by Daniel Nettels and Ben Schuler (University of Zurich, Zurich, CH) and available online at https://schuler.bioc.uzh.ch/wpcontent/uploads/2020/09/Fretica20200915.zip. Fluorescence lifetimes were obtained via a convolution with the Instrument Response Function (IRF), measured as described ^15^. Time resolved anisotropies were computed as described ^15^. Fits for titration curves and 95% confidence intervals were calculated using Mathematica (Princeton, NJ).

### MD simulations and analysis

For molecular simulation, we used crystal structures for free CP (PDB 1IZN) ^16^ and the V-1-CP complex (PDB 3AAA) ^9^ as starting points, rebuilding any missing residues using Modeller ^17^. There is no co-complex structure for WASHCAP and CP, but CD2AP has good sequence alignment with WASHCAP. We used the CD2AP complex PDB 3LK4 ^13^ to build the N-terminal portion of WASHCAP (starting at V992) and PDB 3AA6 ^9^ to build the C-terminal part (up to A1024). The two missing residues at either end of the WASHCAP CPI were added using Modeller ^17^. The C-terminal tentacle of CPβ is dynamic and has a wide range of motion. In control simulations with and without the β-tentacle, we found no significant difference in the dynamics of the rest of CP and therefore truncated CPβ at R530 so that the simulation box would be much smaller, and simulations would run faster.

Each system (free CP, V-1-CP and WASHCAP-CP) was solvated using TIP3P water with 10 Å padding. Na^+^ and Cl^-^ were added to neutralize the system and give an ionic strength of 50 mM. NAMD 3.0 ^18^ was used to perform the simulations using the CHARMM36 forcefield ^19^. Following minimization, the systems were heated and equilibrated at a temperature of 300 °K and 1 atm of pressure (NpT conditions). Simulations were performed with 2 fs timesteps by fixing hydrogens. We employed Particle Mesh Ewald ^20^ for long-range electrostatics and used a 10 Å cut-off and 8.5 Å switch distance for van der Waals interactions. Following 0.5 µs equilibration of each system, 3-5 independent replicates of each system were run for at least 1.5 µs each, resulting in more that 10 µs of total simulation data. Analysis was performed using Bio3D ^21, 22^ in R.

### Calculation of FRET Efficiencies from MD Simulations

Using the full MD trajectories, we adopted the geometric accessible volume (AV) method to simulate and predict FRET efficiencies utilizing the Python module *FRETraj* ^23^. For every conformation of the trajectory, AV projects a spheroid onto the donor and acceptor locations that represents all sterically allowed dye positions ^24^. The spheroid is calculated using five dye parameters: three radii of the dye as well as the width and length of the linker.

## Results

To monitor in-solution conformational changes via single-molecule FRET, we designed full-length CP constructs that interrogate the structural changes upon ligand binding (see Supplementary Text, *Design of Labeling Positions on CP*). We identified two sets of labeling positions that sample different regions of the protein and provide complementary information. CP α1N44C β2E84C (henceforth “α44β84”) and CP α1S9C β2S161C (henceforth “α9β161”) (Figure 1) probe the conformations between the stalk and the lower cap region and between the stalk and the higher cap region, respectively. Cysteine residues were inserted in a Cysteine-less CP variant that maintains the biochemical activity of wild-type CP (see Supplementary Figure 1 and Supplementary Table 1). For each construct, mass spectrometry analyses revealed *quasi-site-specific* labeling of the CP heterodimer, with donor fluorophore predominantly (or exclusively) attached to the β subunit and acceptor fluorophore predominantly or exclusively attached on the α subunit for both constructs (see Methods, Figure 1 and Supplementary Figure 2).

**Figure 2.**
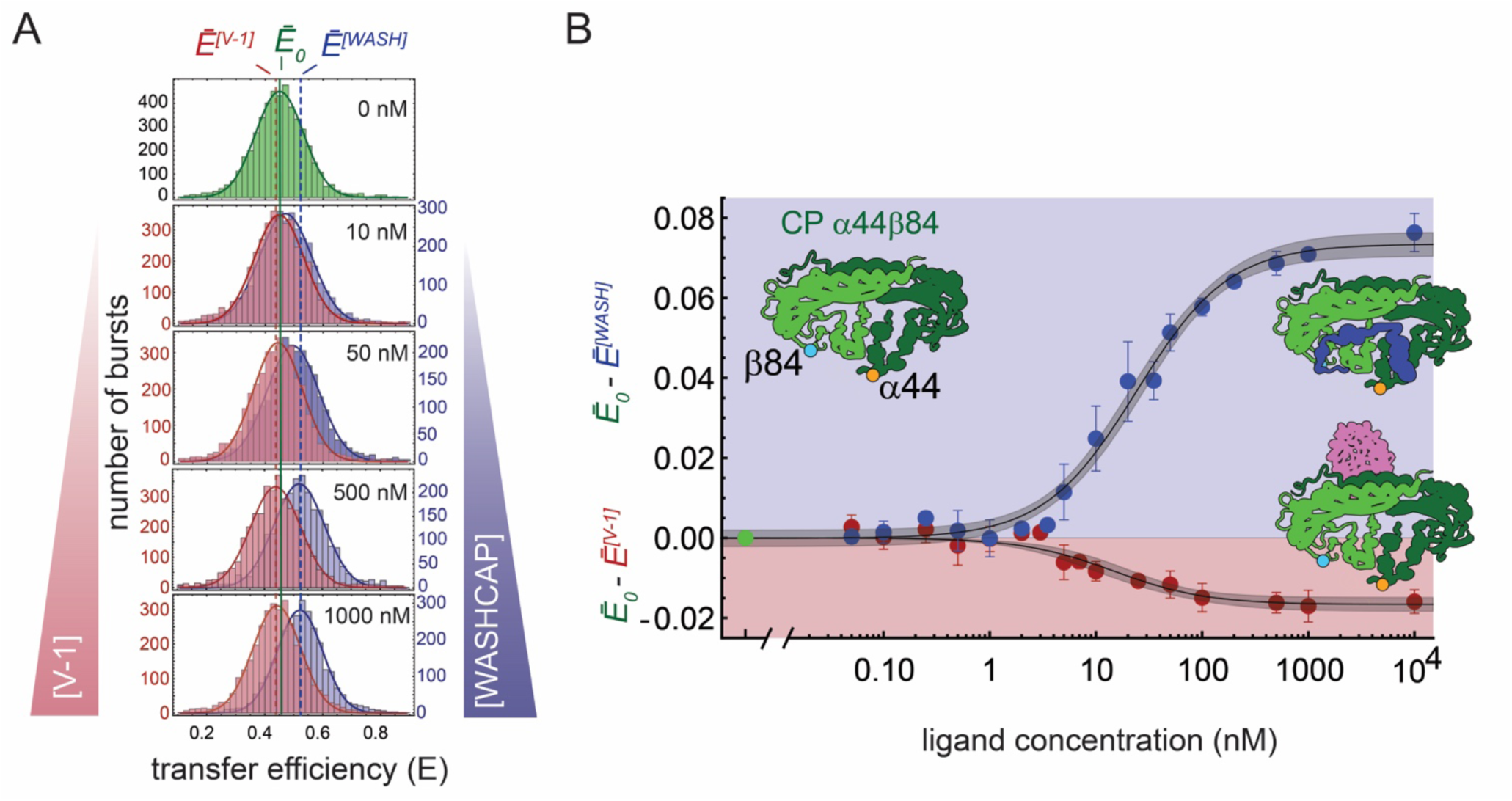
Single-molecule FRET analysis of CP α44β84 binding to WASHCAP and V-1. **A.** Transfer efficiency histograms for free CP (Apo-CP, 0 nM ligand, green) and CP titrated with WASHCAP CPI-motif peptide (blue) or V-1 (red). WASHCAP induces a shift to a higher Ē value, and V-1 causes a shift to a lower Ē value. Measurements were performed in triplicate; representative histograms are shown. **B.** Titration curves for WASHCAP (blue) or V-1 (red) plot the mean normalized change in Ē (Ē_0_ -Ē^[WASH/V-1]^) vs increasing concentrations of WASHCAP or V-1. Plotted data points are mean values, and error bars are standard deviation. Gray shading represents the 95% confidence interval of the fit as determined by Mathematica.

Finally, we tested whether inserting Cysteine residues at specific probe positions in CP, coupled with the addition of fluorescent dyes, affected the binding of CPI-motif peptides and V-1 (see Methods). Dissociation binding constants (K_D_) obtained from ITC measurements with double-acceptor labeled CP are listed in Table 1. The K_D_ values for CPI-motif peptides were similar to ones obtained from ITC measurements with wild-type CP (McConnell *et al.* 2020 PMID 32133840, and Supplementary Table 1), and the K_D_ values for V-1 were similar to values for wild-type CP (Supplementary Table 1).

**Table 1.**
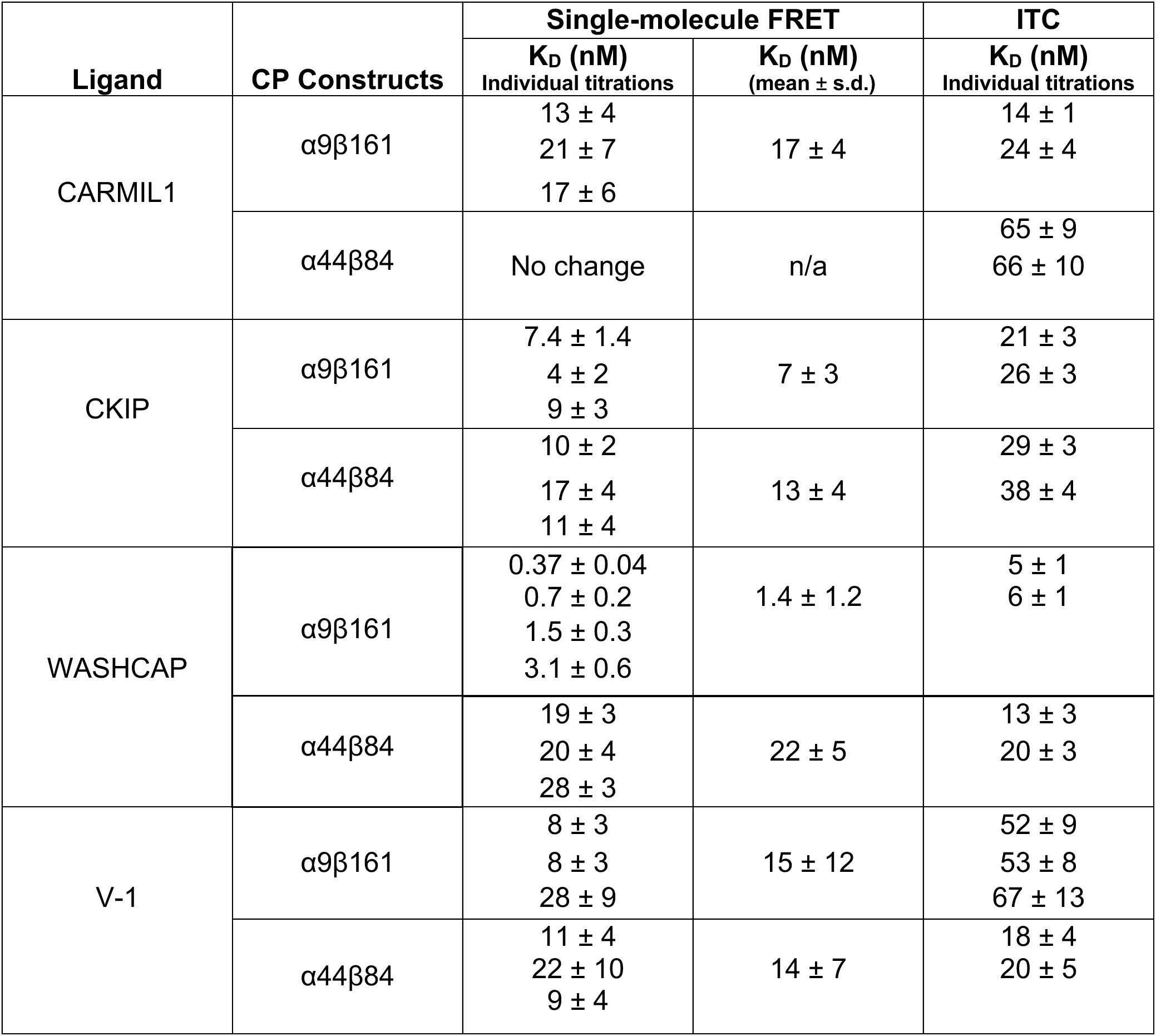
K_D_ values for ligand binding to dye-labeled CP constructs measured by single-molecule FRET and ITC. For single-molecule FRET, CP α9β161 and CP α44β84 were labeled with donor and acceptor dyes. K_D_ values are calculated from the change in transfer efficiency as a function of the specified ligand. For ITC, CP α9β161 and CP α44β84 were labeled with acceptor dye at both donor and acceptor positions. K_D_ values are from ITC titrations with specified ligand.

| Ligand | CP Constructs | Single-molecule FRET |  | ITC |
| --- | --- | --- | --- | --- |
| | | $K_D$ (nM)<br>Individual titrations | $K_D$ (nM)<br>(mean ± s.d.) | $K_D$ (nM)<br>Individual titrations |
| CARMIL1 | $\alpha 9\beta 161$ | 13 ± 4<br>21 ± 7<br>17 ± 6 | 17 ± 4 | 14 ± 1<br>24 ± 4 |
| | $\alpha 44\beta 84$ | No change | n/a | 65 ± 9<br>66 ± 10 |
| CKIP | $\alpha 9\beta 161$ | 7.4 ± 1.4<br>4 ± 2<br>9 ± 3 | 7 ± 3 | 21 ± 3<br>26 ± 3 |
| | $\alpha 44\beta 84$ | 10 ± 2<br>17 ± 4<br>11 ± 4 | 13 ± 4 | 29 ± 3<br>38 ± 4 |
| WASHCAP | $\alpha 9\beta 161$ | 0.37 ± 0.04<br>0.7 ± 0.2<br>1.5 ± 0.3<br>3.1 ± 0.6 | 1.4 ± 1.2 | 5 ± 1<br>6 ± 1 |
| | $\alpha 44\beta 84$ | 19 ± 3<br>20 ± 4<br>28 ± 3 | 22 ± 5 | 13 ± 3<br>20 ± 3 |
| V-1 | $\alpha 9\beta 161$ | 8 ± 3<br>8 ± 3<br>28 ± 9 | 15 ± 12 | 52 ± 9<br>53 ± 8<br>67 ± 13 |
| | $\alpha 44\beta 84$ | 11 ± 4<br>22 ± 10<br>9 ± 4 | 14 ± 7 | 18 ± 4<br>20 ± 5 |

Having established that the double-labeled constructs were well-behaved, with biochemical properties similar to wild-type CP, we investigated the ligand-induced conformational changes *via* single-molecule FRET.

### Conformational Changes Between Stalk and Lower Cap Region

The α44β84 construct probes the configuration between the stalk and the underside of the cap region of CP (Figure 1). In absence of ligand (Apo-CP), the histogram of FRET efficiencies reveals one population with mean transfer efficiency (Ē) of 0.458 ± 0.007 (Figure 2 A). The plot of donor lifetime vs Ē suggests that the transfer efficiency population reflects a dynamic ensemble of the protein and not simply a single rigid distance (Supplementary Figure 3). After accounting for the contribution of the dye linker, we estimated the distance distribution sampled by the protein (Supplementary Figure 4), which can be compared to the value calculated from the Apo crystal structure (PDB: 1IZN). The single-molecule measurements report on a dynamic ensemble with a root mean square distance of approximately 55 Å (Supplementary Figure 3-5). The crystal structure C_α_ – C_α_ distance is 45 Å, which is within the distribution sampled by the observed conformational ensemble.

**Figure 3.**
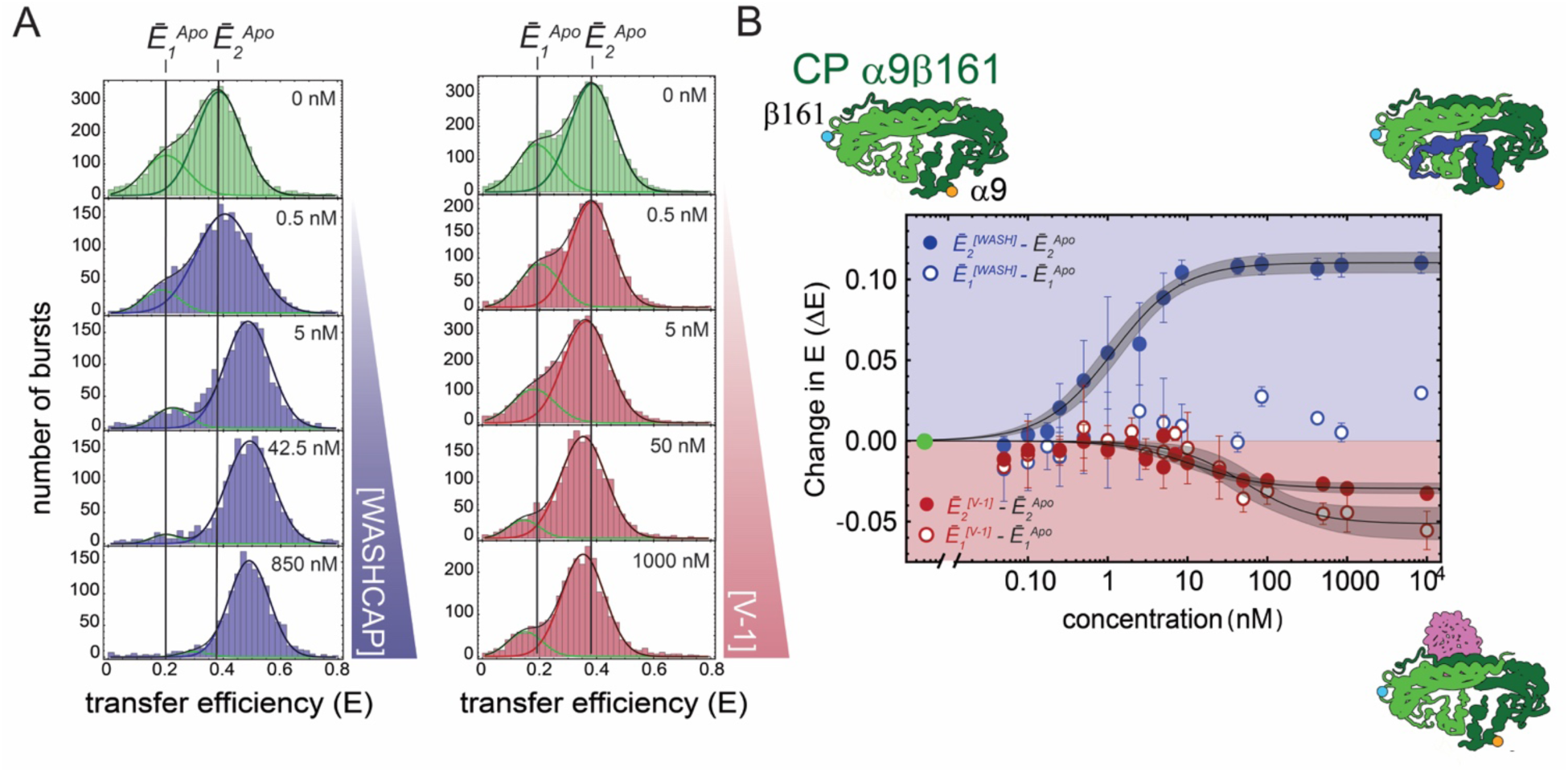
Conformational changes observed for CP α9β161 upon binding of WASHCAP CPI-motif peptide or V-1. **A.** Two conformations are observed for Apo-CP, indicated as E_1_^Apo^ and E_2_^Apo^. For WASHCAP (blue), the titration histograms show a complete shift to a higher mean transfer efficiency for both populations upon saturation. Titration histograms for V-1 (red) show a shift to a lower mean transfer efficiency for both E_1_ and E_2_ populations upon binding; however, a portion of the low-efficiency population remains, in contrast to the results for WASHCAP. Measurements were performed in triplicate. Representative histograms are shown. **B.** Titration curves for WASHCAP (blue) or V-1 (red) plot the mean normalized change in Ē_1_ (Ē_1_^[WASH]^- Ē_1_^Apo^ and Ē_1_^[V-1]^- Ē_1_^Apo^, open circles) or Ē_2_ (Ē_2_^[WASH]^- Ē_2_^Apo^ and Ē_2_^[V-1]^- Ē_2_^Apo^, closed circles) *vs* increasing concentrations of WASHCAP or V-1. Each data point is the mean of three values. Error bars indicate standard deviation. The gray shading represents the 95% confidence interval of the fit. Data for Ē_1_^[WASH]^- Ē_1_^Apo^ do not display a significant difference with addition of WASHCAP; therefore, a fit was not performed.

**Figure 4.**
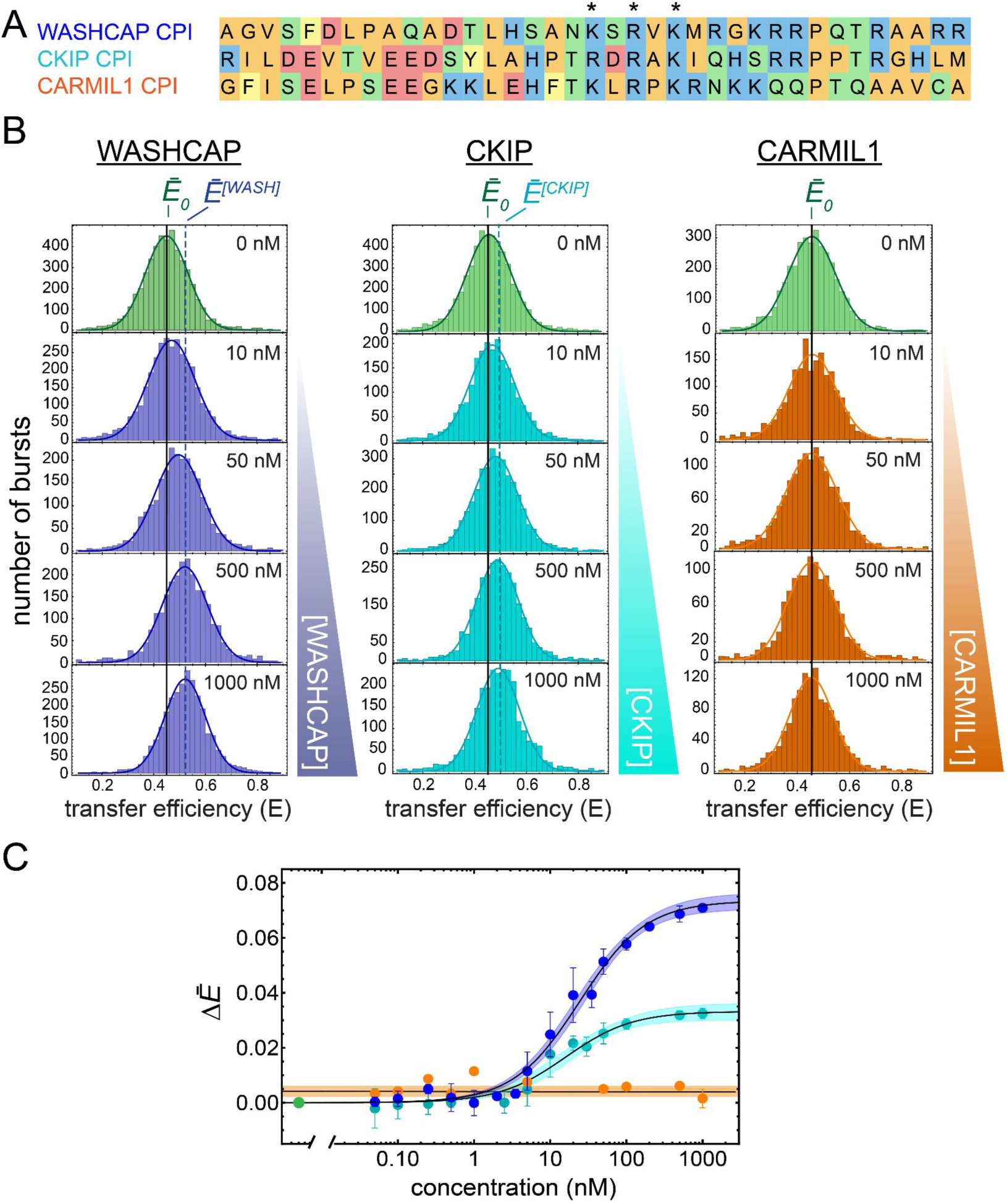
Conformational changes observed for CP α44β84 upon CPI-motif peptide binding. **A.** Sequences of the CPI-motif peptides. Asterisks denote key residues necessary for the biochemical activity of the CPI-motif peptides on CP. **B.** FRET distributions and titration curves for WASHCAP (blue), CKIP (cyan) and CARMIL1 (orange). Measurements were performed in triplicate, and representative histograms are shown. **C.** Titration curves plot the change in mean transfer efficiency (ΔĒ) with increasing concentrations of WASHCAP (blue), CKIP (cyan) and CARMIL1 (orange). Data points are mean and standard deviation from the replicates. Color shading indicates the 95% confidence interval of the fit. Data of CARMIL1 are not fitted because a binding curve is not apparent; in this case, the line denotes the average of the values, and the shading is standard deviation.

**Figure 5.**
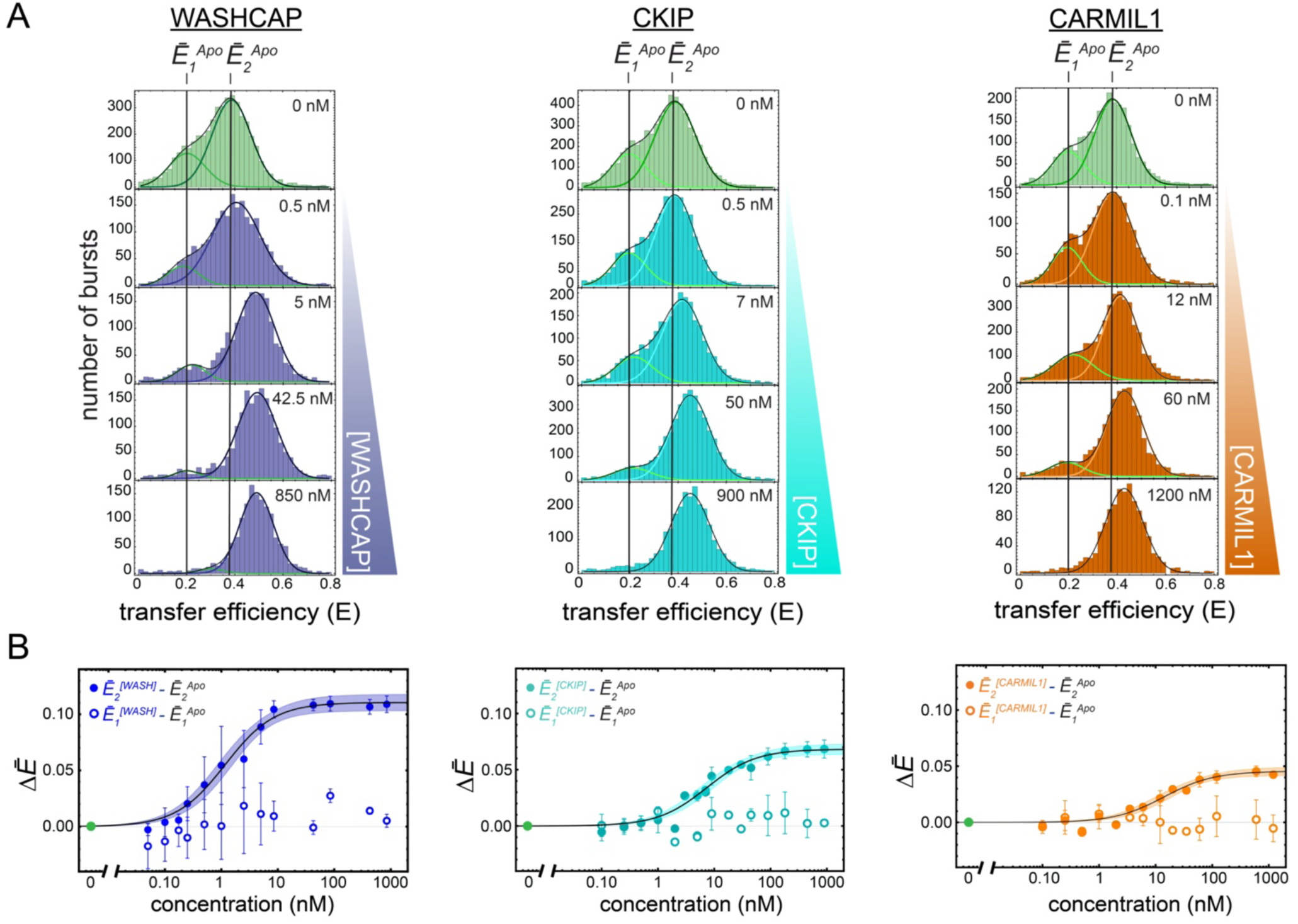
Conformational changes observed for CP α9β161 upon CPI binding. **A.** Transfer efficiency histograms for WASHCAP (blue), CKIP (cyan) and CARMIL1 (orange) CPI-motif peptides. Measurements were performed in triplicate, and representative histograms are shown. **B.** Titration curves plot the mean normalized change in transfer efficiency Ē_1_ (Ē_1_^[WASH]^- Ē_1_^Apo^, Ē_1_^[CKIP]^- Ē_1_^Apo^ and Ē_1_^[CARMIL1]^- Ē_1_^Apo^, open circles) or Ē_2_ (Ē_2_^[WASH]^- Ē_2_^Apo^, Ē_2_^[CKIP]^- Ē_2_^Apo^ and Ē_2_^[CARMIL1]^- Ē_2_^Apo^, closed circles) *vs* increasing concentrations of WASHCAP (blue), CKIP (cyan) and CARMIL1 (orange). Titrations were performed in triplicate in separate experiments and plotted as the mean and standard deviation. Shading represents the 95% confidence interval of the fit.

CPI motifs bind directly to the stalk of CP (Figure 1). To understand whether CPI-motif binding affects the conformations of the stalk region, we performed single-molecule FRET measurements in the presence of the CPI-motif peptide from WASHCAP, which is a potent regulator of CP ^7^. With increasing concentration of ligand, the distribution of transfer efficiencies shifted to higher values and reached saturation at a value above 1 μM (Figure 2 B). The change in Ē was sufficiently small (ΔE^[WASH]^ = +0.071 ± 0.001, Supplementary Table 2) that one cannot resolve two distinct peaks for the Apo-CP and ligand-bound states. The titration curve of Ē vs ligand concentration is well fit assuming a 1:1 binding model, which results in a K_D_ of 22 ± 5 nM (Table 1 and Figure 2 B), similar to the values from corresponding ITC measurements for CP labeled at both Cys residues with the acceptor dye (Table 1 and Supplementary Figure 6).

**Figure 6.**
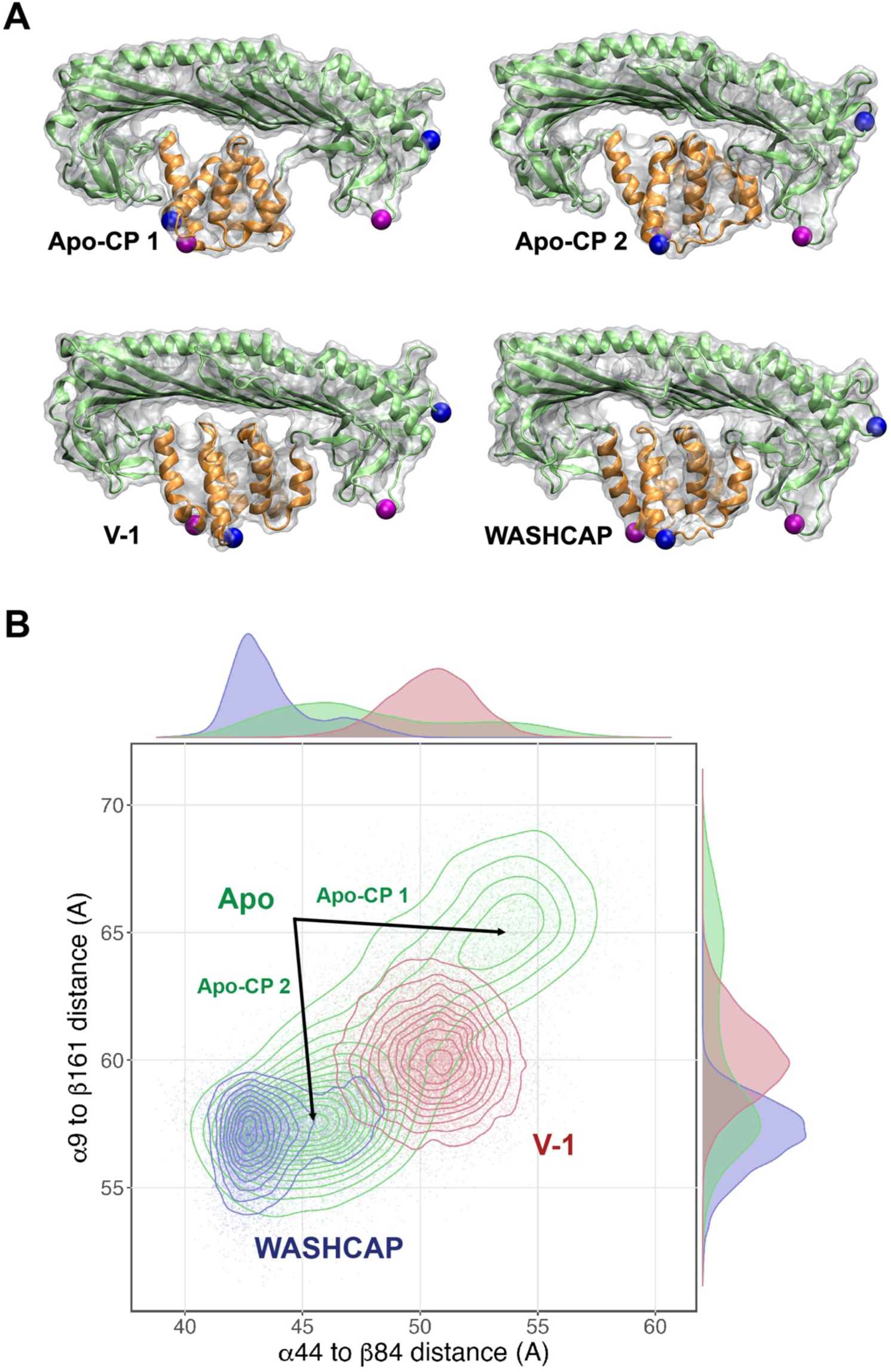
Molecular dynamics simulation results: Equilibrium conformations for the two states of unbound CP (Apo- CP 1 and Apo-CP 2) and for the CP-V-1 and CP-WASHCAP complexes. **A.** The Apo-CP 1 and Apo- CP 2 states primarily differ in the tilt and twist of their stalk regions, relative to the cap. The V-1 and WASHCAP structures have very similar stalk structures, in which the α helices are more aligned with each other than they are in the Apo-CP structures. The locations of the FRET probes are shown in blue (α9 and β161) and magenta (α44 and β84). **B.** MD simulations of C_α_ – C_α_ distance distributions for the attachment points of the fluorophores in the two constructs, comparing Apo-CP (green) with the bound states of V-1 (red) and WASHCAP CPI peptide (blue). Abscissa values are for α44β84, and ordinate values are for α9β161. The distance for the V-1-bound state is longer than the primary Apo-CP 2 state for both FRET pairs while the WASHCAP state is shorter for both pairs.

A structural interpretation of the shift in transfer efficiency as a function of distance requires one to rule out contributions from quenching or rotational hindrance of the fluorophores. To this end, we analyzed the fluorescence lifetime of the donor-only and corresponding anisotropy decays in Apo-CP with and without ligand (Supplementary Tables 3-4). No significant changes were apparent, excluding quenching and rotational effects Therefore, the transfer efficiency and the fluorescence lifetime of the donor in presence of acceptor are most consistent with a small decrease in the mean distance between the positions probed by the dyes (∼ 3 Å) as well as a reduction in the width of the corresponding distance distribution (Supplementary Figure 3-5). This distribution encompasses distances calculated from crystal structures of other CPI-motif peptides in co-complex with CP, suggesting that our observed conformational changes are compatible with previously determined structures.

In biochemical assays, CPI-motif peptides antagonize the binding of V-1 to CP, and V-1 antagonizes CPI-motif binding to CP ^6, 7^. To investigate this mechanism, we measured how binding of the V-1 ligand on the top surface of the CP cap affects the conformation of the stalk and the underside of the cap, where CPI-motif peptides bind (Figure 1). With increasing concentrations of V-1 (Figure 2 A and B), the mean transfer efficiency Ē monotonically shifted to lower values. Most significantly, the change in Ē for V-1 moved in the opposite direction (more expanded with respect to Apo-CP) compared to the change in Ē produced by the WASHCAP CPI-motif peptide (more collapsed with respect to Apo-CP) (Supplementary Table 2, Figure 2). The magnitude of the change in Ē from the Apo state to saturation was small, ΔE^[V-1]^ = -0.017 ± 0.004. The fact that V-1 induced a decrease in Ē is consistent with the difference in distance between residues α44 and β84 in corresponding crystal structures ^9, 13, 16^. No significant alterations are introduced in the width of the distance distribution (Supplementary Figures 3-5), when compared to the Apo configuration. Fit of the change in transfer efficiency yields a K_D_ of 14 ± 7 nM, assuming a 1:1 stoichiometry, in agreement with K_D_ values obtained by ITC (Table 1 and Figure 2 B) and with previously published values for binding affinity and stoichiometry ^7, 8^.

### Conformational Changes Between Stalk and Upper Cap Region

The CP α9β161 construct probes the configuration between the stalk and the upper cap region of CP (Figure 1), which is the region where CP interacts with F-actin barbed-ends and V-1. In contrast to α44β84, the distribution of transfer efficiencies of Apo-CP α9β161 contained two populations (Ē_1_^Apo^ = 0.18 ± 0.02 and Ē_2_^Apo^ = 0.37 ± 0.03, Figure 3 A, top histograms, and Supplementary Table 2), suggesting two conformational states of the protein. This result was not predicted by previous structural studies of CP, including X-ray crystal structures. Analysis of fluorescence anisotropies (Supplementary Table 4) indicated that the two populations do not arise because of slow rotation of the fluorophores; instead, they reflect two different slow-exchange (>1 ms) configurations of CP.

Upon titration with WASHCAP peptide, the Ē_1_^Apo^ and Ē_2_^Apo^ populations decreased in favor of a new third population with change in mean transfer efficiency Ē_2_^[WASH]^ - Ē_2_^Apo^ = +0.112 ± 0.006 (Figure 3, panel A and B, Supplementary Table 2, and Supplementary Figure 7). At saturation with ligand, only the Ē_2_^[WASH]^ population was observed. The transfer efficiency change of the main population can be fitted to a 1:1 binding model, which reports a K_D_ of 1.4 ± 1.2 nM (Table 1 and Figure 3 B). To better account for the multiple states identified, we estimated the relative fraction of each population (associated with the mean transfer efficiencies Ē_1_^Apo^, Ē_2_^Apo^, and Ē^[WASH]^) by computing the corresponding areas under the curve at each ligand concentration (Supplementary Figure 8 A and B, WASHCAP). We then performed a global fit of the three populations, assuming a 3-state model (Apo-CP 1, Apo-CP 2, Bound) (Supplementary Text and Supplementary Figure 9-10). From the model, we computed an effective K_D_*, representing the dissociation constant between the Apo and bound configurations, of 2 ± 2 nM, which is similar to the values obtained by ITC for labeled molecules (Supplementary Table 5). However, a close inspection of histograms reveals a change in the equilibrium between the two Apo configurations upon addition of ligands, indicating that a 4-state model (Apo-CP 1, Apo-CP 2, Bound 1, Bound 2) may be more appropriate (Supplementary Figure 9-10). It is important to note that connectivity between different states cannot be ruled out in equilibrium experiments; therefore, we considered the most generic case where both Apo states can bind the ligand and the bound states can interconvert one into another. Corresponding analysis reveals two dissociation constants K_D1_ and K_D2_ (between the corresponding Apo and Bound states, Supplementary Table 6) equal to 20 ± 20 and 1 ± 1 nM, respectively. The one order of magnitude difference in the K_D1_ value explains why we can adequately describe the data with a 3-state model, since the contribution from the Bound 1 state is negligible.

**Figure 7.**
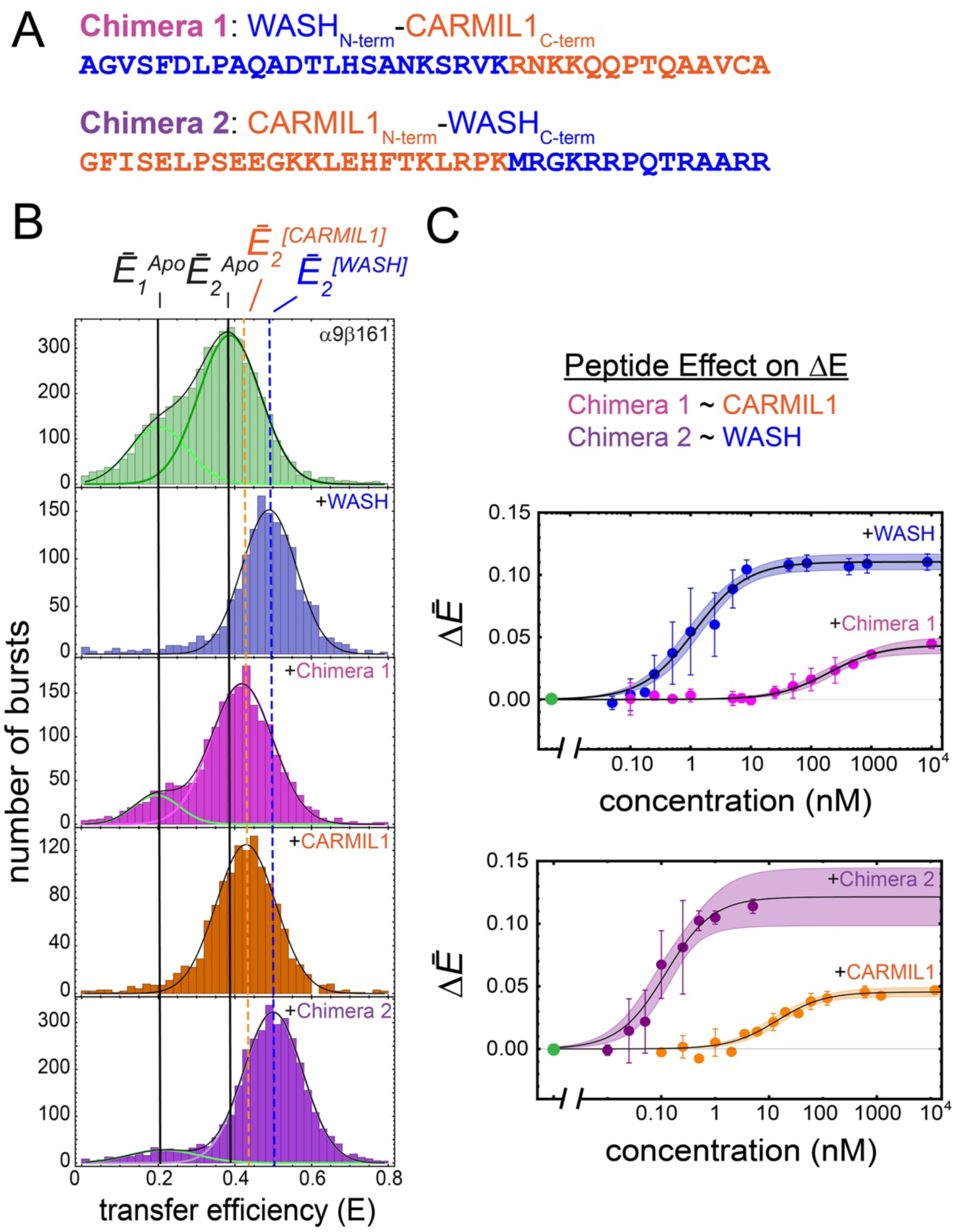
Dissection of CPI-motif peptide effects with chimeras. **A**. Sequences of chimeric peptides, WASH_N_-C1_C_(Chimera 1) and C1_N_-WASH_C_ (Chimera 2). CARMIL1 residues are orange, and WASHCAP residues are blue. **B**. Transfer efficiency histograms for CP α9β161 with WASHCAP (blue), Chimera 1: WASH_N_ - C1_C_ chimera (fuchsia), CARMIL1 (orange), and Chimera 2: C1_N_ - WASH_C_ chimera (purple). Experiments were performed in triplicate and representative histograms are shown for ligand saturation at 1 μM. **C**. Titration curves for all concentrations of peptide are shown with the average change in mean transfer efficiency relative to Apo-CP, ΔĒ, plotted versus concentration of peptide. The value for Ē_1_^Apo^ did not change with chimeric peptide concentration, similar to results for WASHCAP and CARMIL1 peptides.

**Figure 8.**
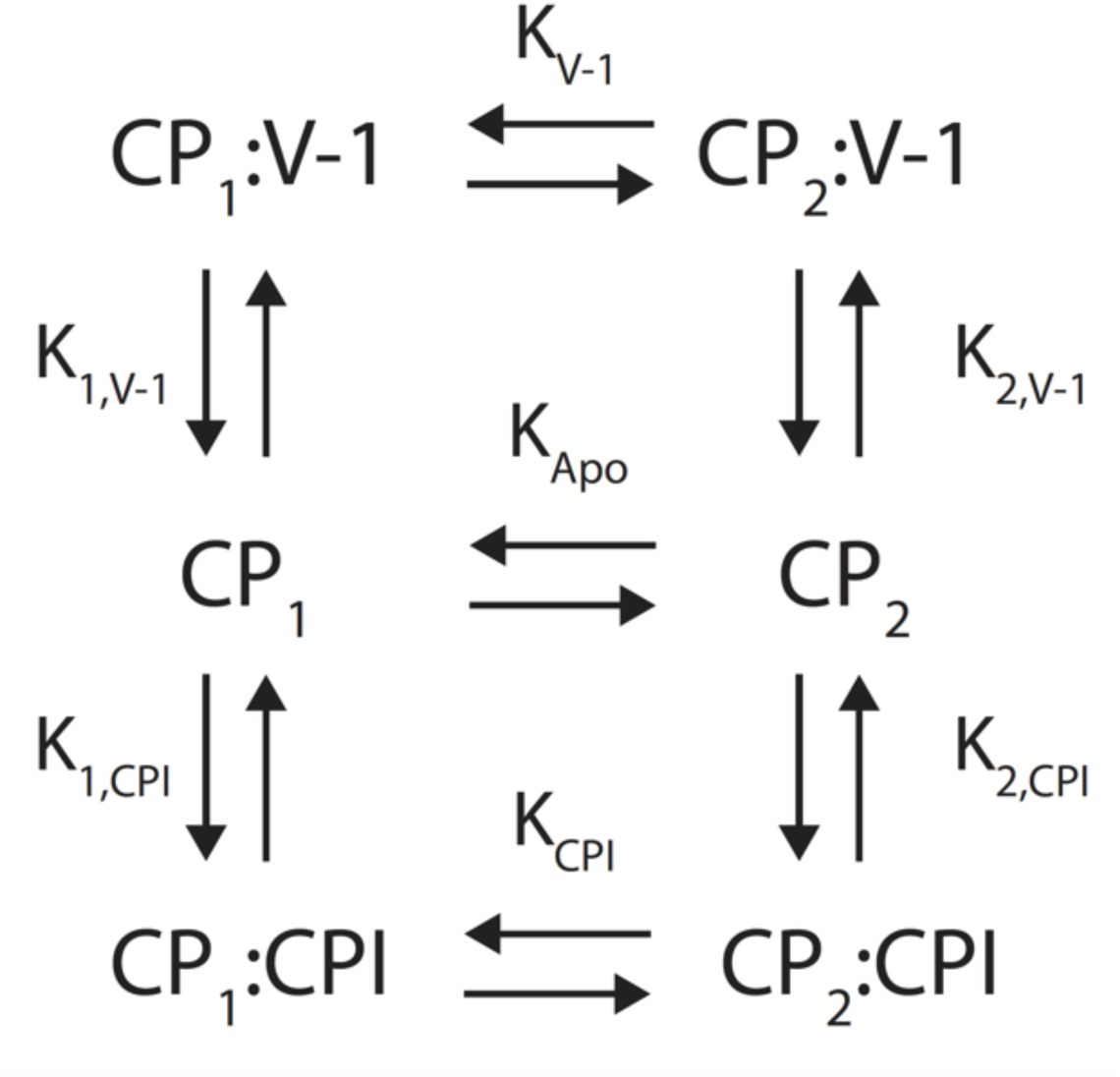
Equilibrium scheme of CP states in Apo and bound states. CP interconverts between two Apo unbound configurations, Apo-CP 1 (CP_1_) and Apo-CP 2 (CP_2_), which can both bind to either V-1 or CPI motif peptides. Equilibrium association constants are shown between each state.

Next, we titrated CP α9β161 with increasing concentrations of V-1. The E_1_^Apo^ and E_2_^Apo^ peaks shifted toward lower mean transfer efficiency values, with ΔĒ_1_^[V-1]^ = -0.056 ± 0.012 and ΔĒ_2_^[V-1]^ = -0.033 ± 0.003 (Figure 3, panel A and B, and Supplementary Table 2, and Supplementary Figure 11). These changes are in the same direction as the changes observed for the CP construct α44β84 (Supplementary Figure 5). Fitting the change in transfer efficiency as a function of the V-1 concentration results in a K_D1_ of 50 ± 20 nM for the Apo-CP 1 state (representing the transition from unbound to Bound 1) and a K_D2_ of 12 ± 3 nM for the Apo-CP 2 state (representing the transition from unbound to Bound 2) (Figure 3). To better assess the binding affinity in the model, we globally analyzed the distribution of transfer efficiencies assuming a 4-state model (Apo-CP 1, Apo-CP 2, Bound 1, Bound 2). The global fit of the titration provides a K_D1_ of 25 ± 2 nM for V-1 binding to Apo-CP 1 and K_D2_ of 19 ± 2 nM for V-1 binding to Apo-CP 2 (Supplementary Figure 12 and Supplementary Table 6). Reducing the model to represent only three states does not reproduce the experimentally determined distribution of transfer efficiencies at high (μM) ligand concentration (Supplementary Figure 12-13).

Therefore, our experimental results are consistent with the coexistence of four states across the titration. The similarity between the two determined dissociation constants, K_D1_ and K_D2_, is consistent with the coexistence of the two bound populations at saturating concentration and the overall values are similar to the corresponding ITC results (Supplementary Table 5).

Comparison of lifetime vs transfer efficiency for Apo-CP and V-1 bound states suggests that the binding of V-1 does not significantly alter the distribution of conformations, rather impacts the relative abundance of the two configurations (Supplementary Figures 3-5).

### CPI sequence-specific effects on CP conformations

The amino-acid sequences of CPI-motif regions from different CPI-motif proteins are conserved across evolution and CPI-motif peptides from those proteins have distinct biochemical effects on CP ^7^. We investigated whether these biochemical differences stem from disparate conformational changes upon binding. To this end, we compared how interaction of WASHCAP affects CP conformations differently from two other CPI-motif peptides: CKIP and CARMIL1 (Figure 4 A).

As discussed above, for CP α44β84, binding of WASHCAP produced a positive increment of ΔĒ^[WASH]^ = 0.071 ± 0.001 (Figure 4 B and C and Supplementary Table 2). The interaction with CKIP resulted in a smaller increase, +0.031 ± 0.002, whereas the binding of CARMIL1 produced no significant shift in Ē (Figure 4 B and C and Supplementary Table 2). To exclude the possibility that the absence of changes for CARMIL1 is due to a lack of interaction of the peptide with the labeled CP, we verified the binding by ITC (Table 1). To confirm that these results reflect conformational variations, we measured fluorescence lifetimes and time-resolved anisotropy for the donor-only and acceptor-only populations of the Apo-CP and the CPI-motif-saturated CP samples. No significant differences in lifetimes were observed, indicating the absence of dye quenching and a lack of deviations from the assumption of rapid rotation of the fluorophores (Supplementary Table 3). Analysis of the donor dye lifetime in presence of the acceptor (at saturating conditions of ligand) suggests that binding of CARMIL1 does restrict the distance distribution explored by CP while maintaining an average distance similar to that seen for Apo-CP (Supplementary Figures 3- 5). In contrast, the changes in Ē with WASHCAP and CKIP represent structural variations characterized by larger mean squared inter-dye distances and equal or broader distributions of distances compared to the Apo-CP (Supplementary Figures 4-5).

Next, we compared the effect of the three CPI-motif peptides on CP α9β161. Similar to results for CP α44β84, CKIP produced a smaller shift in transfer efficiency, ΔĒ_2_^[CKIP]^ = +0.068 ± 0.008 (Supplementary Table 2) than WASHCAP, ΔĒ_2_^[WASH]^ = +0.112 ± 0.006 (Figure 5 A and B and Supplementary Table 2). Different from CP α44β84, we observed a measurable change in the mean transfer efficiency for CARMIL1, ΔĒ_2_^[CARMIL1]^ = +0.042 ± 0.002 (Supplementary Table 2). Corresponding values for K_D_ are reported in Table 1, and data are shown in Supplementary Figures 7 and 8. Analysis of the lifetime versus transfer efficiency for CP α9β161 reveals that interaction with both CARMIL1 and WASHCAP reduces the distribution of distances sampled by CP, whereas interaction with CKIP adopts a similar distribution to Apo-CP (Supplementary Figures 3-5).

One important difference among the three CPI-motif peptides is that they produce quantitatively different changes in mean transfer efficiency and lifetime for both CP constructs, α44β84 and α9β161, indicating that the extent to which the CP conformations are modulated upon interaction with the CPI-peptide depends on the nature of the CPI-peptide ligand itself. Comparing the distance distributions determined from lifetime analysis of the CARMIL1 – CP interactions for both α44β84 and α9β161 probe sets, indicates that the CPI-motif of CARMIL1 has the strongest effect on restricting CP conformations, whereas CKIP has the least effect allowing for a broader dynamic ensemble of CP (Supplementary Figures 3-5).

### Atomistic models of conformational ensembles from MD

The single-molecule FRET results described herein reveal two distinct conformational states of Apo-CP, which have not been observed previously. These results also reveal small but significant structural changes in response to the binding of different regulatory ligands. Single-molecule FRET experiments provide a simple coarse-grained perspective (single distance per residue pair studied) of the conformational changes in CP upon binding. Therefore, to investigate and understand the possible nature of conformational changes throughout the entirety of the CP molecule, we employed a molecular dynamics approach to construct atomistic models of the conformational states in the Apo and ligand-bound conditions. Simulations were performed for each starting configuration based on the protein crystal structures in the absence of ligand, with WASHCAP CPI, and with V-1 (See Methods). Multiple independent simulations were performed in each case, with over 10 µs of aggregate simulation time.

For Apo-CP, MD simulations show two stable conformations. This result supports the experimental observation of two states for CP α9β161 in the single-molecule FRET measurements. Movies showing interpolations between the two equilibrium states reveal the stalk region of CP undergoing twisting and tilting (Figure 6 A, Movie 1 and Supplementary PDB files). One state, termed Apo-CP 2, was occupied more often than the other, termed Apo-CP 1 (illustrated in Figure 6 A and Supplementary Figure 14). This difference is qualitatively consistent with the relative abundance of the two populations observed in the single-molecule FRET experiments.

The binding of ligand, for both V-1 and WASHCAP CPI peptide, results in the α helices of the stalk region of CP becoming more aligned with each other, relative to the two Apo states. The effect on the mushroom cap is also similar for both ligands, resulting in a flatter cap as compared to the Apo states (illustrated in Figure 6 A, Movie 2, Movie 3 and Supplementary PDB files). Focusing on the dye positions in the cap region, we observe that they show relatively small changes between the Apo, V-1 and WASHCAP states (Figure 6 A). Conversely, the helices in the stalk region show changes in both alignment and rotation of the helix bundle, and this leads to larger changes in the dye positions, and hence, the distances between FRET pairs.

These results are consistent with the conclusions of the single-molecule FRET experiments, in which CPI-motif binding results in the dyes moving closer to one another and V-1 binding results in the dyes moving farther apart. We note that only a single population is reported for both WASHCAP and V-1 bound CP conformations in the simulation (Figure 6 B and Supplementary Figure 14). This outcome is consistent with WASHCAP single-molecule measurements but is at variance for what concerns V-1, where two bound populations are experimentally determined at saturation of ligand (Figure 3). We interpret this discordance between experiments and simulations as an indication of an incomplete exploration of the protein energy landscape in the bound state (where low frequency states may be rarely explored) or a small variation in energetics (temperature).

To provide a more detailed comparison of simulations and experiments, we used the molecular dynamics simulation trajectories to calculate transfer efficiencies as would be expected for a confocal single-molecule FRET experiment. Using the conformations from the simulations, a dye cloud was generated for each fluorophore to illustrate the possible dye conformations (Supplementary Figure 15), using the Accessible Volume method described by Sindbert and colleagues ^24^. These dye clouds, coupled with photon detection statistics (shot-noise), then allowed us to calculate a distribution of transfer efficiencies for each CP construct in Apo conformation and when bound to V-1 and WASHCAP.

For CP α44β84, the two different conformational states of the protein result in two populations with mean transfer efficiencies of 0.407 and 0.567, respectively (Supplementary Figure 14). This differs from the single-molecule FRET measurements of a single peak with mean transfer efficiency of 0.458 in the experiments (Figure 2). With WASHCAP CPI peptide bound, the simulated transfer efficiency shifts to a higher value (Supplementary Figure 14), which agrees qualitatively with the experimental measurements for single-molecule FRET (Figure 2). In addition, when V-1 is bound, the simulated transfer efficiency shifts to a lower value (Supplementary Figure 14), again consistent with the results of single-molecule FRET (Figure 2).

For the construct CP α9β161, the transfer efficiency distribution from the two Apo-CP states reflects the corresponding fractions observed in the simulations and partially overlaps (Supplementary Figure 14). This result agrees qualitatively with what was observed by single-molecule FRET (Figure 3), though predicted mean transfer efficiencies are lower than the detected ones. We speculate that the shift in measured transfer efficiency reflects changes in the Förster radius due to alterations in the κ^2^ parameter (see Supplementary Text and Supplementary Table 7). We cannot, however, exclude that the two Apo-CP states observed in simulation would be detected as a single averaged mean transfer efficiency in the single-molecule FRET experiment if their exchange occurs on a timescale faster than a millisecond (approximately the diffusion time of the protein). A weighted average of the simulated mean transfer efficiencies of 0.14 and 0.22 is indeed consistent with the experimentally measured mean value of Ē_1_^Apo^ = 0.18 ± 0.02, suggesting that observation of Ē_2_^Apo^ = ∼ 0.37 in the simulations would require a further exploration of the protein energy landscape. In the following, we restrict the interpretation of simulations and their comparison with experiments to the first scenario, where discrepancies are attributed to alterations in the κ^2^ parameter.

With WASHCAP CPI-motif peptide bound, the simulated transfer efficiency shifts to a single population at a higher transfer efficiency (Figure 6 B and Supplementary Figure 14). This result corresponds to what was measured by single-molecule FRET upon WASHCAP binding (Figure 3). With V-1 bound, the simulated transfer efficiency shifts to a single population at a higher efficiency (Supplementary Figure 14, α9β161, lower panel). This result is not predicted from the C_α_ – C_α_ distance measurements provided by the molecular dynamics simulation (Figure 6 B). In addition, this result differs from the result for single-molecule FRET. The single-molecule FRET results showed a shift to a lower mean transfer efficiency for both Apo states, and both states were present after V-1 saturation (Figure 3).

Overall, the MD simulation results are in general agreement with the single-molecule FRET results and conclusions, within known limitations in the conversion of distances from simulated data to mean transfer efficiencies. The MD simulations provide a consistent atomistic description of the conformational changes that occur in CP following regulator ligand binding. This information provides new insight into possible details of the allosteric mechanism of CP regulators.

### Dissecting the CPI-motif

The three CPI-motif peptides produced consistently different results for both CP constructs, α9β161 and α44β84. We asked which portions of the CPI-motif peptides were responsible for the differences. Examining the amino-acid sequences and performing whole-molecule alignment of crystal structures of the CP / CARMIL1 (PDB 3LK2) and CP / CKIP (PDB 3AA1) peptide complexes, we observed that the α-carbon backbones in the N-terminal halves of the CARMIL1 and CKIP peptides are in very good alignment, while the α-carbon backbones differ substantially in the C-terminal halves (Supplementary Figure 16 and Movie 4). We hypothesized that the single-molecule FRET differences between the CARMIL1 and CKIP CPI-motif peptides might result from the differences in their C-terminal halves. To test this hypothesis, we created chimeras of CPI-motif peptides, combining the N-terminal and C-terminal halves of WASHCAP (which had the largest changes in mean transfer efficiencies) with those of CARMIL1 (which had the smallest changes in mean transfer efficiencies). The dividing line between the N-terminal half of one peptide and the C-terminal half of the other peptide was chosen as C-terminal to the central core of the most highly conserved amino-acid residues essential for all CPI-motif peptide binding to CP, namely **K**X**R**X**K** (Figure 7 A) ^25, 26^ . We refer to these two chimeras as WASH_N_-C1_C_ (N-term WASHCAP with C-term CARMIL1) and C1_N_-WASH_C_ (N-term CARMIL1 with C-term WASHCAP).

The chimeric peptides were titrated into CP α9β161 to saturation, and the corresponding transfer efficiency distributions were compared to each other and to those of the wild-type (non-chimeric) peptides. The Ē difference upon binding of WASH_N_-C1_C_ was ΔĒ_2_^[WN-C1C]^ = +0.047 ± 0.004 (Figure 7 C and Supplementary Table 2), which is equivalent to the change observed for CARMIL1 of +0.042 ± 0.002 (Supplementary Table 2). The change in Ē upon binding of C1_N_-WASH_C_ was ΔĒ_2_^[C1N-WC]^ = +0.114 ± 0.006 (Supplementary Table 2), which is within errors of the +0.112 ± 0.006 change observed for WASHCAP (Supplementary Table 2). For both CP constructs, the chimera results show that the effects on transfer efficiency correspond to the C-terminal portion of the peptides. K_D_ values for the chimeric peptides were determined from single-molecule FRET and ITC titrations with similar results (Supplementary Table 8 and Supplementary Figure 17). The WASH_N_-C1_C_ affinity trends along with CARMIL1 affinity, while C1_N_-WASH_C_ affinity trends along with the one of WASHCAP, suggesting that the C-terminal portion of the peptides largely defines the interaction with CP.

We performed analogous experiments with CP α44β84. The chimera WASH_N_-C1_C_ produced very little change in Ē on binding (Supplementary Figure 18), similar to the result for non-chimeric CARMIL1. In contrast, the chimera C1_N_-WASH_C_ produced a larger change in transfer efficiency, with ΔĒ^[C1N-WC]^ = +0.039 ± 0.001 (Supplementary Table 2). This change is significant, although not as large as that observed for the WASHCAP peptide. These results further support that the C-terminal portions of CPI-motif peptides are responsible for differences in Ē.

We further analyzed CP dynamics related to interaction with the chimeric peptides by comparing the lifetime versus transfer efficiency. In contrast to the effect of CARMIL1, which appeared to restrict conformations of the protein, both chimeric peptides have a minor effect on the range of distance distributions for both CP α44β84 and CP α9β161, relative to Apo-CP. Taken together, our results indicate that the C-terminal region of the chimeras determines the overall register of the distance distribution (mean value), but the whole CPI-motif is required to control both CP dynamics (width of the distribution) and affinities.

To extend the analysis, we compared the experimental FRET results with MD simulation results. MD simulations of the WASHCAP CPI-motif peptide bound to CP showed differences in the dynamics of residues along the length of the peptide. The N-terminal section was much more dynamic, with few specific and/or stable interactions with CP. The C-terminal section was more constrained and maintained more stable interactions with CP (Supplementary Figure 19). The MD results agree with the single-molecule FRET results, supporting the conclusion that the C-terminal portion of the peptide is responsible for the differences in conformational changes and affinities (Figure 7).

## Discussion

We investigated the molecular mechanisms for allosteric regulation of CP, using the complementary approaches of single-molecule FRET experiments and MD simulations. Our study identifies two distinct CP conformations in solution, which are further modulated by two classes of regulators: CPI-motif proteins and V-1. One important discovery is that the conformational changes induced by the two classes of regulators are in opposite directions. This result accounts for the antagonism of the two classes of regulators with respect to their binding to CP. Together, our results provide new evidence and insights into the antagonistic mechanism utilized by CP regulators to control its function.

### CP Conformations in Solution

Previous HDX-MS experiments identified time-dependent solvent exposure of Apo-CP, suggesting that CP may exist in a more dynamic configuration than what has been captured by crystallographic studies ^11^. Here, we provide the first physical evidence for the existence of multiple CP conformations in solution. Both single-molecule FRET measurements and MD simulations agree on the presence of two distinct states (Apo-CP 1 and Apo-CP 2, Figure 8) characterized by a difference in the configuration of the stalk and the cap of the CP heterodimer with respect to each other. From the analysis of the distribution of transfer efficiencies and lifetimes, we conclude that the two states are in slow exchange (longer than milliseconds) with each state characterized by an ensemble of configurations that occur on the nanosecond to microsecond timescale. From the simulations, we gain atomistic details for the corresponding structural features of the two states, representing a torsional change of the stalk propagating to the cap. This new conformational ensemble varies from the structures previously identified for the CP ligand-bound states, suggesting that indeed the Apo-CP conformations identified here are unique to CP in its ligand-free state.

### Effects of Regulators on CP Conformation

CPI-motif proteins and V-1 bind directly to CP at different sites ^9, 10, 13, 27^ and display negative linkage ^6, 7^, that is, the binding of one ligand decreases the affinity of the other ligand. This negative linkage is particularly important because the current model for CP function in cells, as proposed by Hammer and colleagues ^6^, is based on antagonism between the effects of CPI-motif proteins, which are found in small amounts and are restricted to distinct membrane locations ^3^, versus the effects of V-1, which is found in high concentration and diffusing freely in cytoplasm ^5^. A molecular mechanism of linkage is suggested by the observation of changes in solvent accessibility for CP at CPI-motif regulator binding site in HDX-MS analysis when bound by V-1 and vice versa ^11^. Solvent accessibility reflects changes in conformation and/or structural dynamics but lacks specific information about the molecular nature of the differences. Here, we discovered that regulators introduce changes in CP conformations and dynamics in solution and, most important, that the conformational changes induced via interaction with the two classes of regulators occur in opposing directions (Supplementary Figure 5). In particular, we have demonstrated that CP, in solution, adopts distinct conformations when bound to either CPI-motifs (the stalk and β-subunit cap regions move closer together in distance) or when bound to V-1 (the stalk and the β-subunit cap regions move farther apart). Recent work analyzing available crystal structures for CPI-motif-bound CP and V-1-bound CP found similar results (Takeda et al., 2021. PMID 33639213).

The conformational states induced by the binding of V-1 and CPI-motif proteins do not correspond exactly to either of the two states observed for Apo-CP. Our experimental results indicate that binding of V-1 does not restrict CP from exploring the two states; however, the binding of CPI-motif peptides favors a single bound conformation (Figure 8). These findings support the hypothesis that the conformational changes in CP underlie linkage, and they account for antagonism in binding. In this model, the steric inhibition of actin capping produced by V-1 binding is antagonized by CPI-motif binding, and the net effect is activation of CP. Thus, diffuse inhibition becomes spatially limited activation. This model can account for regulation of actin polymerization at barbed ends, either by the promotion of Arp2/3-based polymerization or by the inhibition of formin-controlled polymerization.

### Detailed observations from MD simulations

The stalk conformation for V-1 and WASHCAP structures are very similar (Figure 6) while the conformation of the binding site on CP is very different for these two states (Supplementary Figure 20). Since WASHCAP binds to the stalk, this suggests WASHCAP *could* bind to CP-V-1 and then release V-1 from the complex. Conversely, V-1 *would not* bind as strongly to the CP-WASHCAP complex since the V-1 binding site is in a very different conformation. There is also a significant difference in the V-1-binding site between the Apo-CP 1 and Apo-CP 2 structures (Supplementary Figure 20). The V-1 site on Apo-CP 1 diverges more from the V-1-CP structure, suggesting that V-1 would interact more weakly with Apo-CP 1 than with Apo-CP 2. These findings are consistent with experiments that showed that binding of CARMIL to CP alters the stability of the V-1 binding site as well as changing the accessibility to the binding site by a key tryptophan residue in V-1 ^12^.

The MD simulations revealed substantial alignment of the α-helices of the stalk region along with flattening of the cap region when comparing Apo-CP with both bound states – CPI-motif-bound CP and V-1-bound CP (Figure 6, Movies 2 and 3). Flattening of the cap region has previously been observed in structures of CP bound to V-1 ^9^, to twinfilin-actin ^28^, and to the barbed end of F-actin ^29, 30^. Comparing the two ligand-bound structures from the MD simulations, we observed that the V-1-bound CP structure showed greater twisting of the stalk region of CP, compared to the WASHCAP CPI-motif-bound CP structure, along with slightly more flattening of the cap (Figure 6, Movie 5).

Overall, the biggest conformation differences associated with ligand binding are between the stalk region and the β-subunit cap region. When WASHCAP CPI-motif is bound, the stalk and cap regions move closer together; however, when V-1 is bound, the stalk and the cap regions move farther apart. These findings further demonstrate the linked conformational changes from the stalk through the cap regions, which account for the effects on regulator binding and actin filament capping.

### Differences among CPI-motif peptides: contrasts with biochemical effects

It is interesting to compare the biochemical effects and conformational changes imposed by CPI-motif peptides on CP. While one could expect that strong biochemical effects correlate with large conformational changes, we observed the smallest conformational change for CARMIL1, which has a significant biochemical effect in terms of binding affinity for CP and inhibition of V-1 binding ^7^. This seems to suggest a disconnect between the functional activity of CP and conformational changes induced upon interaction with CPI-motif proteins.

However, the combination of lifetime information and transfer efficiencies indicates that CP adopts a dynamic conformational ensemble where regulators alter both the mean and the width of the distribution of distances. Based on the placement of the dye pairs, one in the cap and one in the stalk, the mean of the distribution reports on the average distance between the two, while the width captures the associated dynamics. With respect to dynamics, the binding of CARMIL1 and WASHCAP restricts the range of CP conformations (CARMIL1>WASHCAP), while interaction with CKIP allows for a broader range compared to Apo-CP (Supplementary Figure 5). The fine-tuned biochemical effects imparted on CP by CPI-motif peptides may be described by the change in both the conformation and dynamics. As suggested by our simulation, altering the mean conformation of CP may result in a different binding surface along the cap, which has a lower affinity for V-1 or actin. At the same time, changes in dynamics may modulate affinity by enabling the exploration of more (or less) binding-competent CP conformations. Recent computational work reported similar findings by exploring distinct modulation of the flexibility of CP whether bound to CARMIL or twinfilin, both of which contain CPI motif ^31^. We speculate that changes in CP conformation and dynamics allow for fine-tuning of function as encoded by the specific sequence of the CPI motif peptides not only in terms of changes in affinity, but for adapting surfaces of interactions to different ligands.

### CPI-motif sequence encodes for conformations, dynamics, and affinities

To better understand how the sequence of CPI-motifs encodes for the register, dynamics, and affinities, we studied two chimera peptides that encompass the N- and C-terminal halves of WASHCAP and CARMIL1. We found that the C-terminal half largely controls the register (mean conformation), while the N-terminal half further modulates for dynamics (range of conformational changes) and affinity. In particular, the binding affinities for WASH_N_-C1_C_ are higher than the one observed for C1_N_-WASH_C_. These observations, derived from our single molecule experiments, are consistent with MD simulations of the WASHCAP peptide where the N-terminal half of the sequence is largely flexible compared to the C-terminal half (Supplementary Figure 19). Our simulation results provide a different perspective on CPI conformations in solution when compared to previous crystal structures of CP with CPI-motif peptide bound ^9^, where the N-terminal half of different peptides can be aligned very precisely (Supplementary Figure 16). We speculate this discrepancy may arise from the specific experimental conditions used to determine the protein structure.

## Conclusions

Our single-molecule experiments and MD simulations revealed the coexistence of two distinct conformational states for CP in absence of ligands, characterized by a torsion of the protein stalk and a change in the configuration of the cap. Interaction with ligands alters both the equilibrium between the two configurations of the protein as well as its dynamics. We found that binding of CPI-motif peptides and V-1 act in “opposite” directions, with V-1 holding CP in an intermediate configuration between the two Apo states and CPI-motif peptides altering the cap curvature and positioning. These observations together provide a plausible explanation of the different biochemical effects of these ligands on CP function. In addition, we discovered that the C-terminal region of CPI-motif peptides dictates the register of binding to the stalk of CP, while the N-terminal region further refines dynamics and binding affinity.

Altogether, our data provides a quantitative description of CP dynamics in absence and presence of ligands and enables bridging and reconciling observations from previous structural studies of the protein.

## Conflict of interests

The authors declare no conflict of interests.

## Supporting information

PDB File 1. CP-Apo 1

PDB File 2. CP-Apo 2

PDB File 3. CP-WASHCAP

PDB File 4. CP-V-1.

## Abbreviations

CP: actin capping protein
Apo-CP: free unbound CP
F-actin: filamentous actin
CPI: capping protein interacting
HDX-MS: hydrogen-deuterium exchange mass spectrometry
FRET: Förster resonance energy transfer
MD: molecular dynamics
ITC: isothermal calorimetry
CARMIL: capping protein Arp2/3 and myosin I linker.

## Acknowledgments

We are grateful to members of our laboratories for advice and assistance. This research was supported by the following NIH grants: GM118171 and GM144082 to J.A.C, AG062837 and AI163142 to A.S., and GM136822 to D.S. NAMD was developed by the Theoretical and Computational Biophysics Group in the Beckman Institute for Advanced Science and Technology at the University of Illinois at Urbana-Champaign.

## Supplementary Text

### Design of Labeling Positions on CP

To monitor in-solution conformational changes via single-molecule FRET, we designed CP full-length constructs that would enable site-specific labeling with maleimide chemistry. As described in Materials and Methods, we designed, expressed, and purified a cysteine-less variant of CP (Cys-null CP). The activity of Cys-null CP was similar to that of wild-type CP in assays for capping F-actin barbed ends and for binding to a CPI-motif peptide (Supplementary Figure 1). Also, ITC measurements of the binding affinity of wild-type CP and Cys-null CP with the CPI-motif peptide from CARMIL1 showed no significant differences (Supplementary Table 1).

Next, we identified optimal fluorophore labeling positions. Crystal structures of Apo-CP and ligand-bound CP display rather small changes in structure despite the substantial biochemical changes. We investigated whether larger conformational changes occur in solution upon binding. To identify labeling position candidates, we began by comparing available crystal structures of free CP (Apo-CP, PDB 1IZN) with those of V-1-CP complex (PDB 3AAA) and CARMIL1-CP complex (PDB 3LK3) ^9, 13, 16^ to identify locations with relatively large differences in structure. We analyzed the (C_α_ - C_α_) distance between each pair of residues in the structures, and we computed corresponding transfer efficiencies for Alexa 488 and 594 donor-acceptor dyes (Förster radius R_0_ = 5.4 nm). Pairs of labeling positions were ranked based on the transfer efficiencies and the differential between Apo-CP and ligand-bound structures. The ranking criteria ensured that distances were in the dynamic range measurable by FRET and that the differences between Apo-CP and ligand-bound structures were sufficient to provide a measurable change in transfer efficiency. For each pair of dye positions, we also evaluated the proximity of intrinsic Tryptophan residues, which can cause quenching of the dyes and alter transfer efficiency ^32, 33^. In addition, dye positions were chosen to avoid disrupting secondary and tertiary structure when mutating the residue to a cysteine.

Based on these considerations, we identified the two sets of labeling positions: CP α1N44C β2E84C (“α44β84”) and CP α1S9C β2S161C (“α9β161”) (Figure 1). We labeled the CP samples with a sequential labeling procedure ^15, 33^. Substoichiometric dye concentrations were used for donor dye coupling, followed by purification of single-donor-labeled CP, and then coupling acceptor dye with a molar excess of dye.

We analyzed the labeled constructs by mass spectrometry to verify protein integrity and correct labeling position. Mass spectrometry revealed that the two fluorophores selectively labeled specific Cys residues in the two variants. For α44β84, nearly all acceptor dye was coupled to α subunit residue 44, and nearly all donor dye was coupled to β subunit residue 84 (Figure 1 and Supplementary Figure 2). For α9β161, β161 was predominantly labeled with donor dye, and α9 had more acceptor dye than donor dye, at a ratio of ∼ 2:1 (Figure 1 and Supplementary Figure 2). Based on these results, we expected energy transfer for both constructs to occur between donor fluorophore at one location, predominantly or exclusively the β subunit, and acceptor fluorophore at the other location, predominantly or exclusively the α subunit.

### Determination of dye-linker distributions

In order to describe the contribution of dye linker distribution, we generated interdye distance distributions at each labeling position using the python package FRETraj (https://rna-fretools.github.io/fretraj/intro.html) ^23^ and the crystal structure of the protein. Each distribution of distances was then fitted to a block copolymer model containing a rigid rod of length 𝑋 (representing the rigid distance within the protein) flanked by two ideal flexible polymers of root mean square radius 𝐿 (here representing the dye-linkers, but in general representing any flexibility in the protein beyond the rigid distance) ^34^:

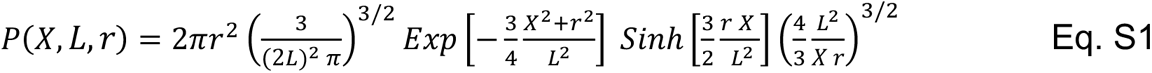

### Contribution of dye-linker dynamics to fluorescence lifetime

The fluorescence lifetime is related to the mean transfer efficiency ^35^ through:

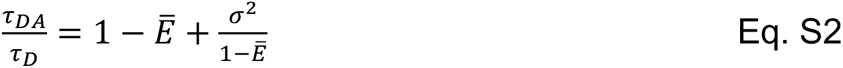

where 𝜎, is related to the variance of the sampled distribution *via*:

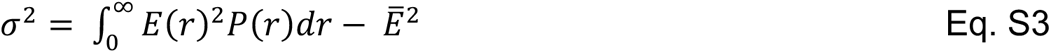

For 𝜎, equal to zero, Eq. S2 reduces to the linear trend expected for a rigid distance.

Eq. S1 was used to generate a 𝑃(𝑟) that is a function of the rigid rod length 𝑋 and of symmetric flexible regions of root-mean-square distance 𝐿, 𝑃(𝑋, 𝐿, 𝑟). Eqs. S2 and S3 were used to fit the dependence of the fluorescence lifetime vs the mean transfer efficiency and obtain estimates of the distribution underlying the measured transfer efficiency, using 𝑋 and 𝐿 as free fitting parameters. For comparison, results were contrasted with the ones expected for a completely flexible polymer:

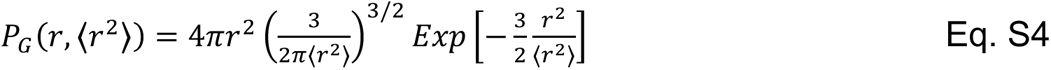

Finally, we subtracted the contribution of the dye-linkers (as estimated from the dye-clouds, see previous paragraph) from the fitted dynamics and estimate the excess dynamics (not due to the dye-linker).

### Binding models

Our single-molecule data revealed the coexistence of two populations in equilibrium. Here, we discuss three models we considered to analyze the experimental data.

As a first model, we consider a simple two-state model with a bound and unbound state in equilibrium. In this case the equilibrium constant can be written as:

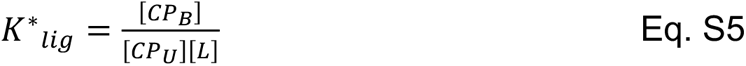

where 𝐾*_lig_* is the equilibrium constant, [𝐶𝑃*_U_*] is the free concentration of unbound CP, [𝐶𝑃*_B_*] is the concentration of bound CP, and [𝐿] is the free concentration of ligand.

When fitting the change in the mean transfer efficiency 𝛥𝐸(𝐿) as a function of the ligand concentration [𝐿], we use Eq. S6 to derive:

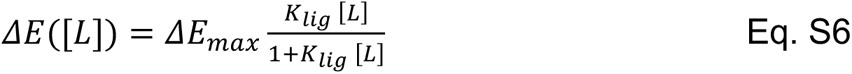

where 𝛥𝐸*_max_* represents the maximum excursion in the variation of transfer efficiency.

As a second model, we considered the scheme in Supplementary Figure 10, with three states in equilibrium, Apo-CP_1_, Apo-CP_2_, and bound CP. In this model the fraction associated to each state can be written as:

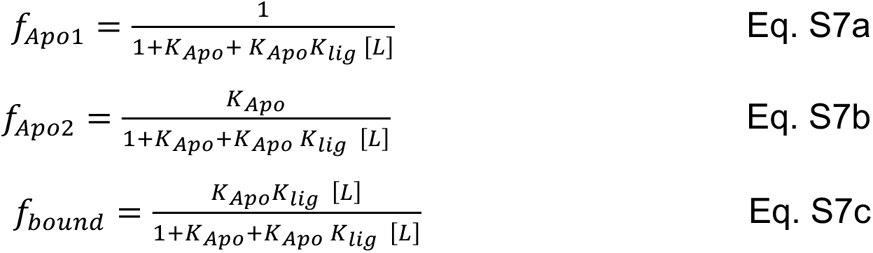

where 𝐾*_Apo_* represents the equilibrium constant between the two Apo configurations and 𝐾*_lig_* represents the equilibrium constant between Apo-CP_2_ and the bound configuration. It is important to note that 𝐾*_Apo_* is not dependent of the ligand concentration [𝐿] and the binding term in Eq. S7c can be reduced to the one in Eq. S6 by a simple mathematical manipulation.

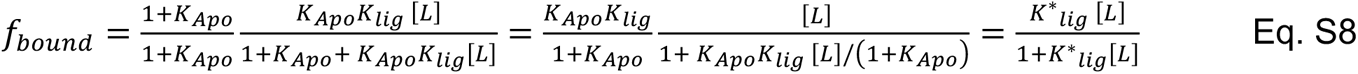

with

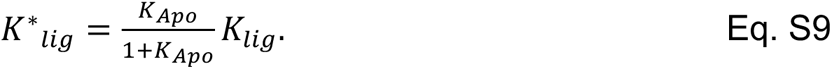

In other words, the protein binds with equilibrium constant 𝐾^∗^*_lig_*, and the Apo states are distributed according to the relative fractions determined by 𝐾*_Apo_*, 1/(1+𝐾*_Apo_*) and 𝐾*_Apo_*/(1+𝐾*_Apo_*). Eqs. S7 are used to fit single-molecule data, and Eq. S9 is used to compute the equilibrium constant (or dissociation constant) for comparison with ITC measurements. The advantage of defining 𝐾*_lig_* is that it enables a direct comparison with the equilibrium constants derived in the third model.

This second model assumes that the equilibrium between the two Apo configurations is maintained upon binding. Careful inspection of bound histograms for all ligands reveals that this is not the case. Therefore, we considered a third model that contains four states: the two Apo configurations and two bound states (see Supplementary Figures 9,12,13).

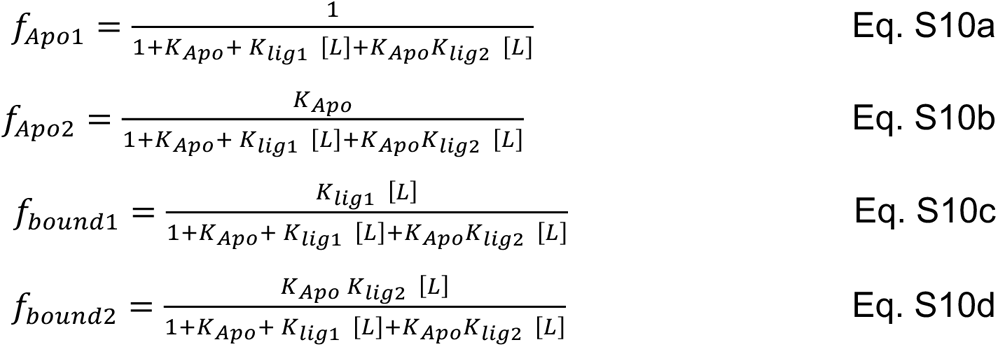

Analogously to the 3-state case, we computed an effective 𝐾^∗^*_lig_* based on the variables of the 4-state model. We started from the total fraction in the bound 

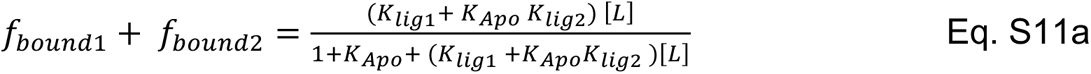

and applied the same calculation of Eq. S9:

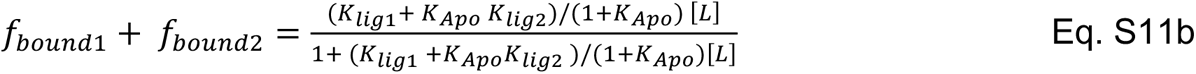

From Eq. S11b, we can write

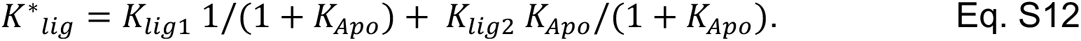

For V-1, it was possible to fit four distinct populations based on the experimental observations. However, for CPI-motif peptides, we needed to combine (sum) Eqs. S9a and S9c to describe the changes in the population at low transfer efficiency. Different from the second model, this model allows for modeling the changes in the equilibrium constant observed between Apo and bound states of CP. However, given the preponderance of one population over the other, the contribution to the overall binding affinity of the low affinity state is largely negligible and the major population binds with similar affinities as those recovered by the second model.

Instead of association constants, the text reports corresponding dissociation constants, computed as 𝐾*_D_* = 1/𝐾.

### Possible differences between single-molecule data and simulations

Discrepancies between single-molecule and simulation data are expected for various reasons, due to current limitations in the quantitative comparison of the two approaches. Occurrence of two populations in the Apo configuration of CP is observed for only one distance pair in single-molecule experiments, compared to expectations from MD simulations. Since the timescale of interconversion between the two states appears to be longer than the burst duration, this implies a discrepancy in the transfer efficiency values determined in experiments and estimated from simulations for CP α44β84. A similar discrepancy occurs for construct CP α9β161. It is useful to discuss the possible factors at the origin of these discrepancies, which are in line with previous observations ^15, 36^.

First, dyes have been added post-hoc on the simulation, accounting only for the excluded volume of the molecules, but not for their charge and hydrophobicity. While this approach allows testing various labeling positions without performing independent simulations for each labeling construct, the simplicity of the method may not capture the actual configurations of the protein with dyes as well as the range of angles explored by the fluorophores.

In addition, transfer efficiencies are computed assuming a specific Förster radius, which is influenced by many factors, including quantum yield and κ^2^. Experimentally, no significant change in photon emission and lifetime was observed when comparing the two labeled constructs, indicating a similar quantum yield. While κ^2^ is commonly assumed to be equal to 2/3, a whole range of values is compatible with the experimental constraints provided by fluorescence anisotropy (see Supplementary Table 7). Distribution of angles determined from the simulated cloud of dyes suggests that the angle restrictions may favor specific values of κ^2^ that depart from the mean of the distribution. Such a correction can contribute to reducing the observed discrepancy.

Finally, static quenching and linker dynamics may contribute to alter the value of the mean transfer efficiency. It is important to note that static quenching of the acceptor would result in lower transfer efficiencies: while this can possibly explain the overlap between populations for CP α44β84, this explanation does not explain the trend observed for CP α9β161.

## Supplementary Figures

**Supplementary Figure 1.**
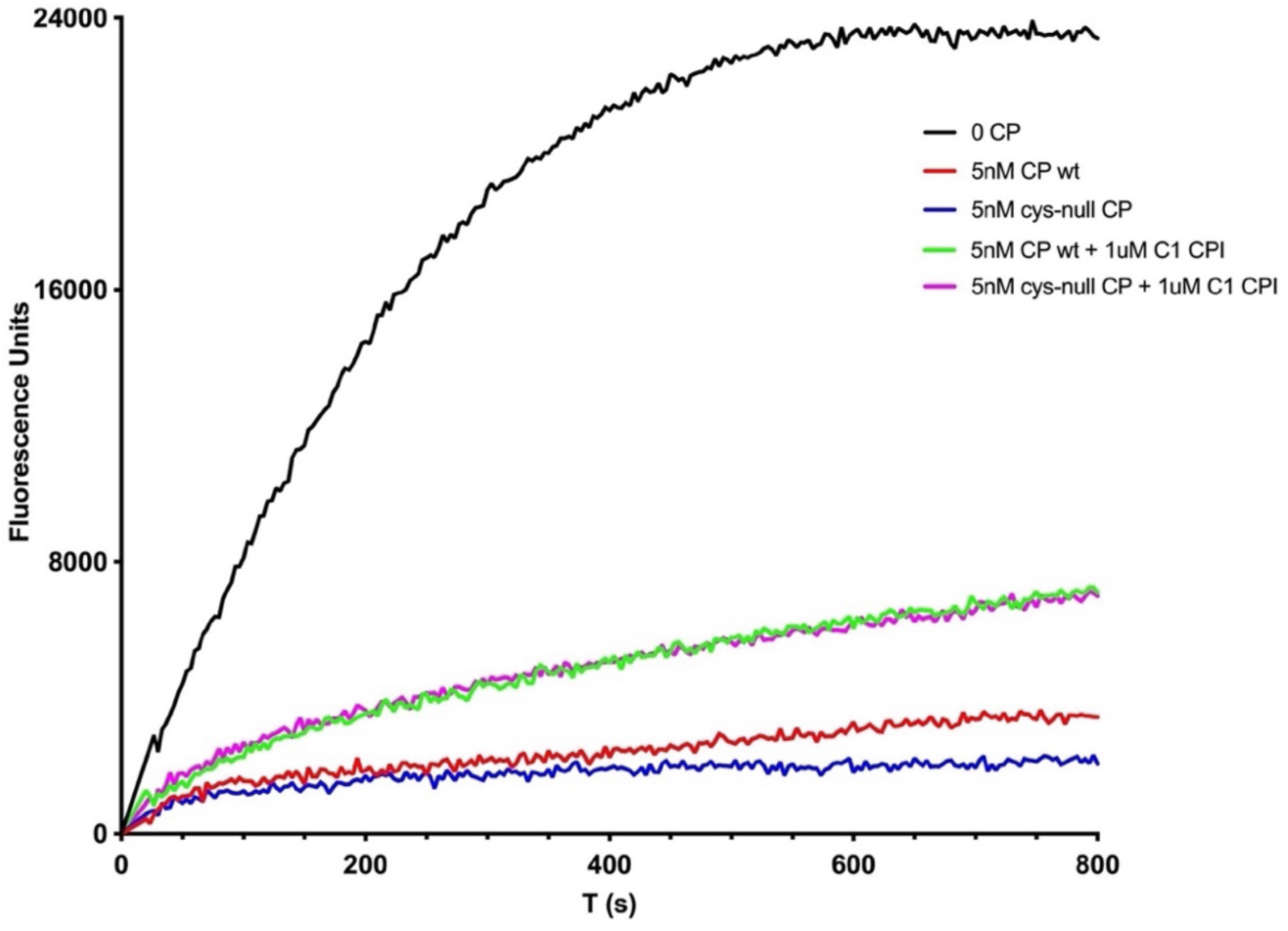
Pyrene-actin polymerization assay for barbed-end capping comparing wild-type CP with Cys-null CP. “C1 CPI” is the CPI-motif fragment of CARMIL1. The assay shows no difference in activity.

**Supplementary Figure 2.**
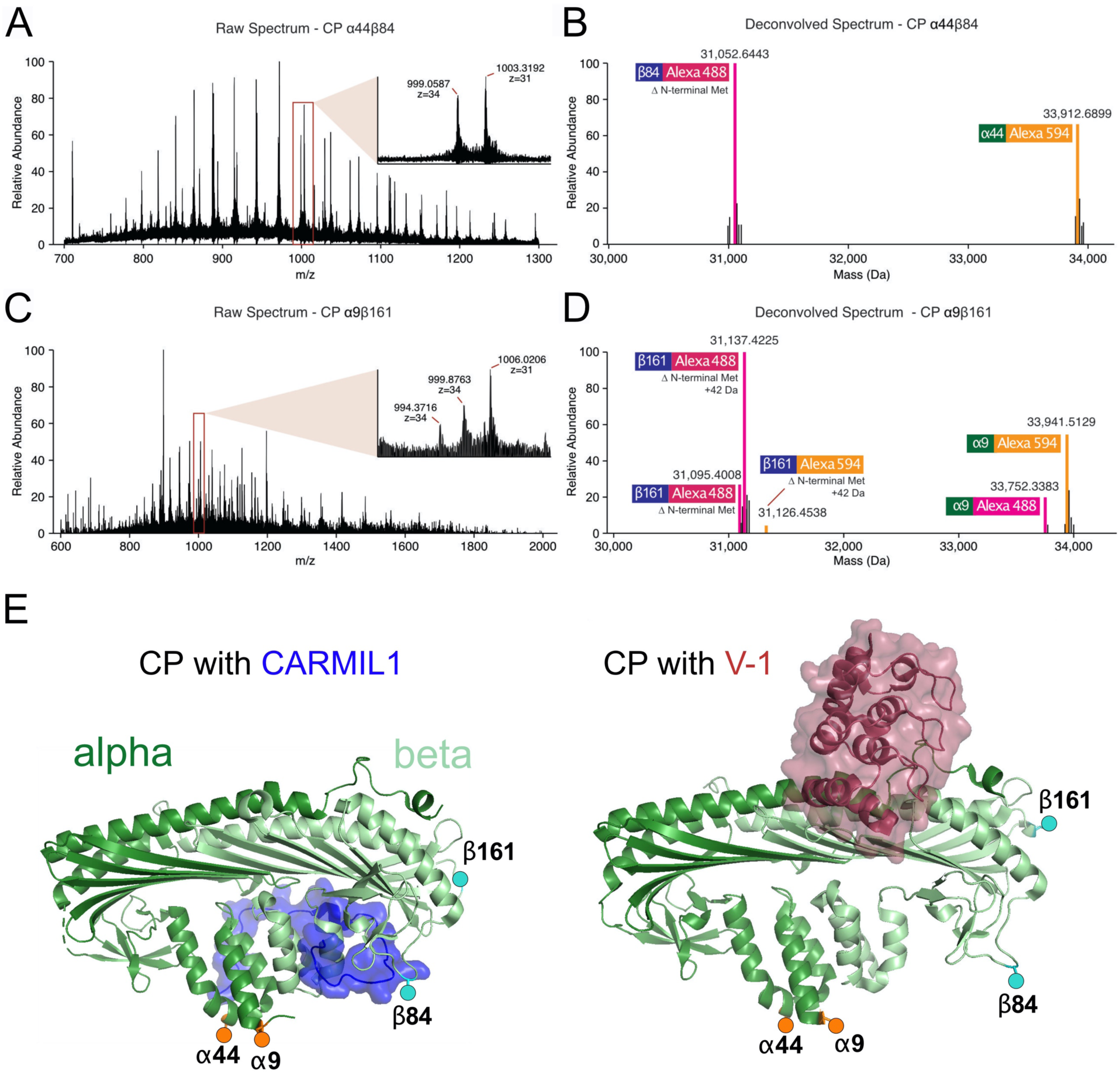
Mass Spectrometry analysis of FRET labeled constructs. **A.** Raw spectrum (100 scan average) of construct CP α44β84. Inset: zoom-in of isotopically-resolved peaks. Peaks corresponding to the most abundant isotopes are labeled. **B.** Deconvoluted spectrum of **A**. Peaks are annotated if they match the expected monoisotopic mass of the α or β subunit conjugated with either the donor dye (Alexa 488) or acceptor dye (Alexa 594). Common mass shifts and ±1 Da deconvolution errors were considered. The β subunit lacks the mass of an N-terminal methionine. **C.** Raw spectrum (100 scan average) of construct CP α9β161. Inset: zoom-in of isotopically-resolved peaks. Peaks corresponding to the most abundant isotopes are labeled. **D.** Deconvoluted spectrum of **C**. Peaks are annotated if they match the expected monoisotopic mass of the α or β subunit conjugated with either the donor dye (Alexa 488) or acceptor dye (Alexa 594). Common mass shifts and ±1 Da deconvolution errors were considered. The β subunit mass shift corresponds to an additional +42 Da following a loss of an N-terminal methionine. **E.** Illustration of dye positions, using previously published structures of CP-ligand complexes (PDB 3LK3 and 3AAA). Donor dye positions are teal circles, and acceptor dye positions are orange circles.

**Supplementary Figure 3.**
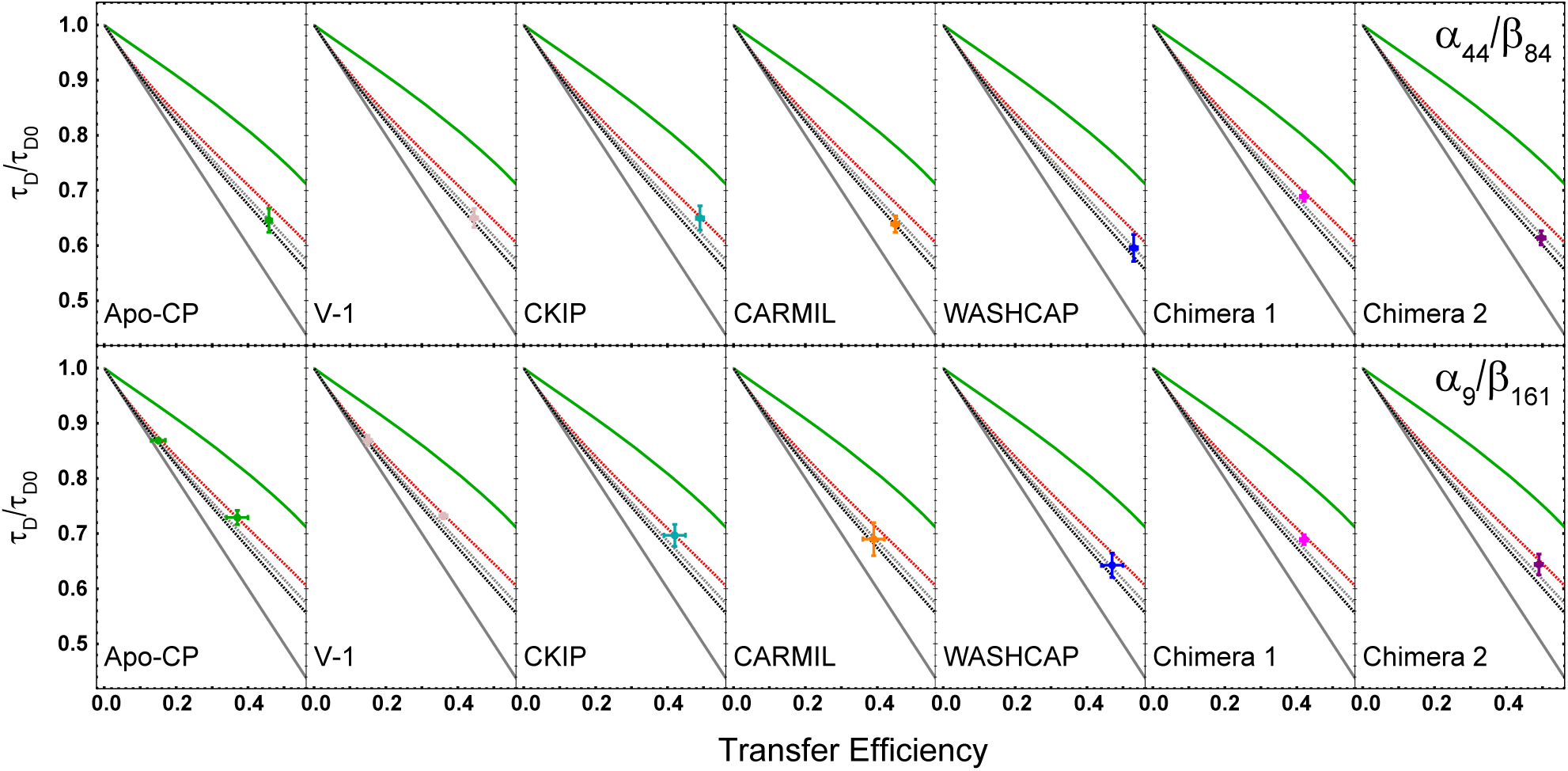
Lifetime vs transfer efficiency plots for Apo-CP and ligand-bound measurements with CP α44β84 (upper panels) and CP α9β161 (lower panels). Each panel includes one (or two) values for experimental data, plotted as a cross (i.e., a dot with vertical and horizontal error bars). Cross values are computed from the mean value and standard deviation of the associated datapoints as obtained by multiple independent measurements. The color of the cross indicates the nature of the ligand, corresponding to the color scheme used throughout. Two experimental values are plotted in the Apo-CP and V-1 panels for α9β161, because the data include two populations. In every panel, all the lines (solid and dotted) are the same and serve as references for the eye of the reader. The two solid lines are the extremes: the gray solid line represents the static case with no linker-dye dynamics; and the green solid line represents the case for a rapidly reconfiguring disordered region (i.e., a Gaussian distribution). Dotted lines correspond to fittings of experimental data: Gray and red dotted lines represent static configurations of Apo-CP with contributions from linker-dye dynamics (gray is CP α44β84 and red is CP α9β161). Black dotted line is CARMIL1-bound; and it corresponds to the least dynamic state observed in the experiments.

**Supplementary Figure 4.**
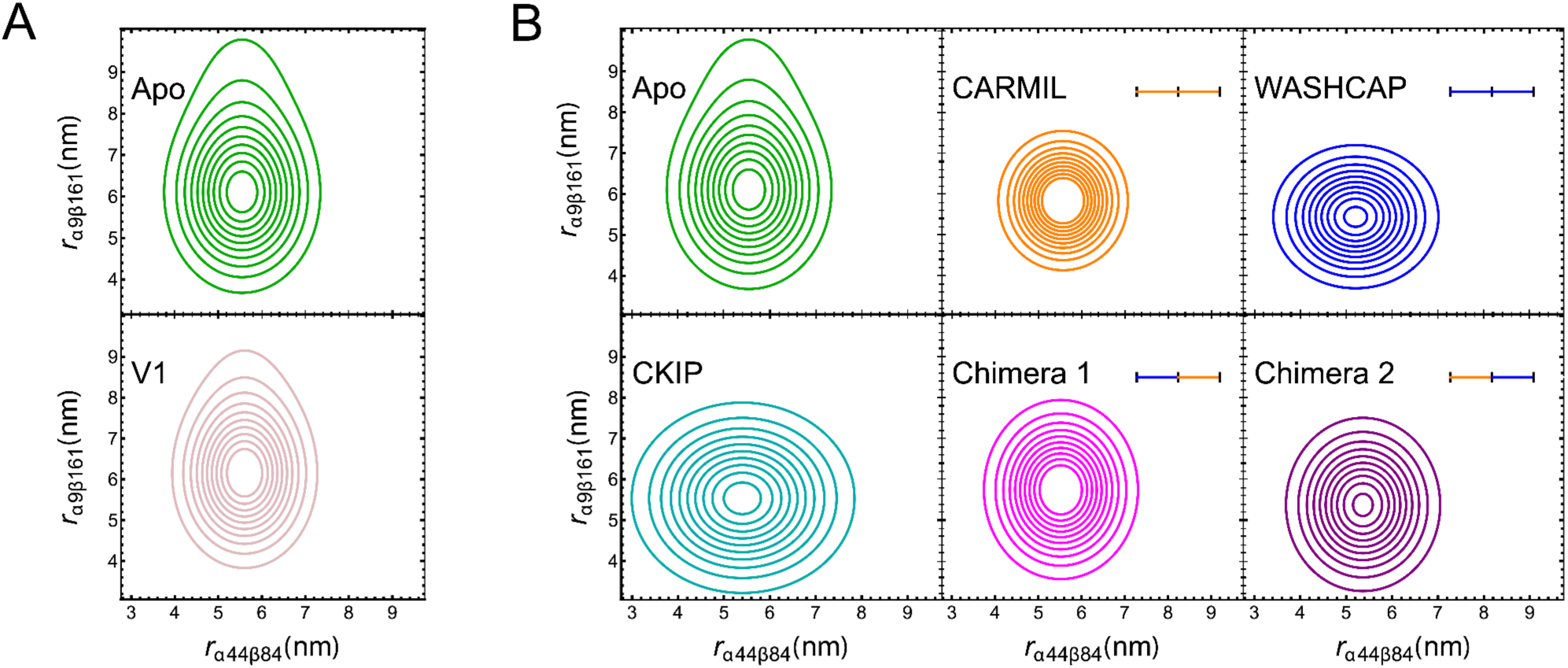
Distance distributions for Apo-CP and ligand-bound CP based on the lifetime analysis. Apo (green) represents the combination of distance distributions sampled by Apo-CP 1 and Apo-CP 2. Binding of V-1 (pink), CARMIL1 (orange), WASHCAP (blue), CKIP (cyan), Chimera 1 (WASH_N_-C1_C_, fuchsia) and Chimera 2 (C1_N_-WASH_C_, purple) induce modifications in the distance distribution.

**Supplementary Figure 5.**
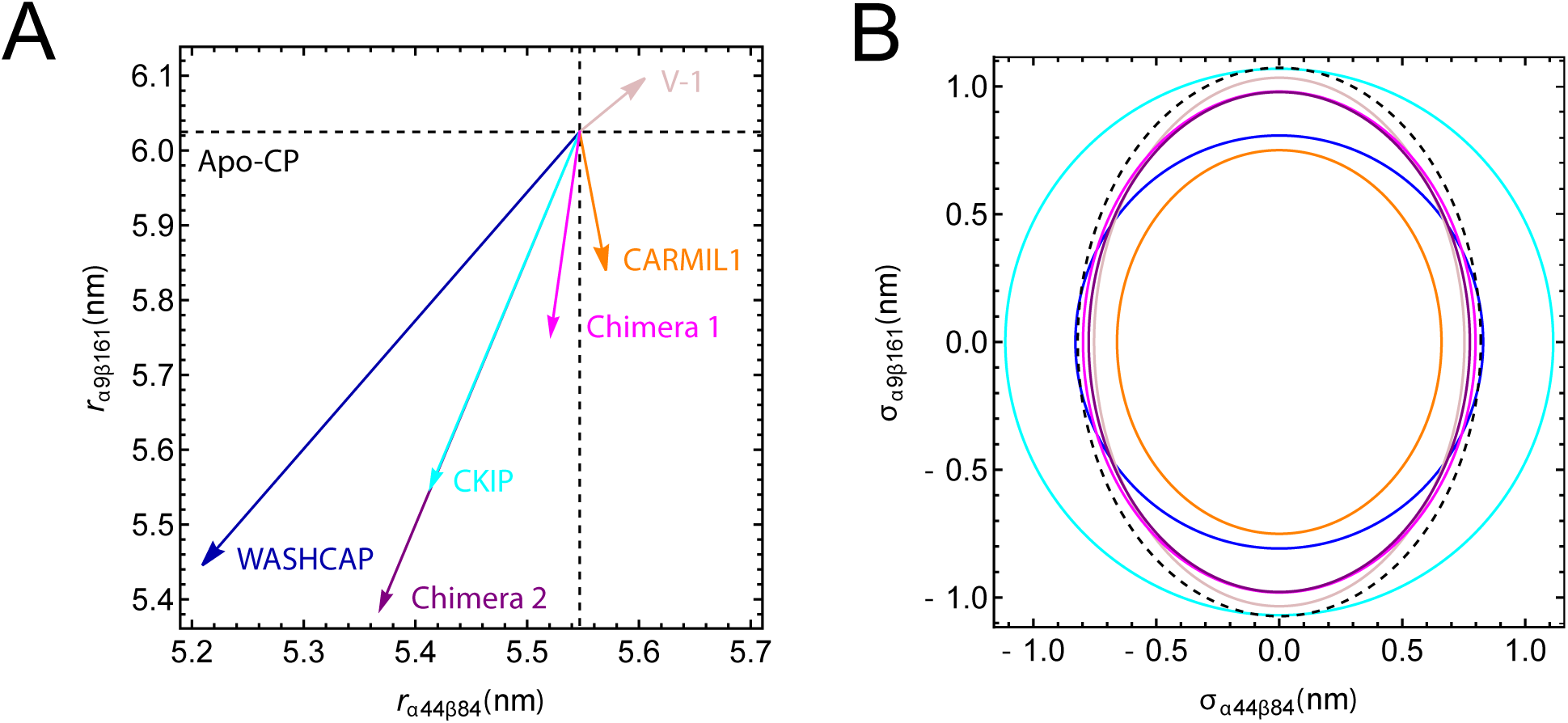
Comparison of conformational changes in the distributions of ligand-bound and Apo-CP conformations. **A.** Arrows reflect changes in the mean distance of the conformational distributions. Black dashed lines refer to Apo-CP 2 conformations, which are used as a reference for comparison with ligand-bound states. **B.** Standard deviations of the conformational distribution for the different ligands. Black dashed lines refer to Apo-CP.

**Supplementary Figure 6.**
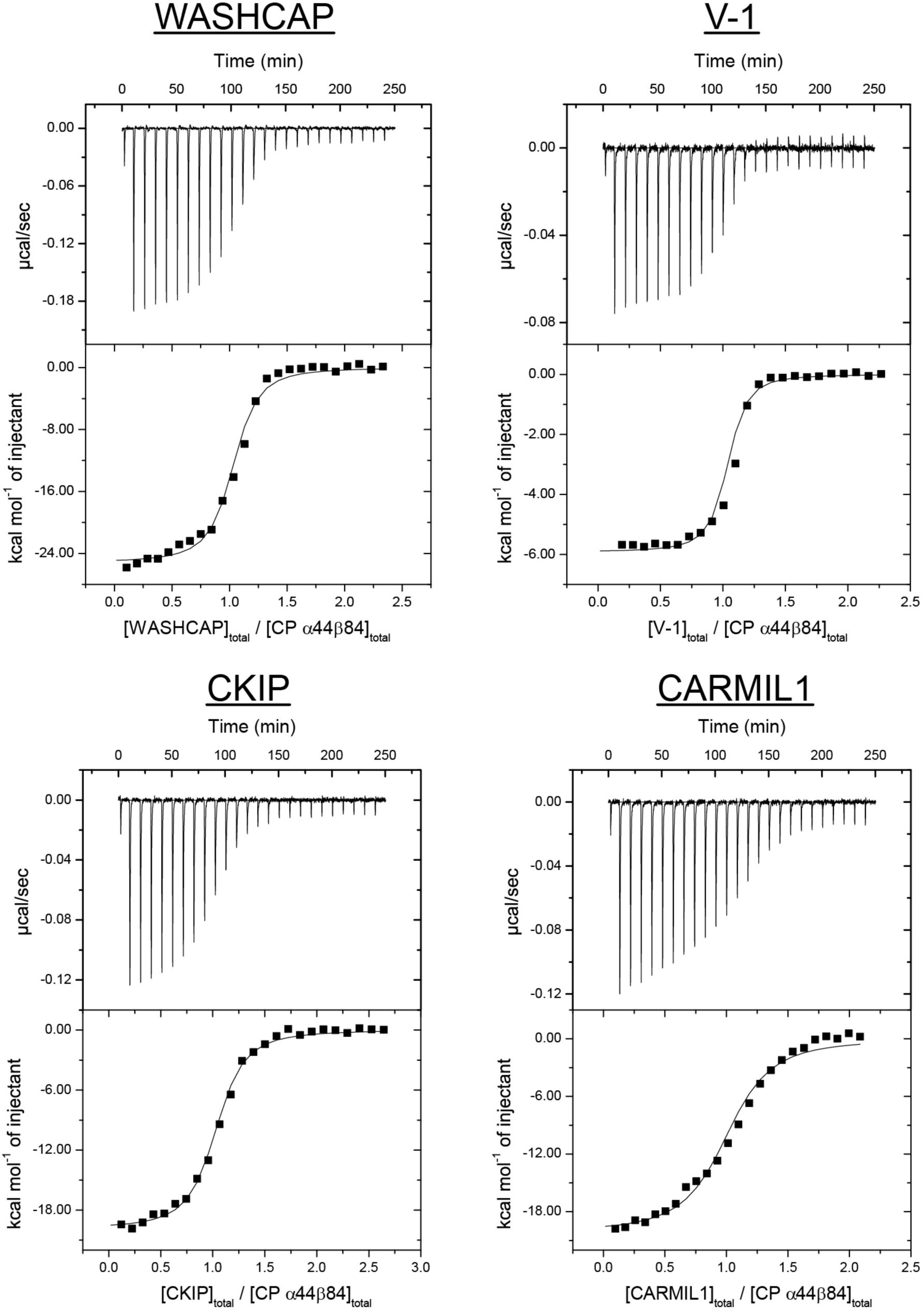
ITC data and binding titration fits for CP with WASHCAP, CKIP, and CARMIL1 CPI-motif peptides and with V-1.

**Supplementary Figure 7.**
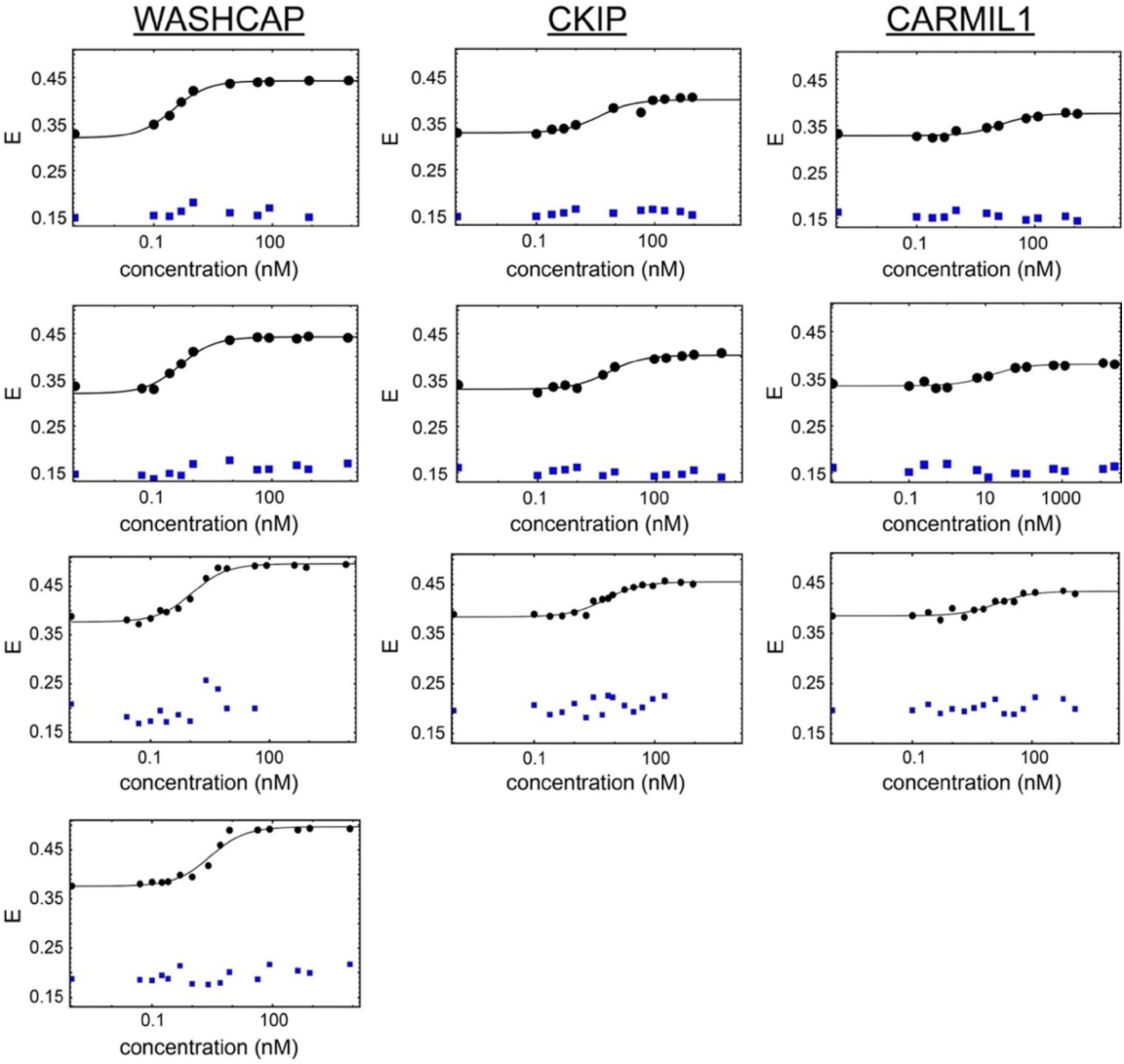
Titration curves for CP α9β161 with CPI-motif peptides. Plots of transfer efficiency (Ē) versus concentration of CPI-motif peptide for each individual experiment. E_1_ (blue) and E_2_ (black) values are shown.

**Supplementary Figure 8.**
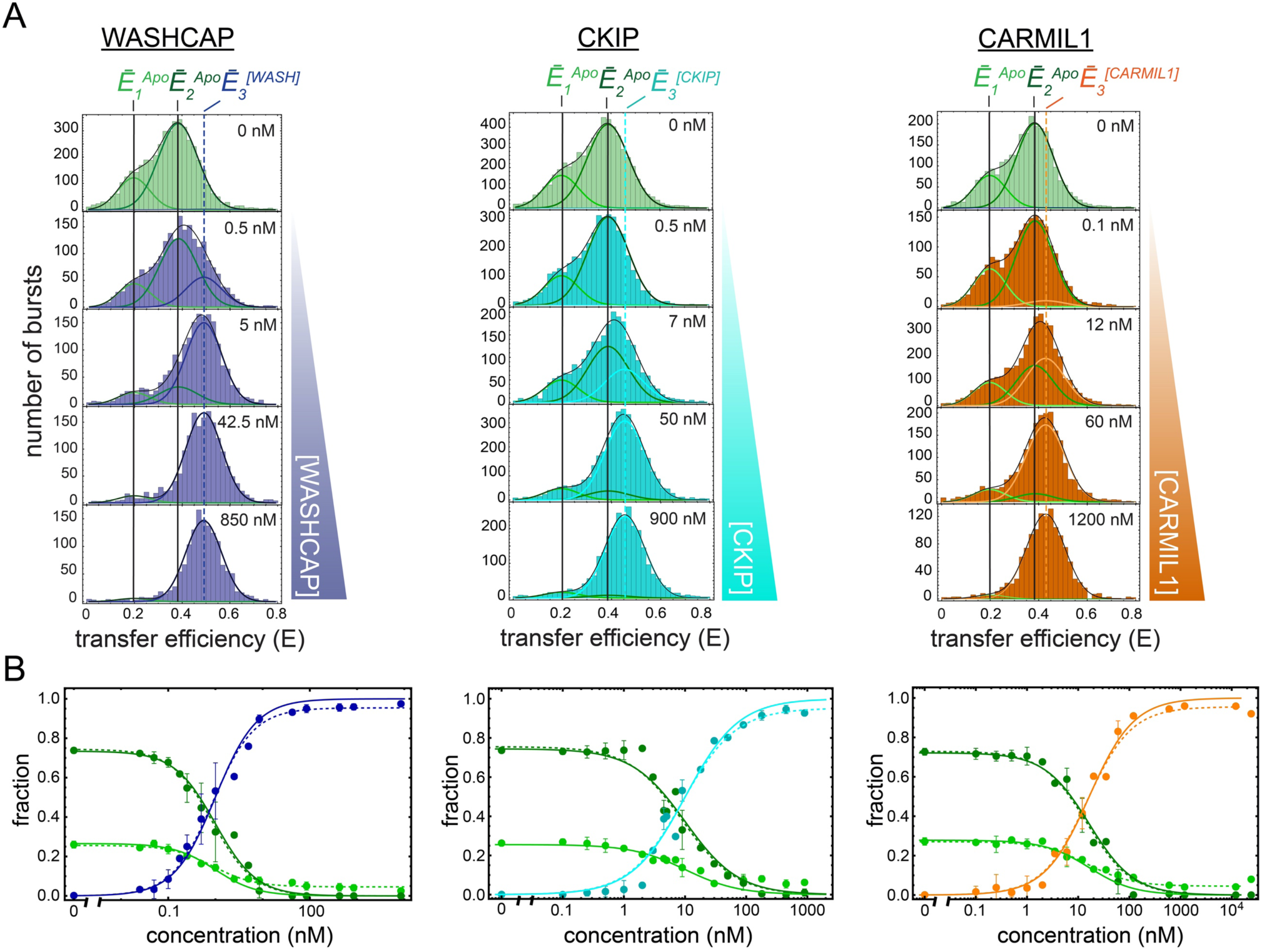
Titration plots for CP α9β161. A. The area under each curve (states with mean transfer efficiencies 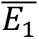 and 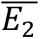 and one bound state with mean transfer efficiency 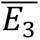 was calculated for each ligand concentration. B. The titration curves plot the fraction of area under the curve for the unbound and bound populations versus the concentration of WASHCAP (blue), CKIP (cyan), or CARMIL1 (orange) CPI-motif peptide. Titrations were performed in triplicate, with each data point plotted as mean and standard deviation.

**Supplementary Figure 9.**
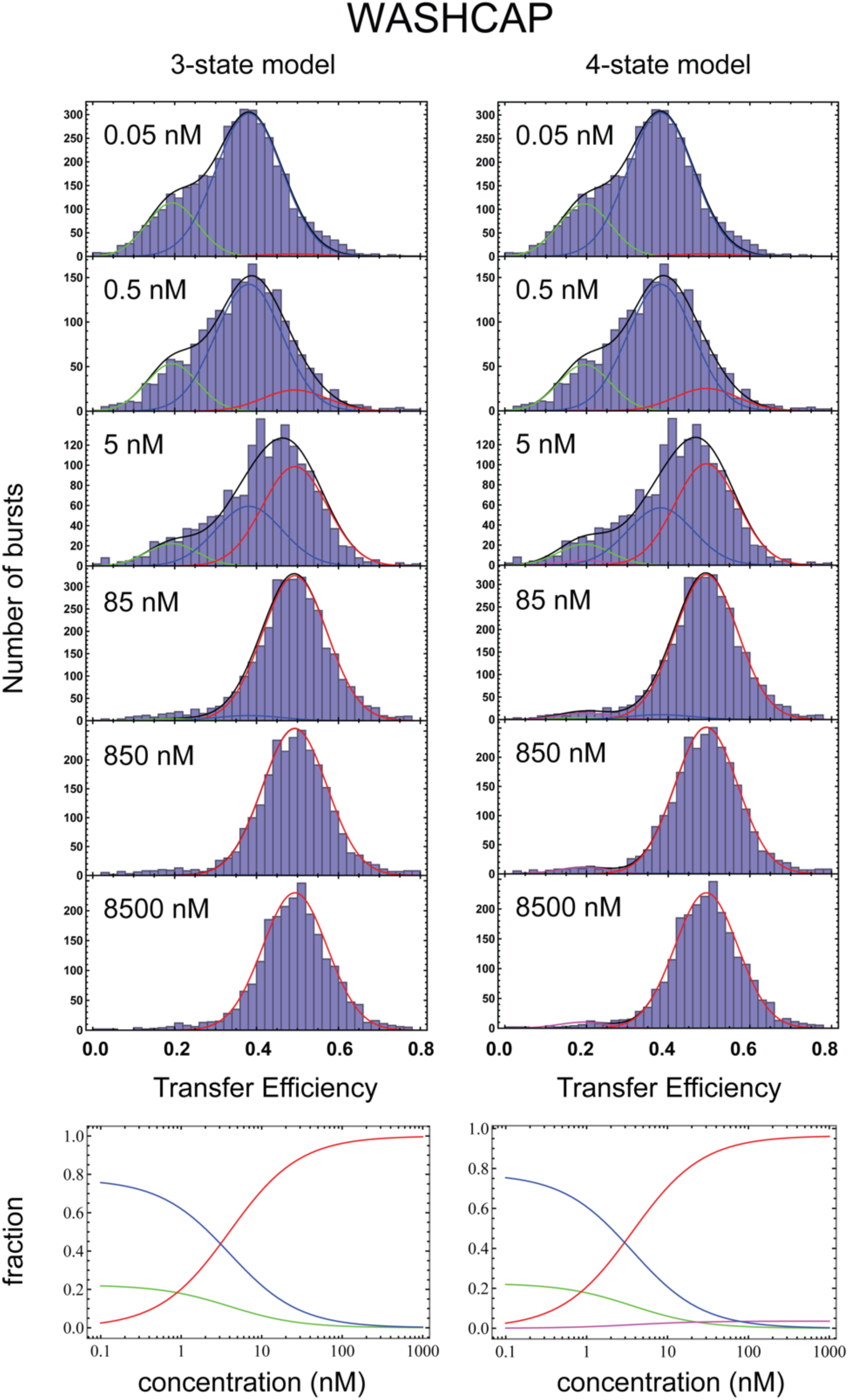
Subpopulation from the analysis of WASHCAP binding for CP α9β161 with global fit of the transfer efficiency distribution using a 3-state or 4-state (red) model. *Top panels.* We analyzed the binding of each ligand by fitting to a 3-state model or a 4-state model, according to Eq. S7-S10. For WASHCAP, in the 4-state model we assigned four distinct mean transfer efficiencies. The three states are described in blue, green, and red. The four states are described in blue, magenta, red, and green. *Lower panels.* Comparison of the estimated fractions of each state across the concentrations for the 3-state and 4-state models.

**Supplementary Figure 10.**
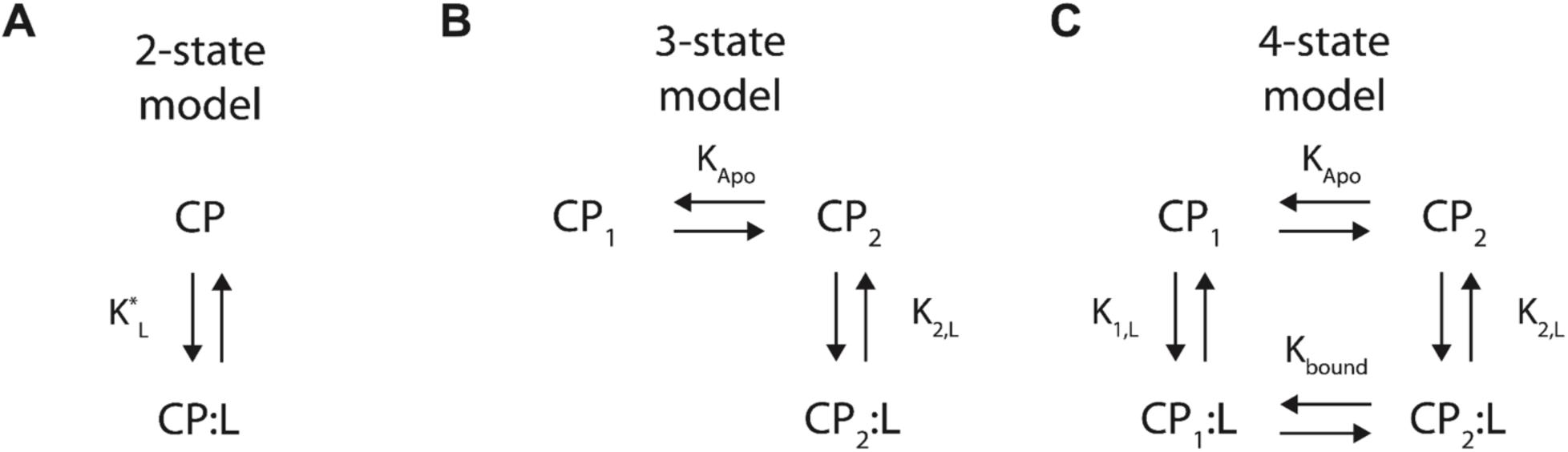
Equilibrium models used to analyze data in ITC and single-molecule FRET experiments. L is ligand, CP_1_ is Apo-CP 1, and CP_2_ is Apo-CP 2.

**Supplementary Figure 11.**
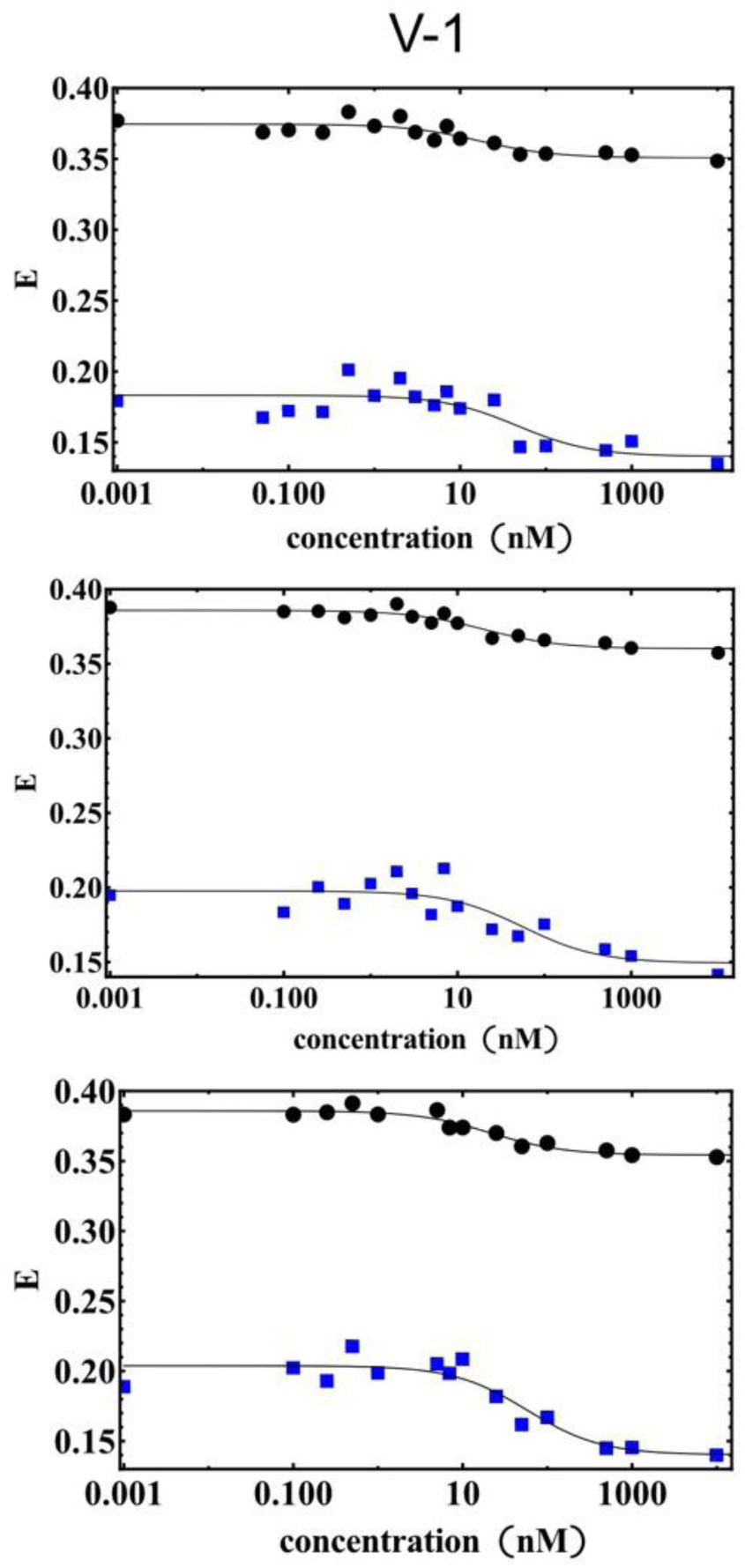
Titration curves for CP α9β161 with V-1. Plots of transfer efficiency (Ē) versus concentration of V-1 for each of three individual experiments. E_1_ (blue) and E_2_ (black) values are shown.

**Supplementary Figure 12.**
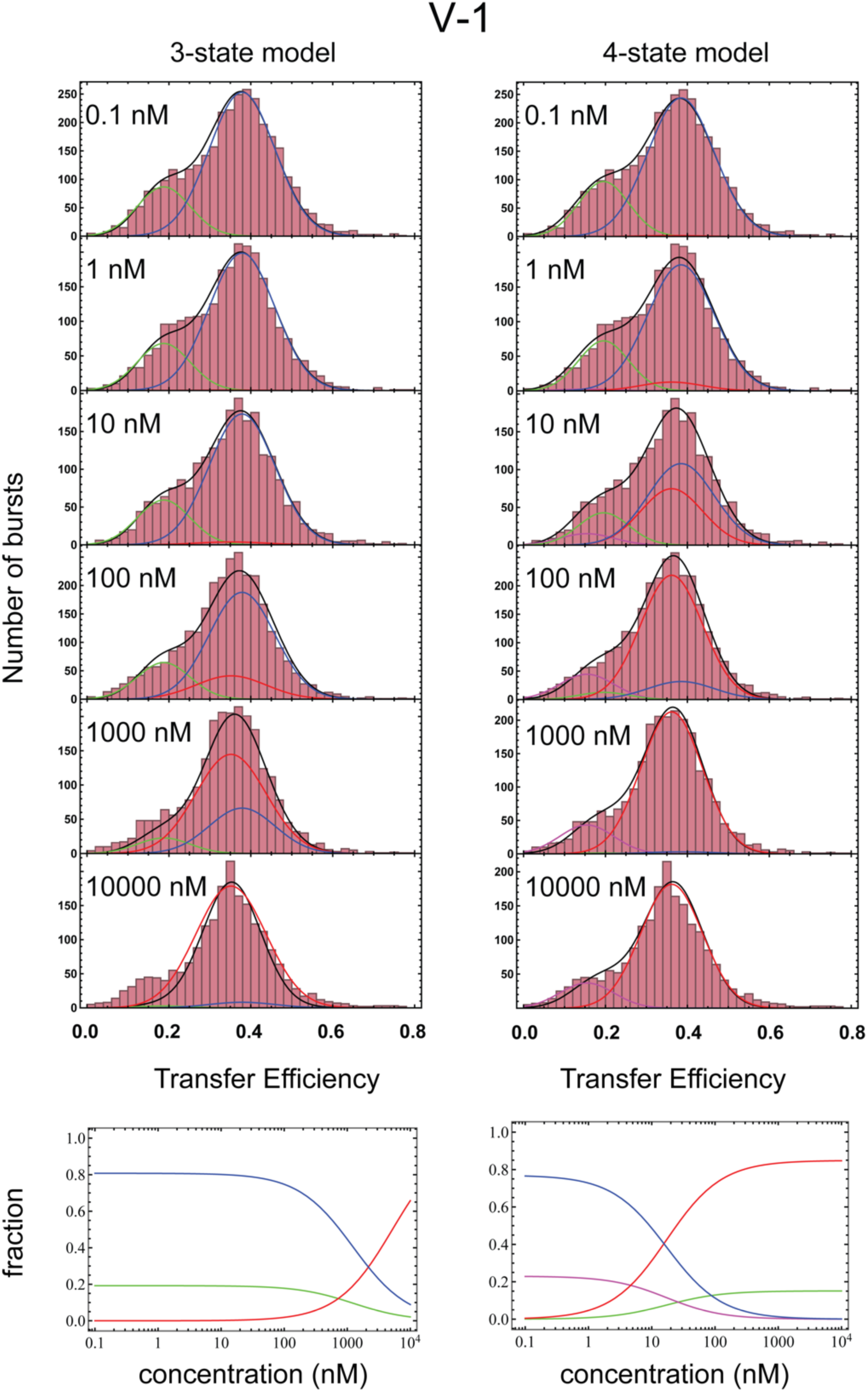
Subpopulations from the analysis of V-1 binding to CP α9β161 with global fit of the transfer efficiency distribution using a 3-state or 4-state (red) model. *Upper panels.* We analyzed the binding of each ligand by fitting to a 3-state or 4-state model, according to Eq. S7-S10. For V-1, in the 4-state model we assigned four distinct mean transfer efficiencies. The three states are described in blue, green, and red. The four states are described in blue, magenta, red, and green. *Lower panels.* Comparison of the estimated fractions of each state across the concentrations for the 3-state and 4-state models. The 3-state model fails to capture the trend of the data.

**Supplementary Figure 13.**
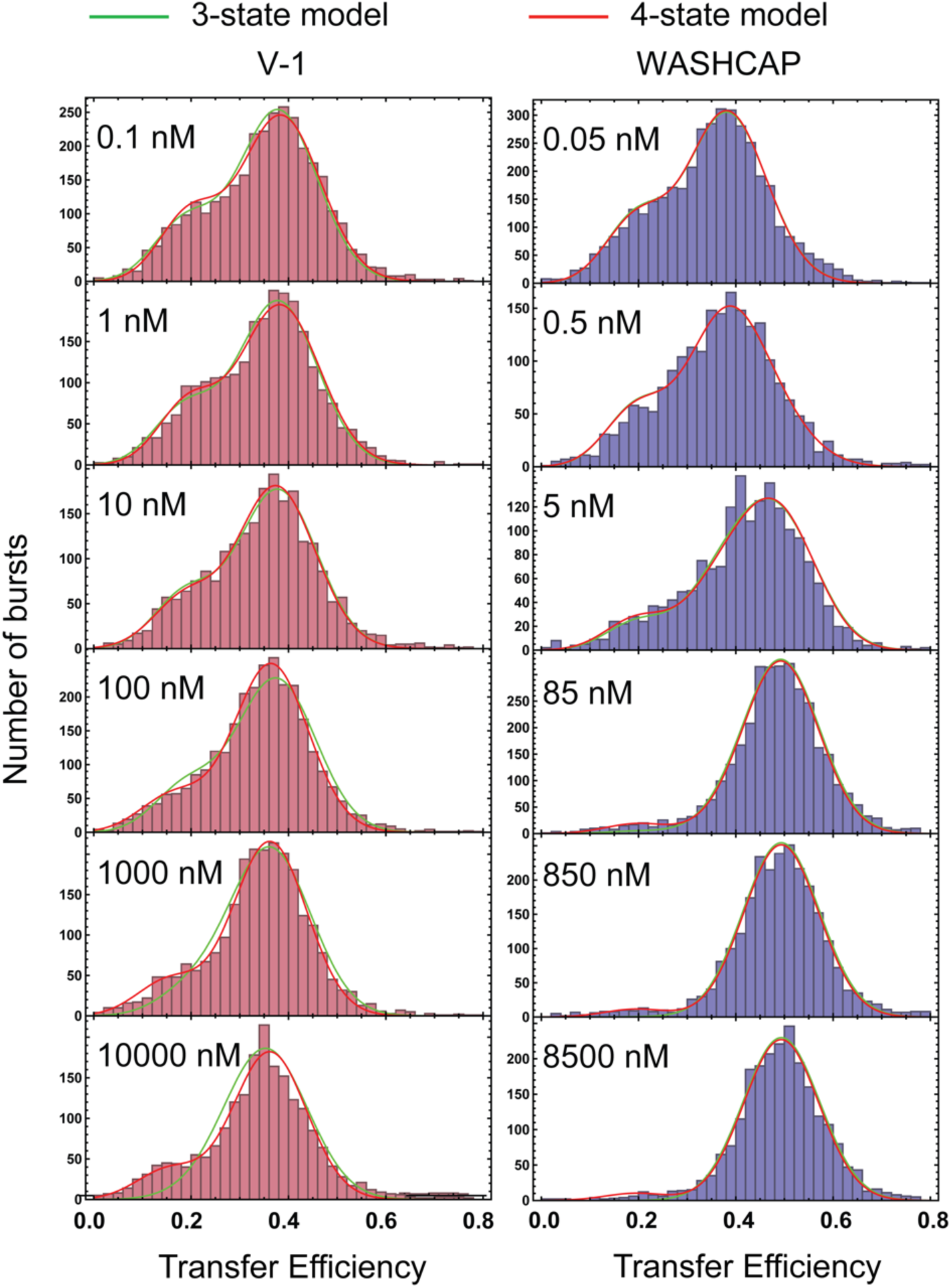
Analysis of V-1 and WASHCAP binding to CP α9β161 with global fit of the transfer efficiency distribution using a 3-state (green) or 4-state (red) model. We analyzed the binding of each ligand by fitting to a 3-state or 4-state model, according to Eq. S7-S10. For V-1, in the 4-state model we assigned four distinct mean transfer efficiencies. For WASHCAP, in the 4-state model we assumed the mean transfer efficiency of the Apo1 population is equal to the one of Bound1 (since the amplitude is too small to properly define a new mean transfer efficiency). The 4-state model better reproduces the data of V-1.

**Supplementary Figure 14.**
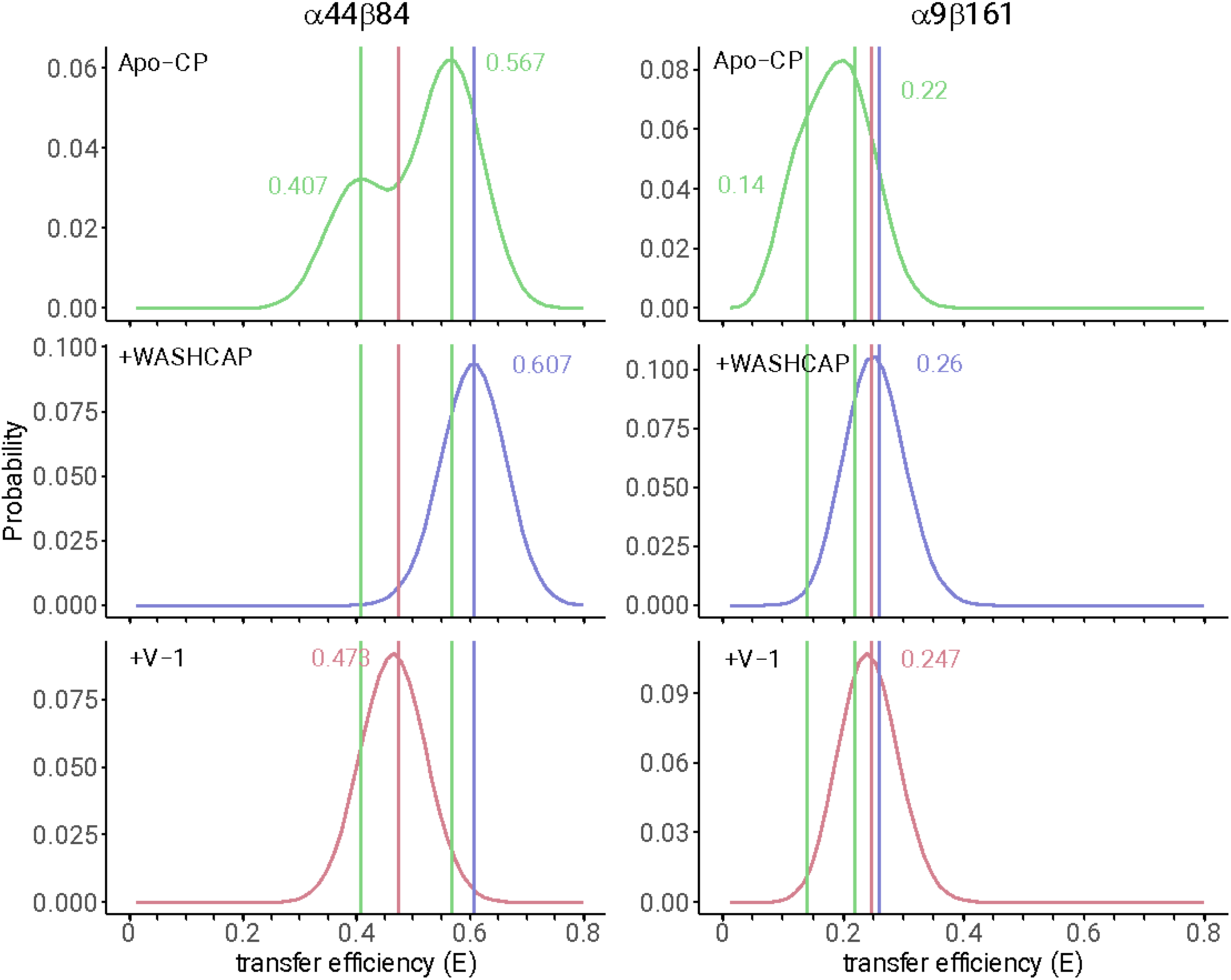
Distributions of transfer efficiencies calculated from MD simulations. Each distribution represents shot-noise limited peaks (from an average of 75 photons) centered in the mean of the transfer efficiency distributions calculated from the full MD trajectory for CP α44β84 and CP α9β161. Data for Apo-CP is in green, and the mean values of transfer efficiency are shown for each basin in the free energy landscape. Data for WASHCAP CPI-motif and V-1 are shown in blue and red, respectively.

**Supplementary Figure 15.**
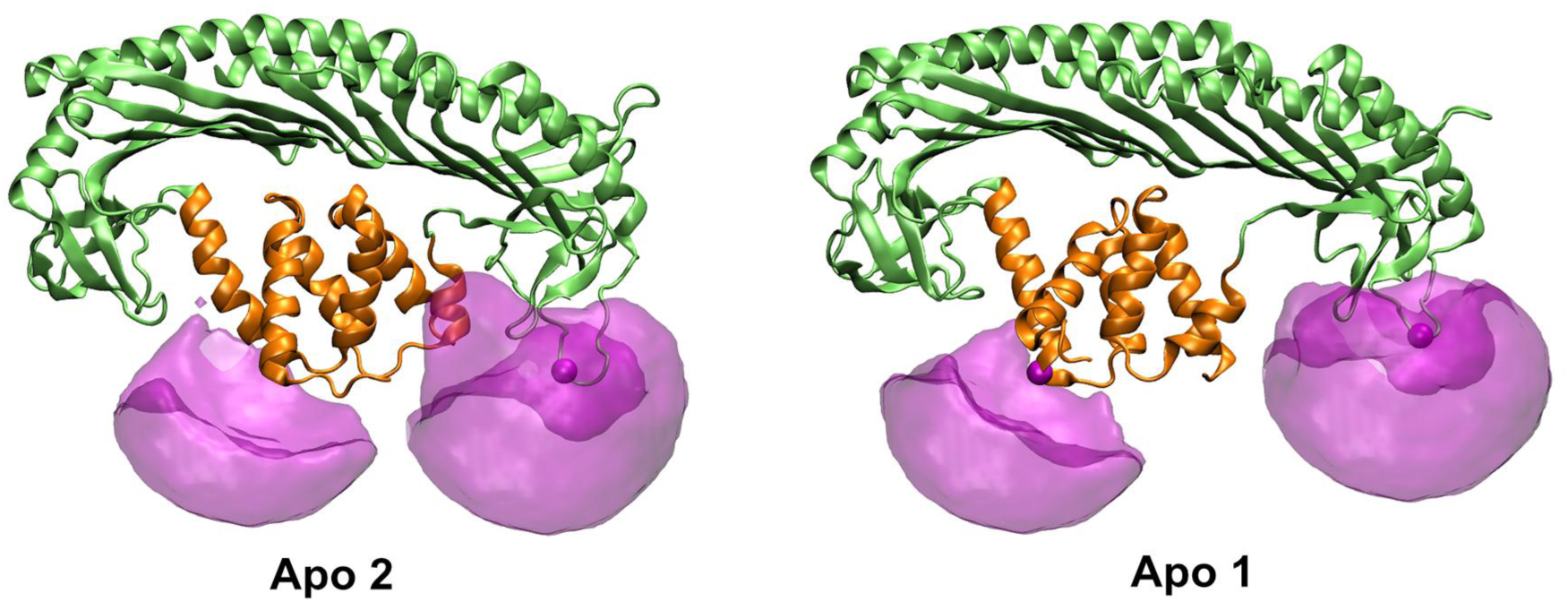
Illustration of the cloud of dye conformations for CP α44β84.

**Supplementary Figure 16.**
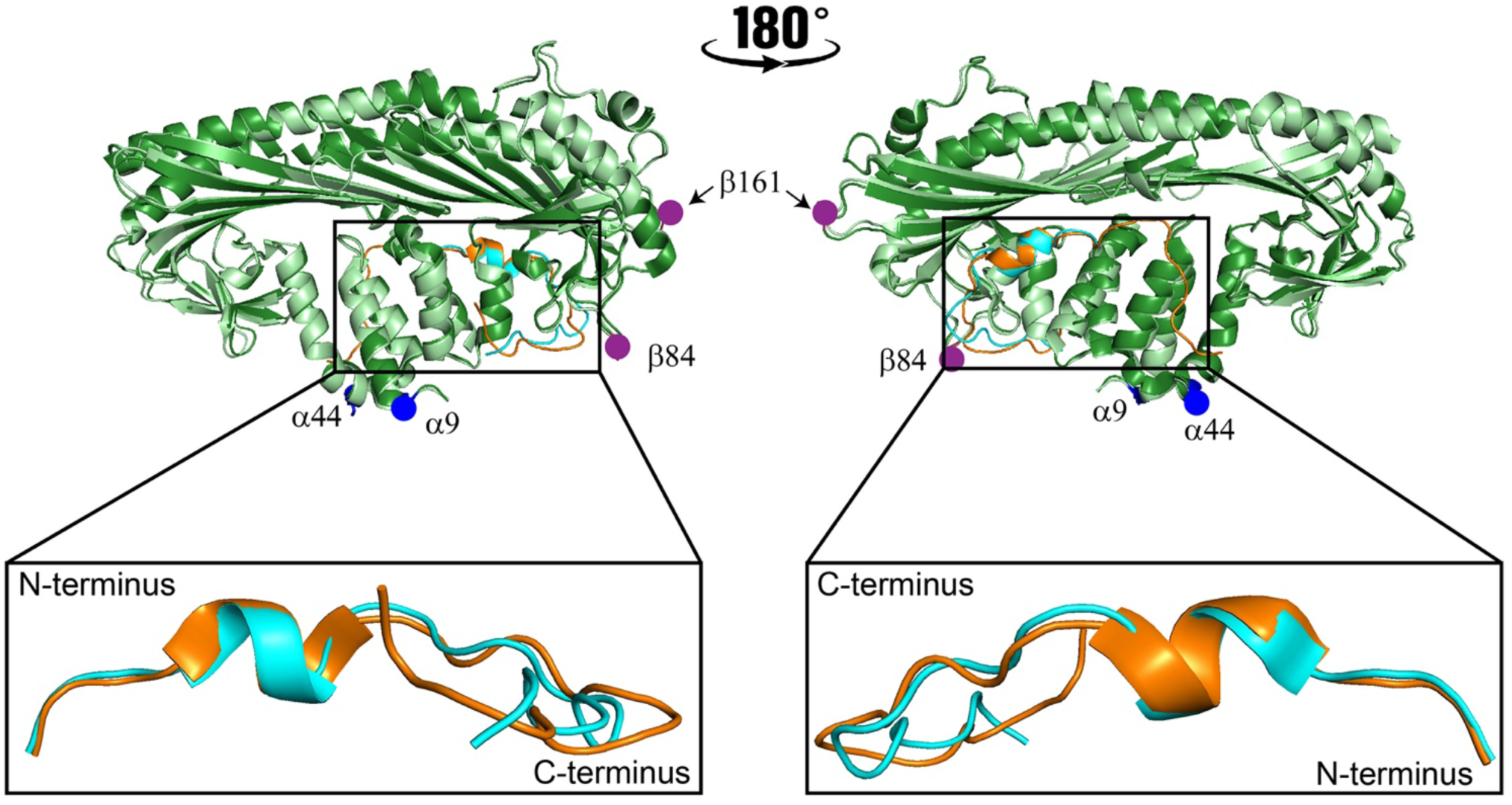
Alignment of crystal structures of the CP/CARMIL1 complex (PDB 3LK2) (light green and orange), and the CP/CKIP complex (PDB 3AA1) (dark green and cyan). Good alignment is observed for the backbone of the N-terminal region of the peptide, with divergence in the C-terminal half of the peptide. See Movie 4.

**Supplementary Figure 17.**
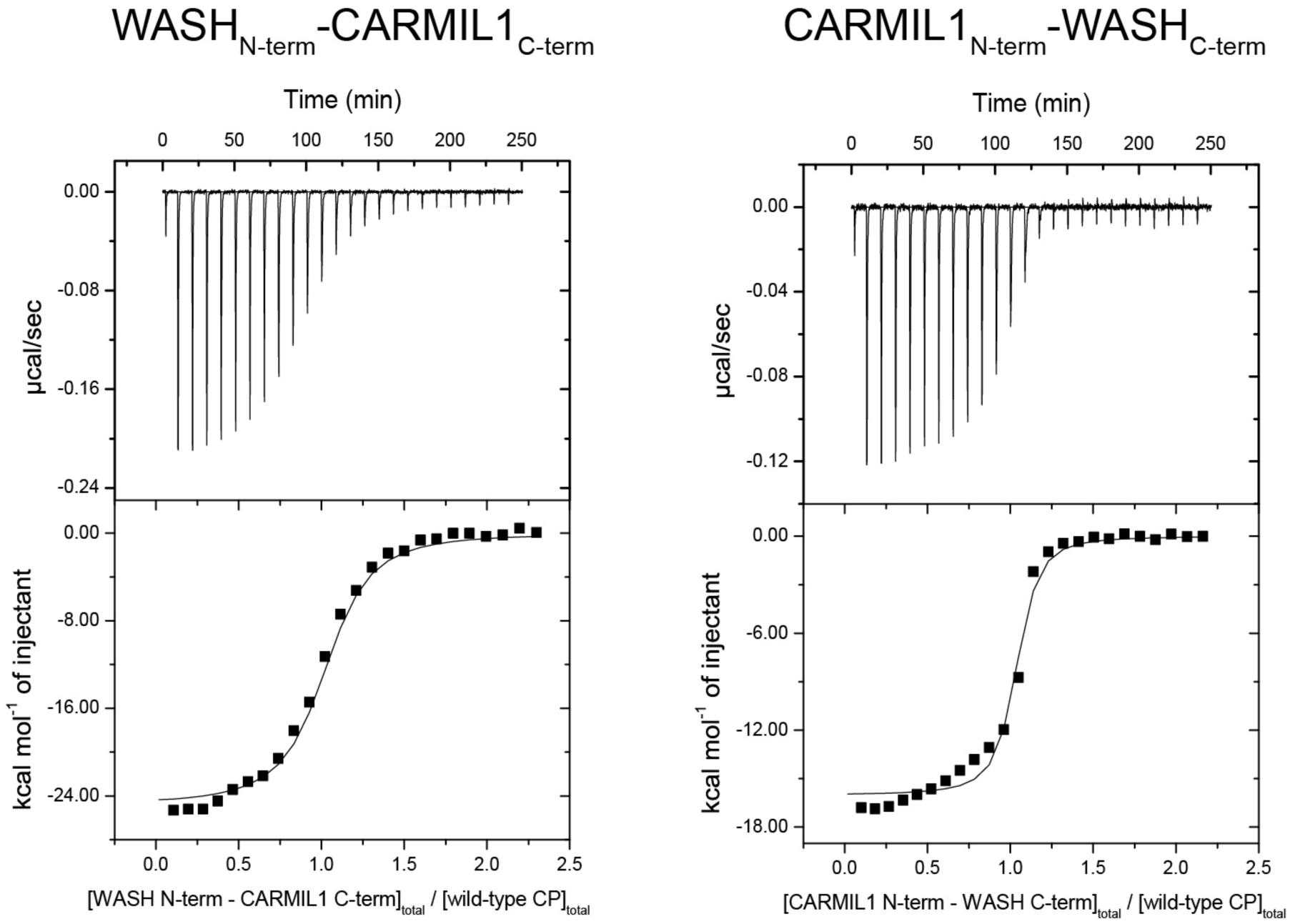
ITC data and fits for CP with WASH_N-term_-CARMIL1_C-term_ chimera and CARMIL1_N-term_-WASH_C-term_ chimera as function of the molar ratio of ligand (chimera peptide) and macromolecule (CP) concentration.

**Supplementary Figure 18.**
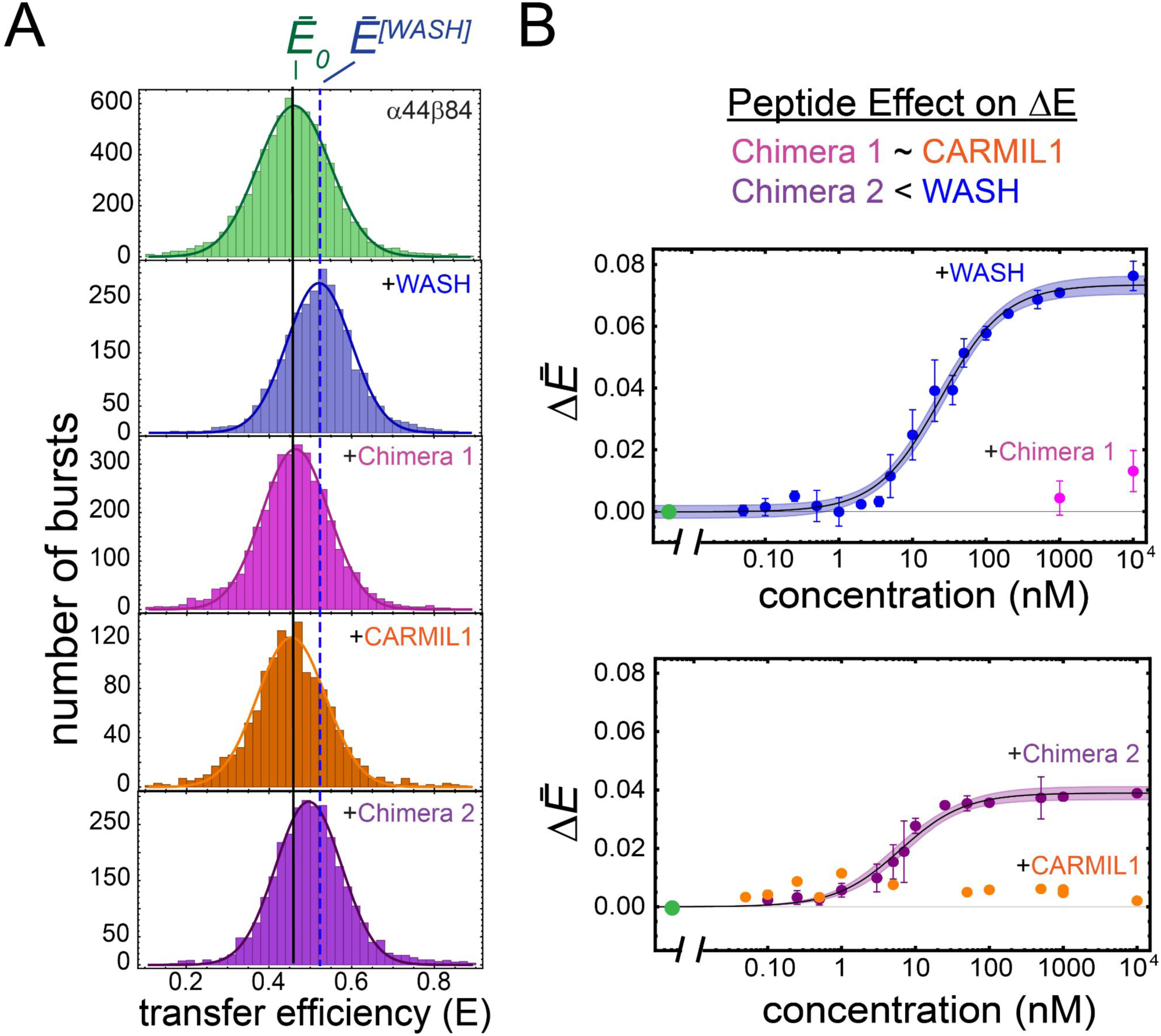
The C-terminus of the CPI motif is responsible for CP conformational changes in CP α44β84. A. Transfer efficiency histograms of CP α44β84 with WASHCAP (blue), chimera 1 WASH_N_-C1_C_ (fuchsia), CARMIL1 (orange), and chimera 2 C1_N_-WASH_C_ (purple). Experiments were performed in triplicate and one representative histogram is shown at saturation of ligand, 1 μM. B. Titration curves for all experiments are shown with the average change in transfer efficiency plotted versus concentration of peptide. The shading represents the 95% confidence interval of the fit.

**Supplementary Figure 19.**
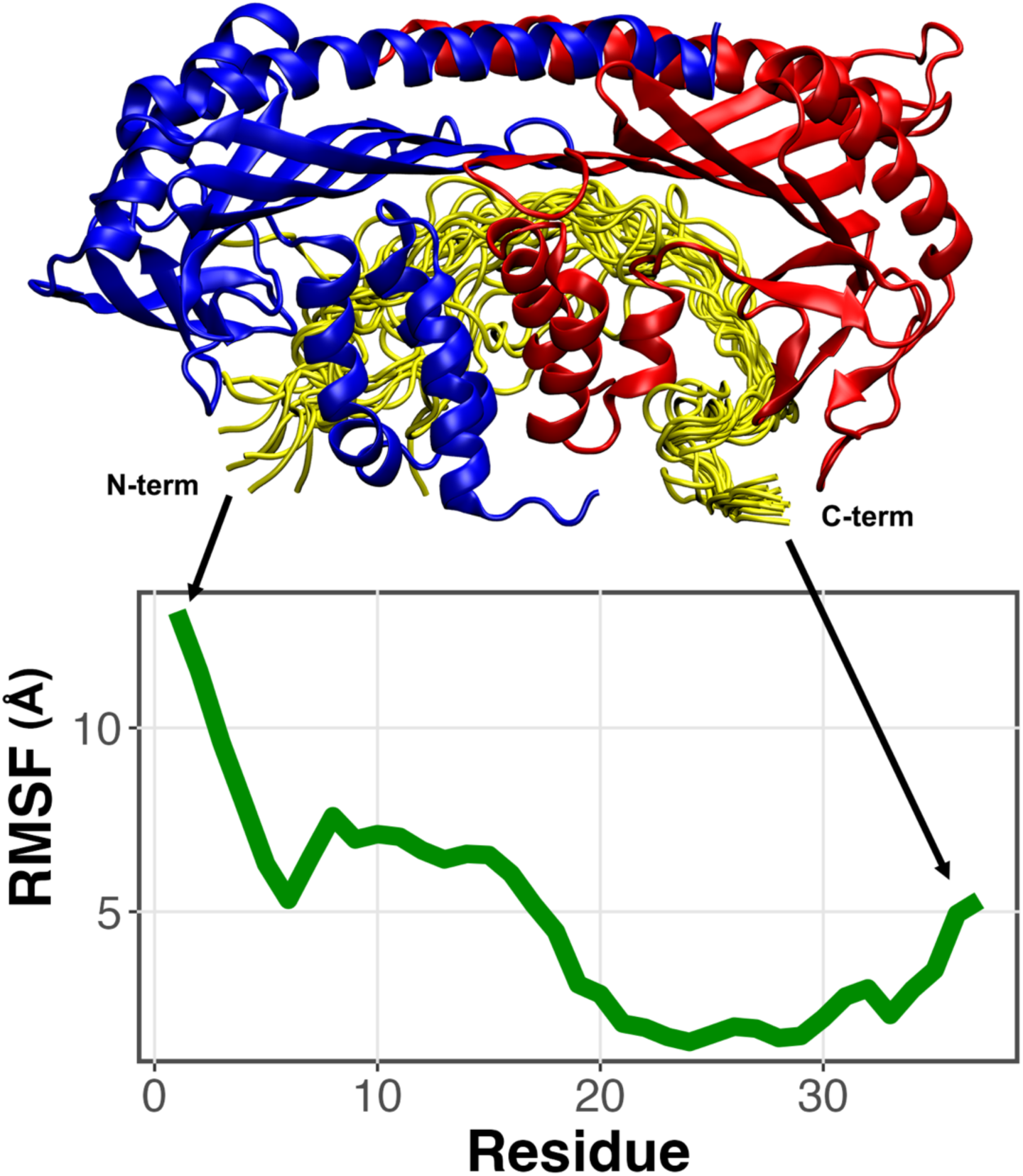
RMSF of the bound CPI motif from MD simulations. The C-terminal portion of the CPI motif shows lower RMSF values than does the N-terminal portion. These MD results are consistent with the chimera biochemical results showing that the C-terminal portion largely confers the function of the CPI motif.

**Supplementary Figure 20.**
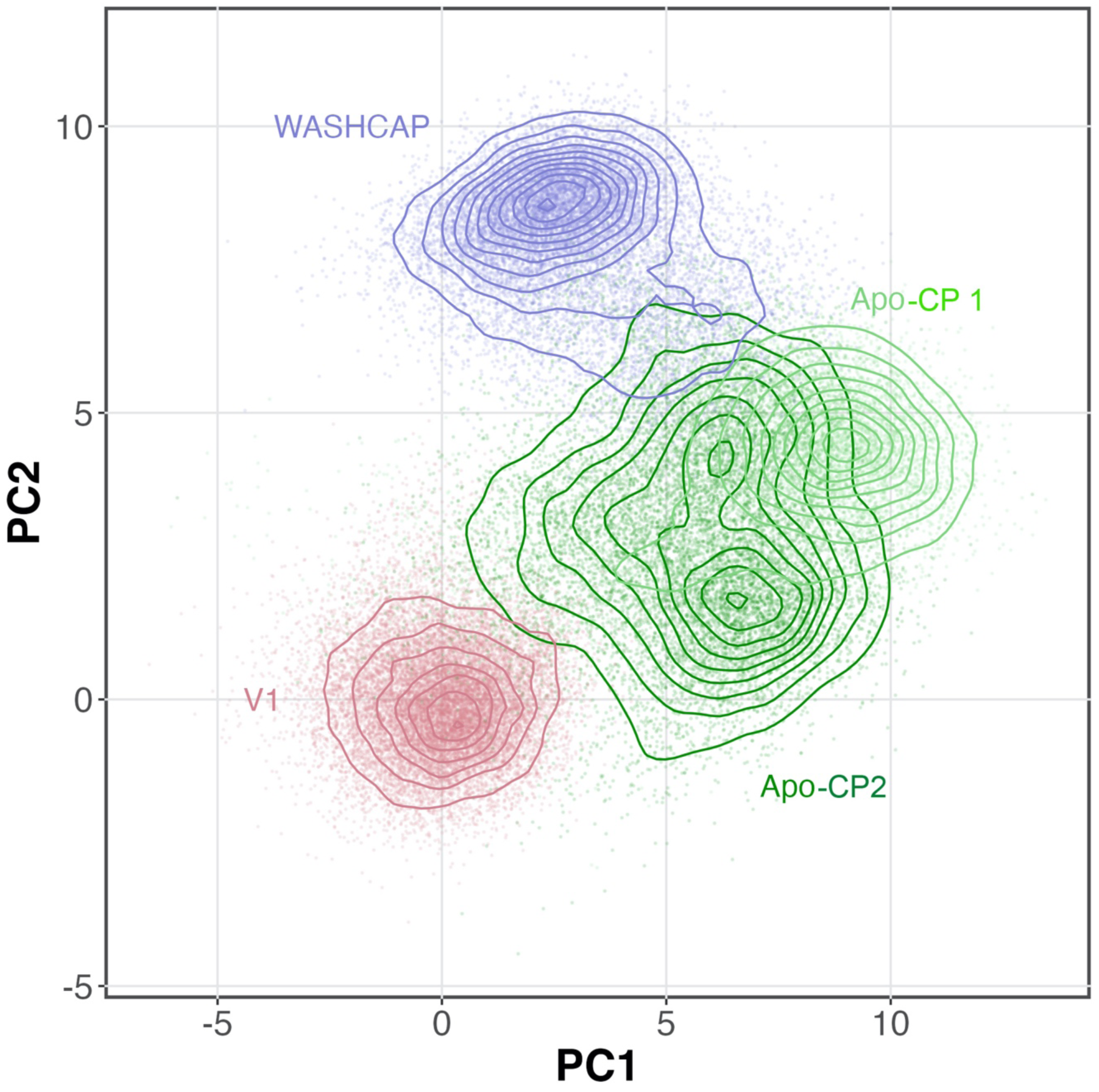
Principle component analysis of the key residues that make up the V-1 binding site on CP. The V-1-CP simulation (red) represents the bound structure. Apo-CP 2 (dark green) is closest in conformation to V-1-CP, while the Apo-CP 1 (light green) and WASHCAP-CP (blue) structures deviate more. These results suggest that V-1 would have a stronger binding affinity for the Apo-CP 2 state and a weaker affinity for the Apo-CP 1 and WASHCAP-CP states.

## Supplementary Movies

**Supplementary Movie 1.** Molecular dynamic (MD) simulation demonstrating the transition between Apo-CP 1 and Apo-CP 2 (PDB File 1 and PDB File 2).

**Supplementary Movie 2.** Molecular dynamic (MD) simulation demonstrating the transition between Apo-CP 2 and WASHCAP-CP bound structures. (PDB File 2 and PDB File 3).

**Supplementary Movie 3.** Molecular dynamic (MD) simulation demonstrating the transition between Apo-CP 2 and V-1-CP bound structures. (PDB File 2 and PDB File 4).

**Supplementary Movie 4.** Rotation of the aligned crystal structures shown in Supplementary Figure 16.

**Supplementary Movie 5.** Molecular dynamic (MD) simulation demonstrating the transition between WASHCAP-CP and V-1-CP bound structures. (PDB File 3 and PDB File 4)

## Supplementary PDB Files

PDB files generated by MD simulations. Structures are shown in Supplementary Movies.

**PDB File 1.** CP-Apo 1.

**PDB File 2.** CP-Apo 2

**PDB File 3.** CP-WASHCAP

**PDB File 4.** CP-V-1.

---

**Supplementary Table 1.**
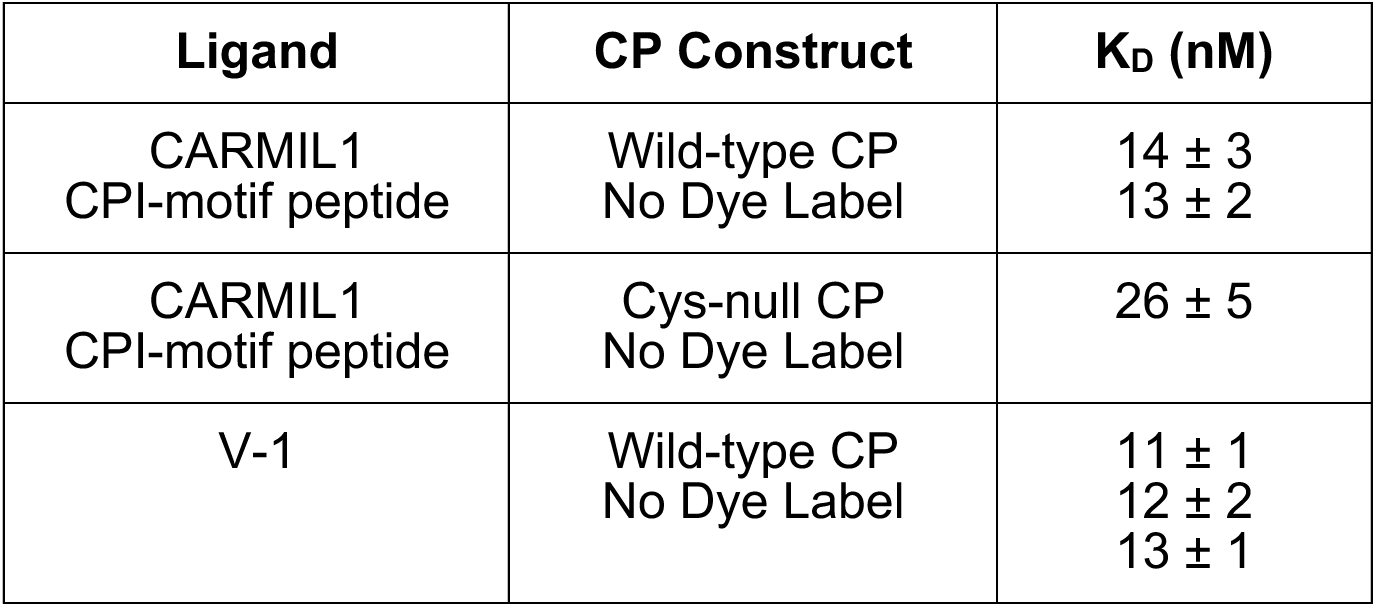
Ligand binding by ITC to CP constructs with no dye label. CARMIL1 CPI-motif peptide binding to unlabeled wild-type CP and Cys-null CP. V-1 binding to unlabeled wild-type CP. Each value represents one independent ITC experiment, with error of fitting.

**Supplementary Table 2.** Transfer Efficiencies for CP α44β84 and CP α9β161 with V-1 and CPI-motif peptides. ΔĒ is the difference between the transfer efficiencies of the Apo state and the corresponding bound state. For α9β161, ΔĒ is reported with respect to the two Apo states for V-1 binding. For CPI-motif peptides, the value is reported only with respect to Ē_2_, since no apparent change occurs for Ē_1._ Reported values are mean and standard deviation of n independent repeats on different days.

| Construct | Ligand | n | | Ligand<br>Concentration ($\mu\text{M}$) | $\Delta \bar{E}$ |
| --- | --- | --- | --- | --- | --- |
| $\alpha 44\beta 84$ | V-1 | 4 | $\bar{E}^{[V-1]}$ | 10 | $-0.016 \pm 0.004$ |
| | WASHCAP | 3 | $\bar{E}^{[WASH]}$ | 1 | $+0.071 \pm 0.001$ |
| | | | | 10 | $+0.076 \pm 0.005$ |
| | CKIP | 3 | $\bar{E}^{[CKIP]}$ | 1 | $+0.031 \pm 0.002$ |
| | CARMIL1 | 4 | $\bar{E}^{[CARMIL1]}$ | 1 | $+0.002 \pm 0.003$ |
| | WASH<sub>N</sub>-C1<sub>C</sub> | 3 | $\bar{E}^{[WN-C1C]}$ | - | $+0.014 \pm 0.006$ |
| | C1<sub>N</sub>-WASH<sub>C</sub> | 4 | $\bar{E}^{[C1N-WC]}$ | - | $+0.039 \pm 0.001$ |
| $\alpha 9\beta 161$ | V-1 | 3 | $\bar{E}_1^{[V-1]}$ | 10 | $-0.056 \pm 0.012$ |
| | | | $\bar{E}_2^{[V-1]}$ | 10 | $-0.033 \pm 0.003$ |
| | WASHCAP | 4 | $\bar{E}_2^{[WASH]}$ | 1 | $+0.112 \pm 0.006$ |
| | | | | 10 | $+0.112 \pm 0.007$ |
| | CKIP | 3 | $\bar{E}_2^{[CKIP]}$ | 1 | $+0.068 \pm 0.008$ |
| | CARMIL1 | 3 | $\bar{E}_2^{[CARMIL1]}$ | 1 | $+0.042 \pm 0.002$ |
| | WASH<sub>N</sub>-C1<sub>C</sub> | 4 | $\bar{E}_2^{[WN-C1C]}$ | - | $+0.047 \pm 0.004$ |
| | C1<sub>N</sub>-WASH<sub>C</sub> | 4 | $\bar{E}_2^{[C1N-WC]}$ | - | $+0.114 \pm 0.006$ |

**Supplementary Table 3.** Analysis of subpopulation-specific time-resolved fluorescence decays. Mean and standard deviation values are reported in units of nanoseconds; n is the number of independent experiments.

| sample | n | $\tau_{\text{Donly}}$ | $\tau_{\text{rot-Donly}}$ | $\tau_{\text{DA}}$ | $\tau_{\text{rot-DA}}$ | $\tau_{\text{FRET}}$ | $\tau_{\text{Aonly}}$ | $\tau_{\text{rot-Aonly}}$ |
| --- | --- | --- | --- | --- | --- | --- | --- | --- |
| Apo-CP | 14 | 3.87±0.05 | 0.34±0.07 | 2.50±0.08 | 0.47±0.06 | 7.1±0.4 | 4.06±0.12 | 0.30±0.06 |
| WASHCAP | 3 | 3.81±0.12 | 0.34±0.04 | 2.27±0.06 | 0.50±0.06 | 5.6±0.6 | 4.08±0.04 | 0.42±0.02 |
| CKIP | 3 | 3.71±0.07 | 0.36±0.02 | 2.41±0.07 | 0.57±0.07 | 6.9±0.6 | 4.09±0.03 | 0.35±0.01 |
| CARMIL1 | 4 | 3.91±0.07 | 0.37±0.03 | 2.50±0.04 | 0.51±0.05 | 6.9±0.5 | 4.09±0.03 | 0.46±0.03 |
| V-1 | 3 | 3.86±0.08 | 0.43±0.09 | 2.51±0.04 | 0.47±0.03 | 7.2±0.2 | 4.08±0.01 | 0.22±0.02 |
| WN-C1C | 3 | 3.80±0.03 | 0.40±0.03 | 2.43±0.03 | 0.47±0.01 | 6.8±0.3 | 4.01±0.07 | 0.79±0.41 |
| C1N-WC | 3 | 3.78±0.07 | 0.40±0.05 | 2.32±0.02 | 0.46±0.02 | 6.0±0.1 | 4.02±0.10 | 0.87±0.51 |

FRET probe α1S9C β2S161C
| sample | n | E | $\tau_{\text{Donly}}$ | $\tau_{\text{rot-Donly}}$ | $\tau_{\text{DA}}$ | $\tau_{\text{rot-DA}}$ | $\tau_{\text{FRET}}$ | $\tau_{\text{Aonly}}$ | $\tau_{\text{rot-Aonly}}$ |
| --- | --- | --- | --- | --- | --- | --- | --- | --- | --- |
| Apo-CP | 13 | E1 | 3.84±0.04 | 0.48±0.05 | 3.35±0.02 | 0.59±0.06 | 26.3±1.7 | 4.15±0.02 | 0.58±0.05 |
| | | E2 | 3.84±0.04 | 0.48±0.05 | 2.80±0.04 | 0.60±0.05 | 10.3±0.7 | 4.15±0.02 | 0.58±0.05 |
| WASHCAP | 2 | E1 | 3.87±0.01 | 0.40±0.08 | 3.40±0.10 | 0.45±0.13 | 28.5±6.4 | 4.07±0.08 | 0.53±0.08 |
| | | E2 | 3.87±0.01 | 0.40±0.08 | 2.41±0.02 | 0.52±0.03 | 6.4±0.1 | 4.07±0.08 | 0.53±0.08 |
| CKIP | 1 | E1 | 3.80 | 0.39 | 3.34 | 0.52 | 28.0 | 4.10 | 0.58 |
|  |  | E2 | 3.80 | 0.39 | 2.56 | 0.60 | 7.9 | 4.10 | 0.58 |
| CARMIL1 | 1 | E1 | 3.88 | 0.44 | 3.32 | 0.81 | 22.9 | 4.06 | 0.52 |
|  |  | E2 | 3.88 | 0.44 | 2.55 | 0.53 | 7.5 | 4.06 | 0.52 |
| V-1 | 3 | E1 | 3.88±0.02 | 0.44±0.05 | 3.37±0.03 | 0.61±0.01 | 25.8±1.0 | 4.12±0.01 | 0.58±0.05 |
| | | E2 | 3.88±0.02 | 0.44±0.05 | 2.84±0.01 | 0.63±0.02 | 10.6±0.2 | 4.12±0.01 | 0.58±0.05 |
| WN-C1C | 4 | E1 | 3.86±0.04 | 0.52±0.05 | 3.34±0.05 | 0.58±0.03 | 24.8±1.3 | 4.13±0.01 | 0.57±0.03 |
| | | E2 | 3.86±0.04 | 0.52±0.05 | 2.66±0.02 | 0.62±0.07 | 8.5±0.2 | 4.13±0.01 | 0.57±0.03 |
| C1N-WC | 4 | E1 | 3.82±0.03 | 0.49±0.02 | 3.35±0.03 | 0.60±0.07 | 27.8±1.5 | 4.12±0.04 | 0.51±0.05 |
| | | E2 | 3.82±0.03 | 0.49±0.02 | 2.46±0.07 | 0.61±0.05 | 6.95±0.5 | 4.12±0.04 | 0.51±0.05 |

**Supplementary Table 4.**
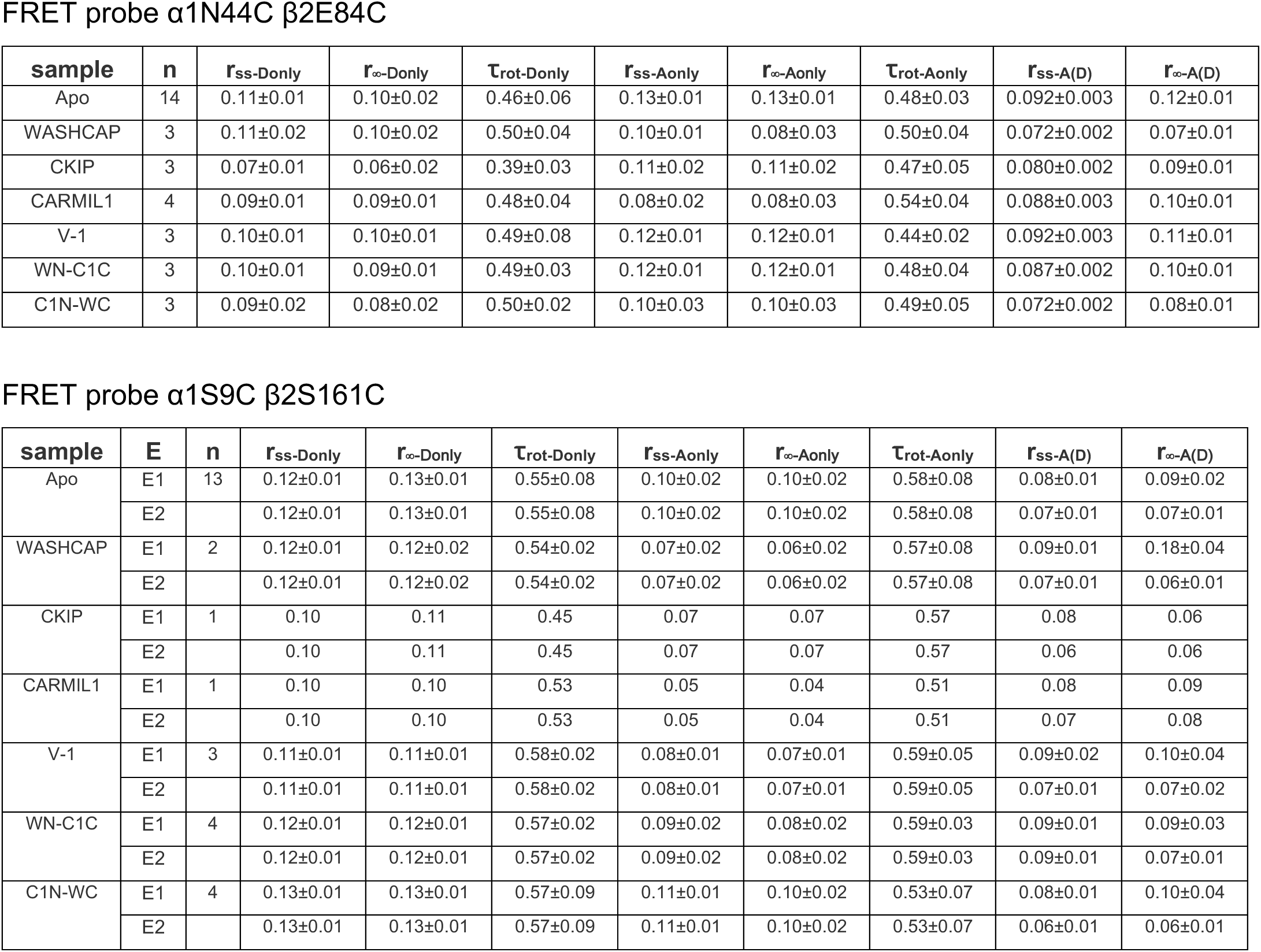
Steady-state and time resolved anisotropies in aqueous buffer conditions. Mean and standard deviation values are reported in units of nanoseconds; n is the number of independent experiments.

**Supplementary Table 5.** Effective dissociation constants estimated from single-molecule FRET and ITC analyses for binding of CARMIL1, CKIP, WASHCAP, and V-1 to CP. For single titration curves the best fit parameter and the error of the fit are reported. From independent titrations, collected on different days, mean and standard deviation (s.d.) are reported. Transfer efficiencies fitted to Eq. S6a. Effective K*_D_ from 3-state and 4-state models obtained from Eq. S9 and S12.

|  | Single-molecule FRET |  |  |  |  |  | ITC |  |
| --- | --- | --- | --- | --- | --- | --- | --- | --- |
|  | Transfer efficiency |  | 3-state model |  | 4-state model |  |  |  |
| FRET labels | $K_D$ (nM)<br>Individual titrations | $K_D$ (nM)<br>(mean ± s.d.) | $K_D^*$ (nM)<br>Individual titrations | $K_D^*$ (nM)<br>(mean ± s.d.) | $K_D^*$ (nM)<br>Individual titrations | $K_D^*$ (nM)<br>(mean ± s.d.) | $K_D$ (nM)<br>Individual titrations | $K_D$ (nM)<br>(mean ± s.d.) |
| **CARMIL1** |  |  |  |  |  |  |  |  |
| $\alpha 9\beta 161$ | 13 ± 4<br>21 ± 7<br>17 ± 6 | 17 ± 4 | 16 ± 2<br>21 ± 3<br>13 ± 2 | 17 ± 4 | 15 ± 1<br>19 ± 2<br>12 ± 1 | 15 ± 3 | 14 ± 1<br>24 ± 4 | 19 ± 7 |
| $\alpha 44\beta 84$ | n/a | n/a | n/a | n/a | n/a | n/a | 65 ± 9<br>66 ± 10 | 66 ± 1 |
| **CKIP** |  |  |  |  |  |  |  |  |
| $\alpha 9\beta 161$ | 7.4 ± 1.4<br>4 ± 2<br>9 ± 3 | 7 ± 3 | 9 ± 1<br>6.6 ± 0.8<br>11 ± 1 | 9 ± 2 | 10 ± 1<br>8.0 ± 0.8<br>5.8 ± 0.4 | 8 ± 1 | 21 ± 3<br>26 ± 3 | 24 ± 4 |
| $\alpha 44\beta 84$ | 10 ± 2<br>17 ± 4<br>11 ± 4 | 13 ± 4 | n/a | n/a | n/a | n/a | 29 ± 3<br>38 ± 4 | 34 ± 6 |
| **WASHCAP** |  |  |  |  |  |  |  |  |
| $\alpha 9\beta 161$ | 0.37 ± 0.04<br>0.7 ± 0.2<br>1.5 ± 0.3<br>3.1 ± 0.6 | 1.4 ± 1.2 | 0.54 ± 0.06<br>0.8 ± 0.1<br>1.5 ± 0.1<br>4.4 ± 0.4 | 2 ± 2 | 9 ± 2<br>1.4 ± 0.1<br>0.66 ± 0.05<br>0.50 ± 0.04 | 3 ± 5 | 6 ± 1<br>5 ± 1 | 5.5 ± 0.7 |
| $\alpha 44\beta 84$ | 19 ± 3<br>20 ± 4<br>28 ± 3 | 22 ± 5 | n/a | n/a | n/a | n/a | 13 ± 3<br>20 ± 3 | 17 ± 5 |
| **V-1** |  |  |  |  |  |  |  |  |
| $\alpha 9\beta 161$ | 8 ± 3<br>8 ± 3<br>28 ± 9 | 15 ± 12 | 540 ± 130<br>130 ± 30<br>510 ± 130 | 400±200 | 22 ± 4<br>23 ± 4<br>25 ± 4 | 23 ± 2 | 52 ± 9<br>53 ± 8<br>67 ± 13 | 57 ± 8 |
| $\alpha 44\beta 84$ | 11 ± 4<br>22 ± 10<br>9 ± 4 | 14 ± 7 | n/a | n/a | n/a | n/a | 18 ± 4<br>20 ± 5 | 19 ± 1 |

**Supplementary Table 6.**
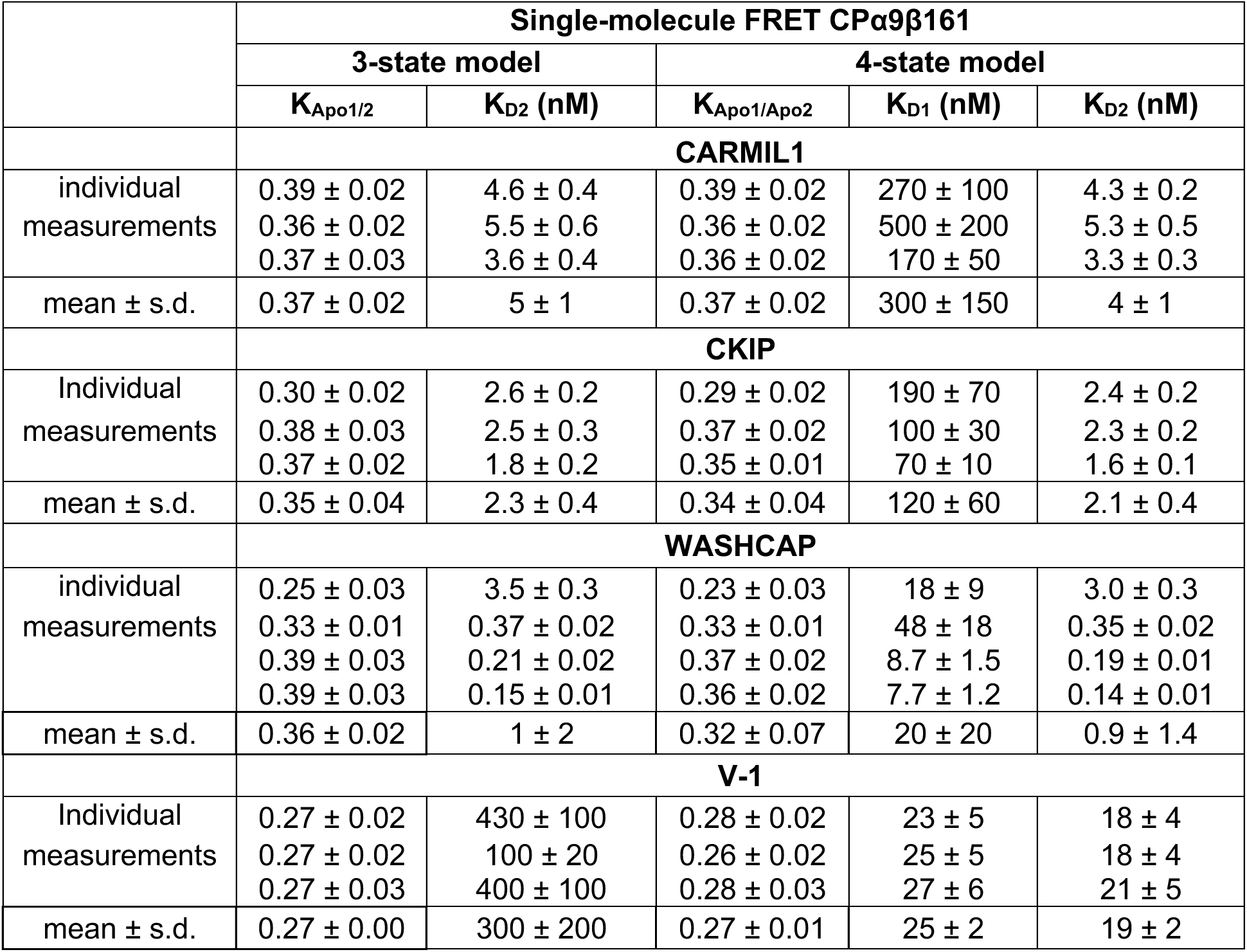
3-state and 4-state model analysis of single-molecule FRET data for CPα9β161 binding to CARMIL1, CKIP, WASHCAP, and V-1. For single titration curves, the best fit parameter and the error of the fit are reported. From independent titrations, collected on different days, mean and standard deviation (s.d.) are reported. 3-state model results were fitted to Eq. and 4-state model results are fitted to Eq. S7 and Eq. S10.

**Supplementary Table 7.**
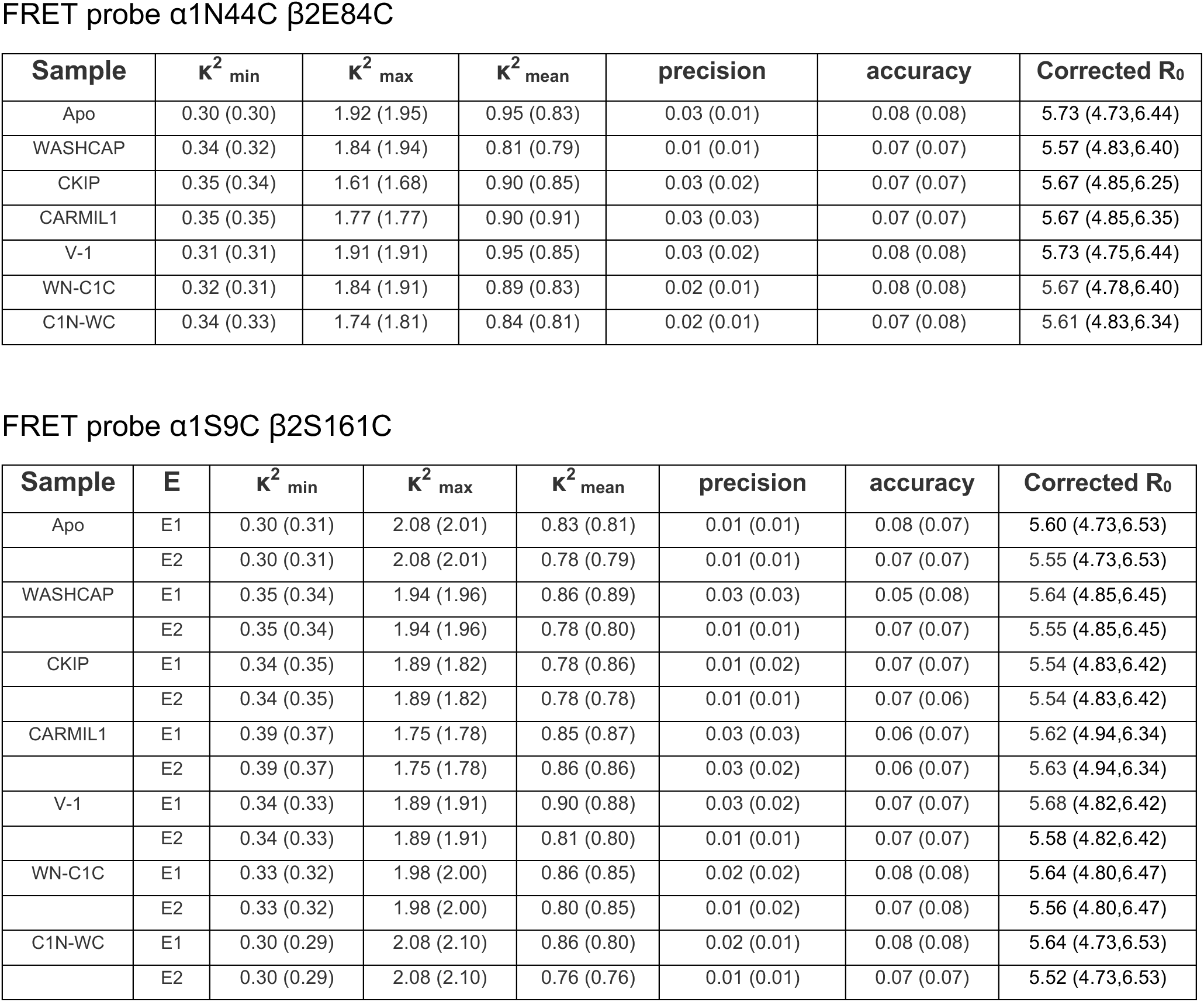
Estimates of deviations in the orientational κ^2^ factor in aqueous buffer conditions. Values in parentheses are estimated from steady-state anisotropies.

FRET probe $\alpha 1N44C$ $\beta 2E84C$
| Sample | $\kappa^2_{\min}$ | $\kappa^2_{\max}$ | $\kappa^2_{\text{mean}}$ | precision | accuracy | Corrected $R_0$ |
| --- | --- | --- | --- | --- | --- | --- |
| Apo | 0.30 (0.30) | 1.92 (1.95) | 0.95 (0.83) | 0.03 (0.01) | 0.08 (0.08) | 5.73 (4.73,6.44) |
| WASHCAP | 0.34 (0.32) | 1.84 (1.94) | 0.81 (0.79) | 0.01 (0.01) | 0.07 (0.07) | 5.57 (4.83,6.40) |
| CKIP | 0.35 (0.34) | 1.61 (1.68) | 0.90 (0.85) | 0.03 (0.02) | 0.07 (0.07) | 5.67 (4.85,6.25) |
| CARMIL1 | 0.35 (0.35) | 1.77 (1.77) | 0.90 (0.91) | 0.03 (0.03) | 0.07 (0.07) | 5.67 (4.85,6.35) |
| V-1 | 0.31 (0.31) | 1.91 (1.91) | 0.95 (0.85) | 0.03 (0.02) | 0.08 (0.08) | 5.73 (4.75,6.44) |
| WN-C1C | 0.32 (0.31) | 1.84 (1.91) | 0.89 (0.83) | 0.02 (0.01) | 0.08 (0.08) | 5.67 (4.78,6.40) |
| C1N-WC | 0.34 (0.33) | 1.74 (1.81) | 0.84 (0.81) | 0.02 (0.01) | 0.07 (0.08) | 5.61 (4.83,6.34) |

FRET probe $\alpha 1S9C$ $\beta 2S161C$
| Sample | E | $\kappa^2_{\min}$ | $\kappa^2_{\max}$ | $\kappa^2_{\text{mean}}$ | precision | accuracy | Corrected $R_0$ |
| --- | --- | --- | --- | --- | --- | --- | --- |
| Apo | E1 | 0.30 (0.31) | 2.08 (2.01) | 0.83 (0.81) | 0.01 (0.01) | 0.08 (0.07) | 5.60 (4.73,6.53) |
|  | E2 | 0.30 (0.31) | 2.08 (2.01) | 0.78 (0.79) | 0.01 (0.01) | 0.07 (0.07) | 5.55 (4.73,6.53) |
| WASHCAP | E1 | 0.35 (0.34) | 1.94 (1.96) | 0.86 (0.89) | 0.03 (0.03) | 0.05 (0.08) | 5.64 (4.85,6.45) |
|  | E2 | 0.35 (0.34) | 1.94 (1.96) | 0.78 (0.80) | 0.01 (0.01) | 0.07 (0.07) | 5.55 (4.85,6.45) |
| CKIP | E1 | 0.34 (0.35) | 1.89 (1.82) | 0.78 (0.86) | 0.01 (0.02) | 0.07 (0.07) | 5.54 (4.83,6.42) |
|  | E2 | 0.34 (0.35) | 1.89 (1.82) | 0.78 (0.78) | 0.01 (0.01) | 0.07 (0.06) | 5.54 (4.83,6.42) |
| CARMIL1 | E1 | 0.39 (0.37) | 1.75 (1.78) | 0.85 (0.87) | 0.03 (0.03) | 0.06 (0.07) | 5.62 (4.94,6.34) |
|  | E2 | 0.39 (0.37) | 1.75 (1.78) | 0.86 (0.86) | 0.03 (0.02) | 0.06 (0.07) | 5.63 (4.94,6.34) |
| V-1 | E1 | 0.34 (0.33) | 1.89 (1.91) | 0.90 (0.88) | 0.03 (0.02) | 0.07 (0.07) | 5.68 (4.82,6.42) |
|  | E2 | 0.34 (0.33) | 1.89 (1.91) | 0.81 (0.80) | 0.01 (0.01) | 0.07 (0.07) | 5.58 (4.82,6.42) |
| WN-C1C | E1 | 0.33 (0.32) | 1.98 (2.00) | 0.86 (0.85) | 0.02 (0.02) | 0.08 (0.08) | 5.64 (4.80,6.47) |
|  | E2 | 0.33 (0.32) | 1.98 (2.00) | 0.80 (0.85) | 0.01 (0.02) | 0.07 (0.08) | 5.56 (4.80,6.47) |
| C1N-WC | E1 | 0.30 (0.29) | 2.08 (2.10) | 0.86 (0.80) | 0.02 (0.01) | 0.08 (0.08) | 5.64 (4.73,6.53) |
|  | E2 | 0.30 (0.29) | 2.08 (2.10) | 0.76 (0.76) | 0.01 (0.01) | 0.07 (0.07) | 5.52 (4.73,6.53) |

**Supplementary Table 8.** K_D_ values for chimeric peptides obtained from the change in transfer efficiency (single-molecule FRET) compared to ITC.

| CPI chimera | CP FRET Pair | Single-molecule FRET $K_D$ (nM) | ITC $K_D$ (nM) with wt-CP |
| --- | --- | --- | --- |
| C1<sub>N</sub>-WASH<sub>C</sub> | α9β161 | 0.19 ± 0.02<br>0.3 ± 0.1 | 6 ± 2<br>9 ± 2 |
| | α44β84 | 10 ± 3<br>4 ± 1<br>5 ± 2 | |
| WASH<sub>N</sub>-C1<sub>C</sub> | α9β161 | 180 ± 20<br>210 ± 30<br>130 ± 17 | 51 ± 6<br>38 ± 5 |
| | α44β84 | No FRET change | |

## Notes

### Competing Interest Statement

The authors have declared no competing interest.

